# Cohesin prevents local mixing of condensed euchromatic domains in living human cells

**DOI:** 10.1101/2025.08.27.672592

**Authors:** Masa A. Shimazoe, Shiori Iida, Katsuhiko Minami, Koichi Higashi, Sachiko Tamura, Yoshiaki Kobayashi, Shin Fujishiro, Le Xiong, Kako Nakazato, S. S. Ashwin, Tomoko Nishiyama, Yu Nagata, Masato T. Kanemaki, Akane Kawaguchi, Yasuyuki Ohkawa, Lothar Schermelleh, Atsushi Toyoda, Liangqi Xie, Ken Kurokawa, Hiroshi Ochiai, Masaki Sasai, Kazuhiro Maeshima

## Abstract

The human genome is folded into chromatin loops by the cohesin complex, forming functional chromatin domains that underlie transcription and DNA replication/repair. However, how cohesin organizes these domains in living cells, especially in active euchromatin, remains elusive. To address this question, we combined single-nucleosome imaging/tracking and super-resolution 3D-structured illumination microscopy (3D-SIM) with euchromatin-specific labeling of histone H3.3. Using this nanoscopic approach, we revealed that euchromatin forms condensed domains that are constrained by cohesin-mediated loops. This organization refines the classical textbook view of euchromatin as largely open, in line with emerging evidence. Transcription machinery appears to be located near the condensed domain surfaces/borders. Cohesin loss increased nucleosome fluidity within these domains without altering their overall compaction, leading to local mixing of domains and compromising transcriptional insulation. These findings uncover an unexpected physical role of cohesin in maintaining the integrity of condensed euchromatic domains and ensuring proper higher-order regulation of gene expression.

## Introduction

In human cells, two meters of genomic DNA are wrapped around core histones to form nucleosomes ^1,2^. Various imaging studies have long suggested that a string of nucleosomes is folded irregularly into functional chromatin domains in the interphase nucleus ^3–10^. Genomics analyses, such as Hi-C ^11^, have also identified similar domains or topologically associating domains, TADs, with distinct epigenetic marks ^12–15^. The textbook view of genomic chromatin describes that transcriptionally inactive heterochromatin forms condensed domains, whereas active euchromatin forms open ones, thereby supposedly making euchromatin accessible to transcription machinery ^16,17^. However, the expression profile of euchromatic genes varies greatly from cell to cell ^18^ ^19^, implying that transcriptional regulation may be governed by higher-order chromatin organization rather than a simple “open-or-closed” binary model. Consistent with this, recent evidence challenges this established textbook view, suggesting that euchromatin is also organized into largely condensed domains, which may contribute to higher-order gene regulation ^7,8,10,20–22^.

While interactions between genomic sequences within and between domains are well recognized as crucial for transcriptional regulation ^13,22–24^, DNA replication ^25^, and repair ^26^, the physical properties of chromatin domains have been comparatively overlooked. Yet, they are equally important. Chromatin dynamics and their physical properties directly influence genomic accessibility ^27,28^ and can modulate the efficiency and probability of these biochemical reactions. Therefore, elucidating how the physical properties of chromatin domains are regulated is crucial.

A key factor influencing both chromatin organization and dynamics is the cohesin complex, composed of four subunits (SMC1, SMC3, RAD21, and STAG1/STAG2) ^29–33^. Cohesin has two main roles in organizing genomic chromatin. One is to hold replicated sister chromatids together for their proper segregation during cell division (“cohesion”) and the other is to form chromatin loops within individual chromosomes to create the domains described above ^34–36^. Cohesin may also regulate the physical properties of chromatin in living cells. Indeed, pioneering work using the *lacO*/EGFP-LacI system in budding yeast demonstrated that cohesin loss increases chromatin motion, presumably via the loss of sister chromatid cohesion ^37,38^. Subsequent studies revealed that in higher eukaryotic cells, cohesin depletion also increases chromatin motion ^5,19,39–41^. However, it is still unclear how cohesin constrains chromatin, whether primarily through sister chromatid cohesion or loop formation, and which chromatin domains, euchromatin or heterochromatin, are most constrained. Furthermore, how cohesin-mediated constraints affect the physical properties of chromatin domains and their genome functions such as transcriptional regulation remains poorly understood.

To address these questions, we developed a combined approach of single-nucleosome imaging ^5,27,42,43^ and 3D-structured illumination microscopy (3D-SIM) ^7,44,45^ ^46^, incorporating a specific labeling strategy with the histone variant H3.3, which is enriched in active chromatin regions ^47–49^. Using these advanced nanoscopic imaging techniques, we uncovered condensed euchromatic domains in living human cells, providing further critical evidence that challenges the established view of euchromatin as largely open. We also demonstrated that cohesin governs the fluidity of these euchromatic domains without affecting their overall compaction. Together with genomic data and computational modeling, these findings highlight a previously underappreciated role of cohesin in preventing local mixing of condensed euchromatin domains, and thus ensuring the effective insulation^50,51^ of transcription among them.

## Results

### Rapid depletion of cohesin increases local nucleosome motion in living human cells

To explore how cohesin organizes chromatin domains in living human cells, we combined single-nucleosome imaging/tracking ^5,27,42,43,52^ with rapid cohesin depletion using the auxin-inducible degron system ^53^. Single-nucleosome imaging/tracking can sensitively detect changes in the chromatin state in living cells. We used HCT116 cells expressing H2B-HaloTag (H2B-Halo) ^43^ and RAD21-mAID-mClover (mAC) ^53^ (Fig. S1a,b). H2B-Halo was labeled with tetramethylrhodamine (TMR) to visualize nucleosomes genome-wide, including those in both euchromatin and heterochromatin (Fig. S1c). Oblique illumination (HILO) microscopy ^54^ enabled imaging of individual nucleosomes as TMR-labeled dots of H2B-Halo (Fig. 1a–c), which were precisely tracked in 2D at 50 ms per frame for a total of 15–20 s, yielding ∼1400 trajectories per cell (Figs. 1b,d, S1d,e; see Methods for further details). Position determination accuracy is 12.5 nm (Fig. S1d). Tracking was specific to nucleosome-incorporated H2B-Halo, as the small free H2B-Halo pool (3.3%) diffused too fast to be observed (Movie S1, Fig. S1f). The mean squared displacement (MSD) analysis revealed sub-diffusive nucleosome motion with an anomalous exponent (α) of 0.43, suggesting constrained motion (black plots in Fig. 1e,g). Fixation nearly abolished this motion (gray plot, Fig. 1e,g).

**Fig. 1:**
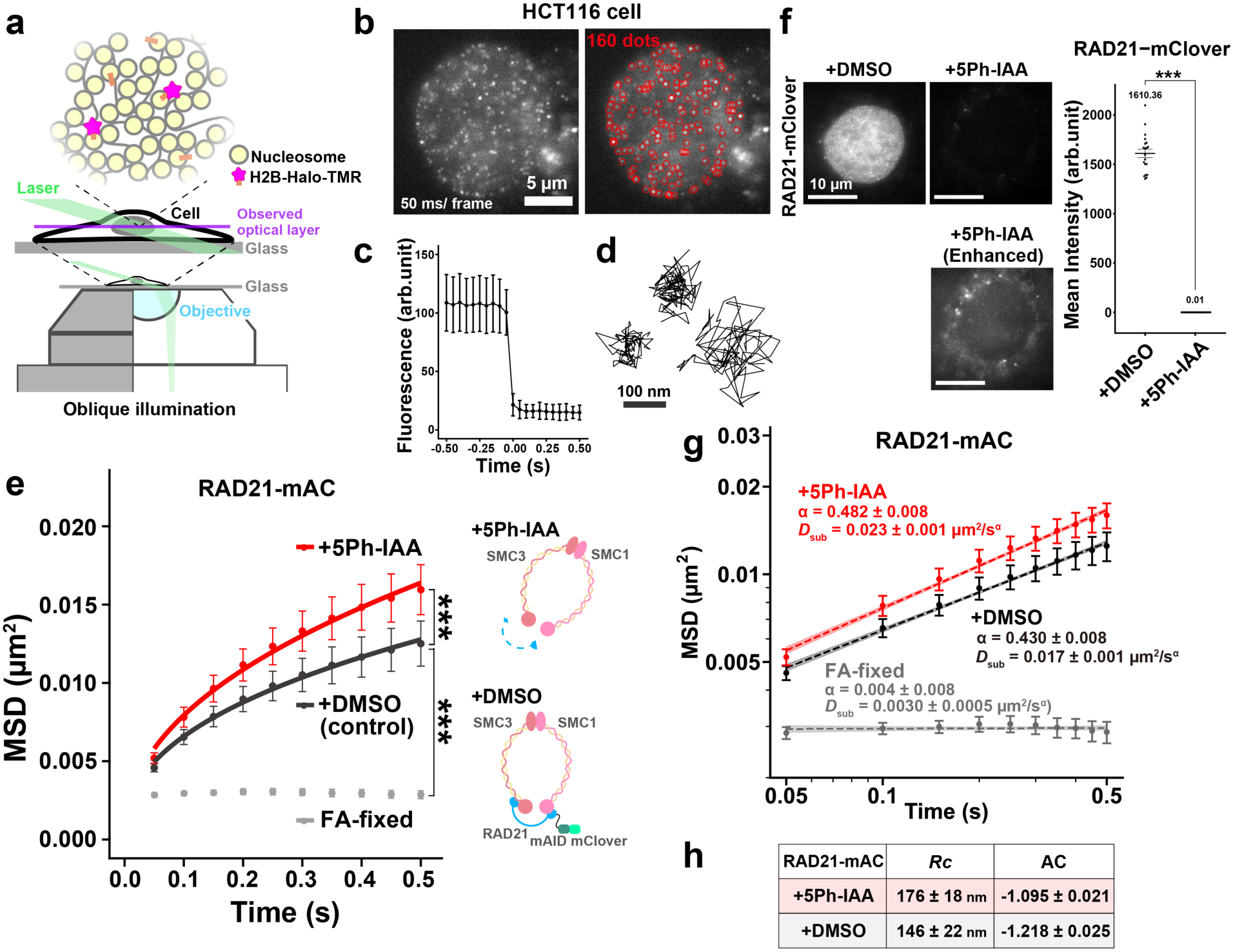
Single-nucleosome imaging revealed increased nucleosome motion upon cohesin depletion. **a**, Upper, a small fraction of H2B-Halo was fluorescently labeled with the TMR-HaloTag ligand (magenta star) and used to track nucleosome movements at super-resolution. Lower, oblique illumination microscopy. The illumination laser (green) excites fluorescent molecules within a thin optical layer (purple) of the nucleus, reducing background noise. **b**, Single-nucleosome (H2B-Halo-TMR) image of a living HCT116 nucleus after background subtraction. Red circles highlight the 160 tracked dots. Dots observed for more than three frames were tracked. **c**, Single-step photobleaching of nucleosome (H2B-Halo-TMR) dots. The vertical axis shows fluorescence intensity of individual TMR dots (± SD among nucleosomes); the horizontal axis is the tracking time series (*N* = 30 nucleosomes). **d**, Representative trajectories of three tracked single nucleosomes. **e**, MSD plots (± SD among cells) of H2B-Halo in HCT116 RAD21-mAC cells under the indicated conditions: DMSO (black, *N* = 20 cells), 5Ph-IAA (ΔRAD21, red, *N* = 20 cells), and FA-fixed (gray, *N* = 15 cells). ***, *P* = 2.1 × 10⁻³ (DMSO vs 5Ph-IAA) and *P* = 5.8 × 10⁻¹² (DMSO vs FA-fixed) by two-sided Kolmogorov-Smirnov test. **f**, Verification of RAD21 depletion by mClover signal in live nuclei. Mean intensity values: 1.6 × 10³ for DMSO (*N* = 20 cells) and 0.010 for +5Ph-IAA (ΔRAD21, *N* = 20 cells). ***, *P* < 0.0001 (*P* = 1.1 × 10⁻⁸) by two-sided Wilcoxon rank-sum test. **g**, Log–log plots of MSD from Fig. 1e. The plots were fitted linearly (dashed lines), and the diffusion exponent α and the coefficient of subdiffusion *D*_sub_ were estimated from the slopes and intercepts of the fitted lines. Note the non-overlapping 95% confidence intervals of the linear fits (shades) between conditions. **h**, Radius of constraints (*Rc*) and asymmetric coefficient (AC) under the indicated conditions: DMSO (*N* = 30 cells), 5Ph-IAA (ΔRAD21, *N* = 30 cells), calculated as Fig. S1g-i.

Cohesin depletion via 5Ph-IAA treatment for 1 hour (Figs. 1f, S1a) led to significantly increased motion, with the increased MSD exponent (0.48) and *D*_sub_ (Fig. 1e,g; Movie S2). The *Rc* (radius of constraint) metric ^55^, which reports the spatial extent of nucleosome motion, was significantly increased (Figs. 1h, S1g). Turning angle distributions of nucleosome trajectories and asymmetry coefficient (AC) analysis ^43,56^ showed reduced pulling-back forces of nucleosomes after cohesin loss, suggesting that cohesin constrains nucleosome movement (Figs. 1h, S1h,i). RAD21 depletion did not induce DNA damage (Fig. S1j). These findings confirm that cohesin constrains local nucleosome motion in living cells, consistent with previous reports ^5,19,40,41^.

### Cohesin constrains local nucleosome motion mainly via loop formation

To determine whether cohesin constrains chromatin via loop domains or sister chromatid cohesion (Fig. 2a), we knocked down Sororin, which is required for sister chromatid cohesion but not for loop formation ^57–59^. Knockdown of Sororin disrupted cohesion in late S–G2 synchronized HCT116 cells (Figs. 2b, S2a) but did not alter nucleosome motion (Fig. 2c), indicating that global constraints arise from loop formation, rather than sister chromatid cohesion. To examine whether cohesion affects local nucleosome motion near cohesion sites, we used a genome labeling system with *tetO*/*lacO* arrays 250 kb apart in chromosome 5 of HT1080 cells (Fig. 2d) ^60^. In this system, the *lacO* array, but not *tetO* array, was located near a cohesin-enriched site (Fig. 2e) ^61^. The *lacO* array more frequently appeared as two separate dots than the *tetO* array, suggesting that the *lacO* array lies near a cohesion site ^60^. These arrays were visualized with TetR-4×mCherry and EGFP-LacI, respectively (Fig. 2d,e). Sororin knockdown (Fig. S2b) significantly increased EGFP-LacI mobility but not TetR-4×mCherry (Fig. 2f,g), whereas RAD21 depletion (Fig. S2c) affected both (Fig. 2f,g), suggesting that cohesion constrains local regions near cohesion sites as previously reported in yeast ^38^, while loop-forming cohesin globally constrains chromatin.

**Fig. 2:**
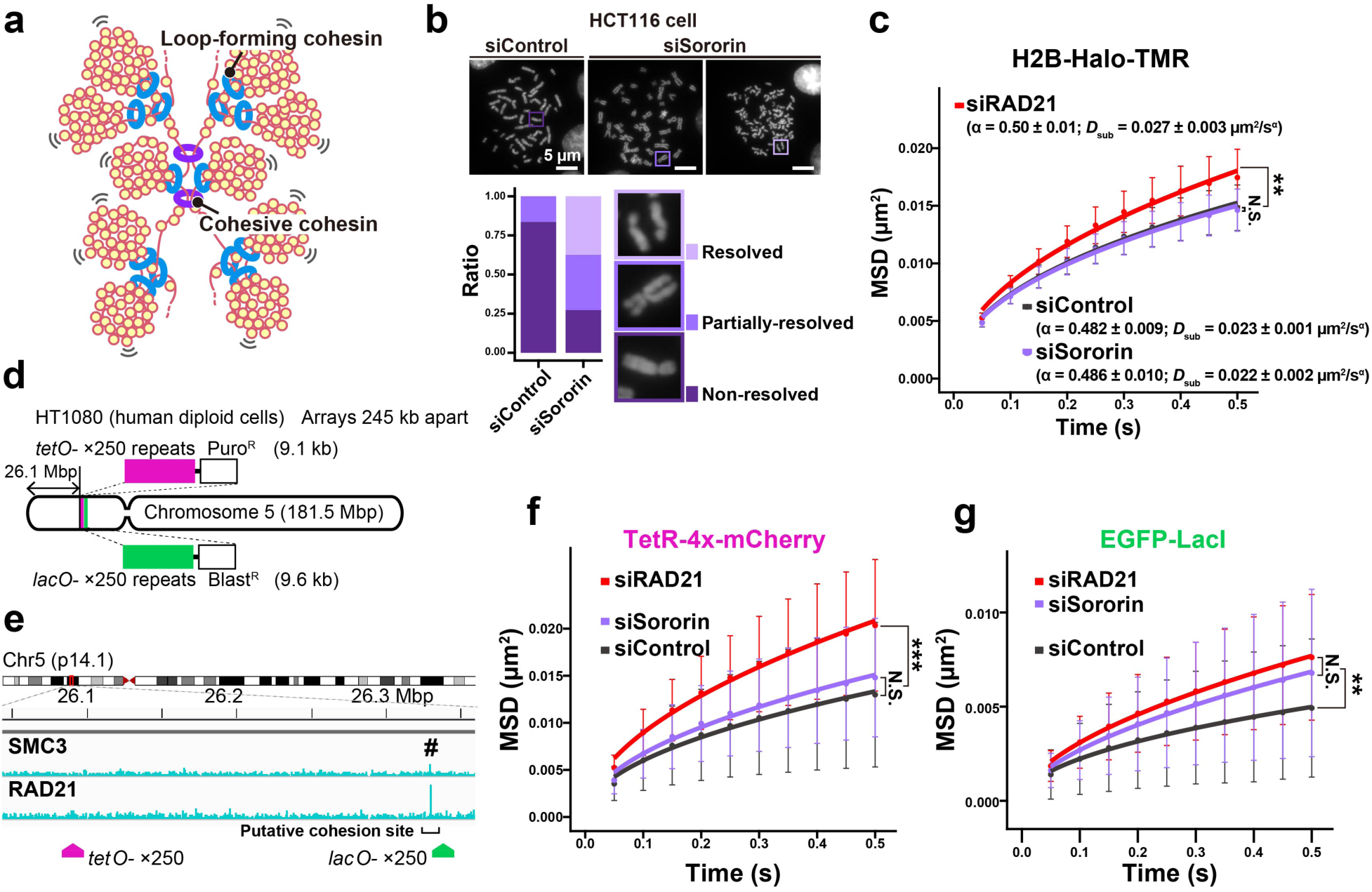
Sister chromatid cohesion does not affect global chromatin motion. **a**, Schematic of loop-forming cohesin and cohesive cohesin. **b**, Chromosome spreads of HCT116 cells with siControl or siSororin. Lower, fraction of chromosomes with the three morphologies. **c**, MSD plots (± SD among cells) of H2B-Halo in late S–G2 synchronized HCT116 cells (*N* = 20 cells) under the indicated conditions: siControl (black), siSororin (purple), and siRAD21 (red). *P* = 7.9 × 10⁻⁴ by the Kruskal–Wallis test, indicating significant overall differences among the groups. **, *P* < 0.01 by Holm-adjusted Dunn’s test for siControl vs siRAD21 (*P* = 1.1 × 10⁻³). N.S., not significant for siControl vs siSororin (*P* = 0.47). **d**, Schematic depicting the locations of *tetO* (×250) and *lacO* (×250) repeats inserted into human chromosome 5. **e**, ChIP-seq data showing the distribution of SMC3 and RAD21 across the genomic region around the integration sites of *lacO* and *tetO*. ChIP-seq data are from published data ^61^. # shows the position of SMC3 and RAD21 peaks. **f**, MSD plots (± SD among cells) of *tetO*–TetR–4×mCherry in late S–G2 synchronized HT1080 cells under the indicated conditions: siControl (black, *N* = 34 cells), siSororin (purple, *N* = 36 cells), and siRAD21 (red, *N* = 37 cells). *P* = 3.7 × 10⁻⁵ by the Kruskal–Wallis test, indicating significant overall differences among the groups. ***, *P* < 0.001 by Holm-adjusted Dunn’s test for siControl vs siRAD21 (*P* = 1.9 × 10⁻⁵). N.S., not significant for siControl vs siSororin (*P* = 0.10). Note that the following two factors make locus MSD noisier than single-nucleosome imaging/tracking. First, each locus provides only one trajectory per cell, so sampling is sparse. Another factor is that the *lacO*/EGFP-LacI and *tetO*/TetR-4×mCherry foci are multi-emitter spots (many LacI/TetR) whose shape/intensity fluctuate, reducing localization precision. **g**, MSD plots (± SD among cells) of *lacO*–EGFP–LacI in late S–G2 synchronized HT1080 cells under the indicated conditions: siControl (black, *N* = 22 cells), siSororin (purple, *N* = 31 cells), and siRAD21 (red, *N* = 26 cells). *P* = 1.1 × 10⁻² by the Kruskal–Wallis test, indicating significant overall differences among the groups. **, *P* < 0.01 by Holm-adjusted Dunn’s test for siControl vs siRAD21 (*P* = 4.0 × 10⁻³). N.S., not significant for siRAD21 vs siSororin (*P* = 0.11). f and g were derived from pooled data from two experiments.

We tested whether increasing the number of loops enhances constraint. Since WAPL releases cohesin from chromatin, WAPL depletion increases the number of loops ^62–64^, particularly when depletion is induced after the completion of DNA replication. WAPL depletion in HCT116-WAPL-mAC cells (Fig. 3a) ^65^ synchronized in late S-G2 phase (Fig. S3a) reduced the WAPL-mClover signal (Fig. 3b) and increased RAD21 chromatin binding (Fig. S3b), which led to decreased MSD and AC values (Fig. 3c). A similar finding was also observed in asynchronous HCT116 cells and siRNA-treated asynchronous HeLa cells (Fig. S3c–e). Thus, more cohesin loops constrain chromatin more strongly. To assess whether loop boundaries contribute to chromatin constraint, we acutely depleted CTCF in HCT116 cells (Fig. 3d,e) ^53^. Despite its role in TAD boundary formation ^66^, CTCF loss did not affect nucleosome motion (Figs. 3f, S3f,g), consistent with unchanged cohesin binding levels ^50,67^ and previous genomic array data ^40,41^.

**Fig. 3:**
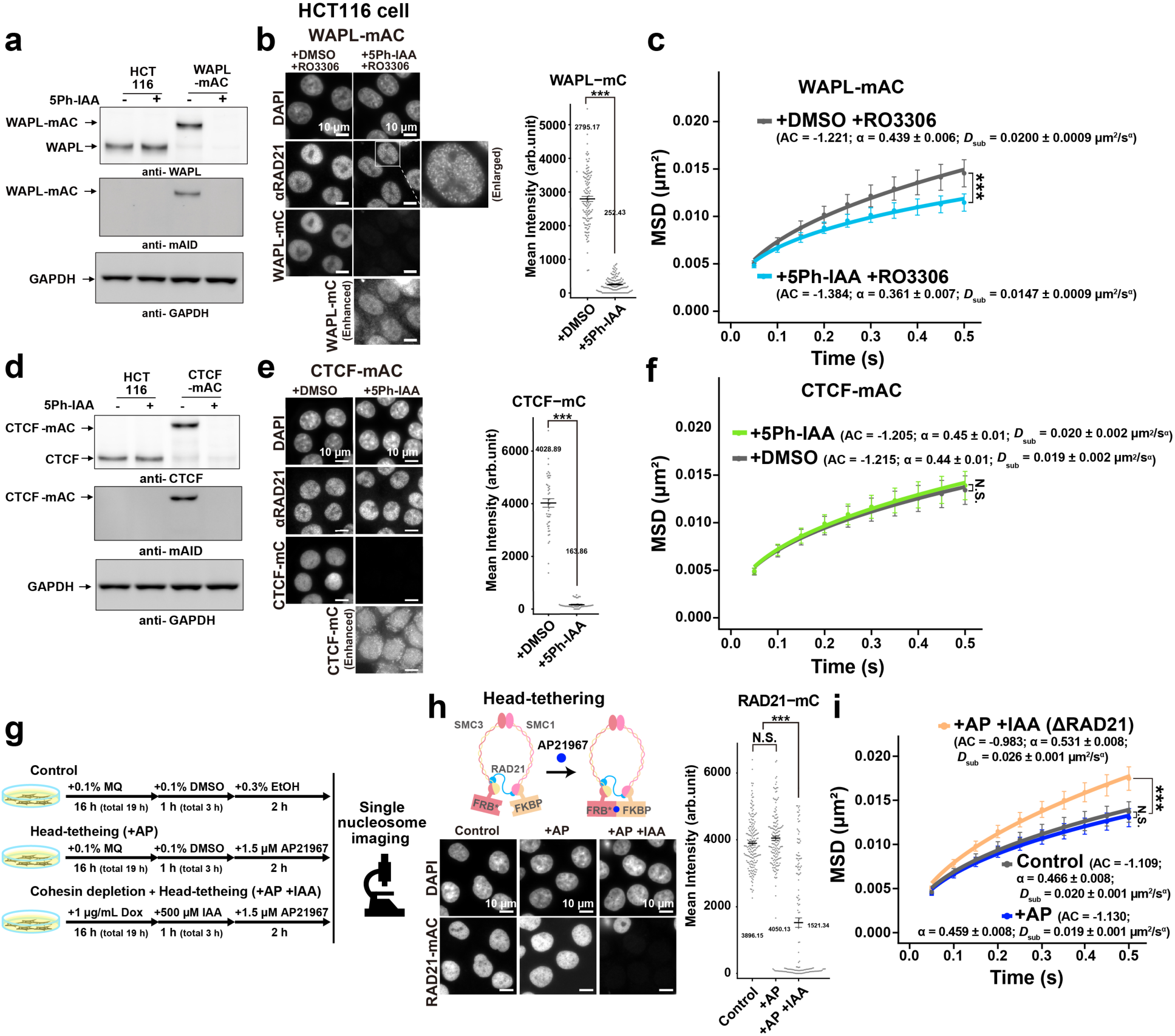
Nucleosome motion changes in response to altered cohesin activity. **a,b**, Verification of WAPL depletion by western blot (a) and mClover (mC) fluorescent signal (b). The enlarged image in panel (b) highlights the vermicelli-like structure in RAD21 staining, consistent with ^62^. Mean intensity values: 2.8 × 10³ for DMSO (*N* = 118 cells) and 2.5 × 10² for 5Ph-IAA (ΔWAPL, *N* = 124 cells). ***, *P* < 0.0001 (*P* = 4.7 × 10⁻⁴¹) by Wilcoxon rank-sum test. **c**, MSD plots (± SD among cells) of H2B-Halo in late S–G2 synchronized HCT116 cells (*N* = 30 cells) with indicated conditions: DMSO (black) or 5Ph-IAA (light-blue). ***, *P* = 8.3 × 10⁻¹² for DMSO vs 5Ph-IAA by the two-sided Kolmogorov–Smirnov test. **d,e**, Verification of CTCF depletion by western blot (d) and mClover fluorescent signal (e). Mean intensity values: 4.0 × 10³ for DMSO (*N* = 47 cells) and 1.6 × 10² for 5Ph-IAA (*N* = 46 cells). ***, *P* < 0.0001 (*P* = 1.0 × 10⁻¹⁶) by Wilcoxon rank-sum test. **f**, MSD plots (± SD among cells) of H2B-Halo in HCT116 cells with indicated conditions: DMSO (black, *N* = 21 cells) or 5Ph-IAA (light-green, *N* = 30 cells). N.S., not significant for DMSO vs 5Ph-IAA (*P* = 0.67) by the two-sided Kolmogorov– Smirnov test. **g**, Time course of head-tethering experiment. **h**, Verification of RAD21 depletion by mClover signal. Mean intensity values: 3.9 × 10³ for Control (*N* = 184 cells), 4.1 × 10³ for AP21967 (AP; *N* = 161 cells), and 1.5 × 10³ for AP+IAA (*N* = 157 cells, DOX pre-treated for the expression of OsTIR1 ubiquitin ligase). N.S., not significant for Control vs AP (*P* = 0.073). ***, *P* < 0.0001 for Control vs AP+IAA (*P* = 2.3 × 10⁻²⁸) by Wilcoxon rank-sum test. Note that only half of the population depleted RAD21. For single-nucleosome imaging, we selected depleted cells for analysis. **i**, MSD plots (± SD among cells) of H2B-Halo in HCT116 cells with indicated conditions: Control (black, *N* = 20 cells), AP (blue, *N* = 30 cells), and AP+IAA (orange, *N* = 20 cells). *P* = 1.0 × 10⁻³ by Kruskal–Wallis test, indicating significant overall differences among the groups. ***, *P* < 0.001 by Holm-adjusted Dunn’s test for Control vs AP+IAA (*P* = 1.4 × 10⁻⁵). N.S., not significant for Control vs AP (*P* = 0.11).

Finally, to examine the possible involvement of active loop extrusion ^32,68^ in the constraining process, we used a head-tethered cohesin system ^69^ that stabilizes cohesin via forced head domain closure using AP21967 (Fig. 3g,h) ^70,71^. Head-tethering weakened loop-extrusion activity *in vitro* ^69^. Head-tethering alone did not change nucleosome motion on the 0.5 s timescale, but head-tethering after RAD21 depletion led to the loss of the entire cohesin complex ^69^ and increased motion (Figs. 3i, S3h,i). These results suggest that chromatin constraint by cohesin may not require active loop extrusion. Also, the difference between head tethering and WAPL depletion explains the distinct outcomes. WAPL depletion suppresses cohesin dissociation, allowing cohesin to remain bound and continue to accumulate on chromatin. By contrast, head-tethered cohesin is immobilized at existing sites and does not promote new chromatin loading. Consequently, WAPL depletion yields much higher cohesin occupancy and stronger chromatin constraint than head tethering (Fig. 3c,i).

We previously observed heterogeneous nucleosome motions (Fig. S4a,b), which suggested the presence of domain subpopulations, potentially linked to chromatin context ^24,28^. To get a clue of whether cohesin preferentially constrains euchromatin or heterochromatin, we used motion classification with the Bayesian-based Richardson-Lucy algorithm (RL algorithm) ^72^. For RL algorithm analysis, we obtained a 10-fold increase in H2B-Halo nucleosome trajectory data using the PA-JF646 HaloTag ligand ^73^ and constant photoactivation with weak 405 nm illumination (Figs. S4c-f). We then classified H2B-Halo trajectories into four motion types: “Super-slow,” “Slow,” “Fast,” and “Super-fast” (Fig. S4g; Movie S3). The “Fast” and “Super-fast” subpopulations are likely euchromatic, localized within the nucleoplasm (Fig. S4h; Movie S3). Cohesin depletion shifted these populations rightward (Fig. S4g), increasing their mobility and anomalous exponent α and *D*_sub_ values (Figs. S4i-k), while the proportions remained similar (Fig. S4l), implying that cohesin might constrain movements of nucleosomes mainly in euchromatin. Our results also indicate that RL analysis can extract new information from bulk motion data that cannot be obtained using conventional MSD analysis.

### Euchromatin-specific nucleosome labeling reveals that cohesin constrains euchromatin

To investigate euchromatin more specifically, we labeled nucleosomes using a euchromatin-specific histone variant H3.3 ^47–49^. We stably expressed H3.3-Halo in HCT116 cells (Fig. 4a).

**Fig. 4:**
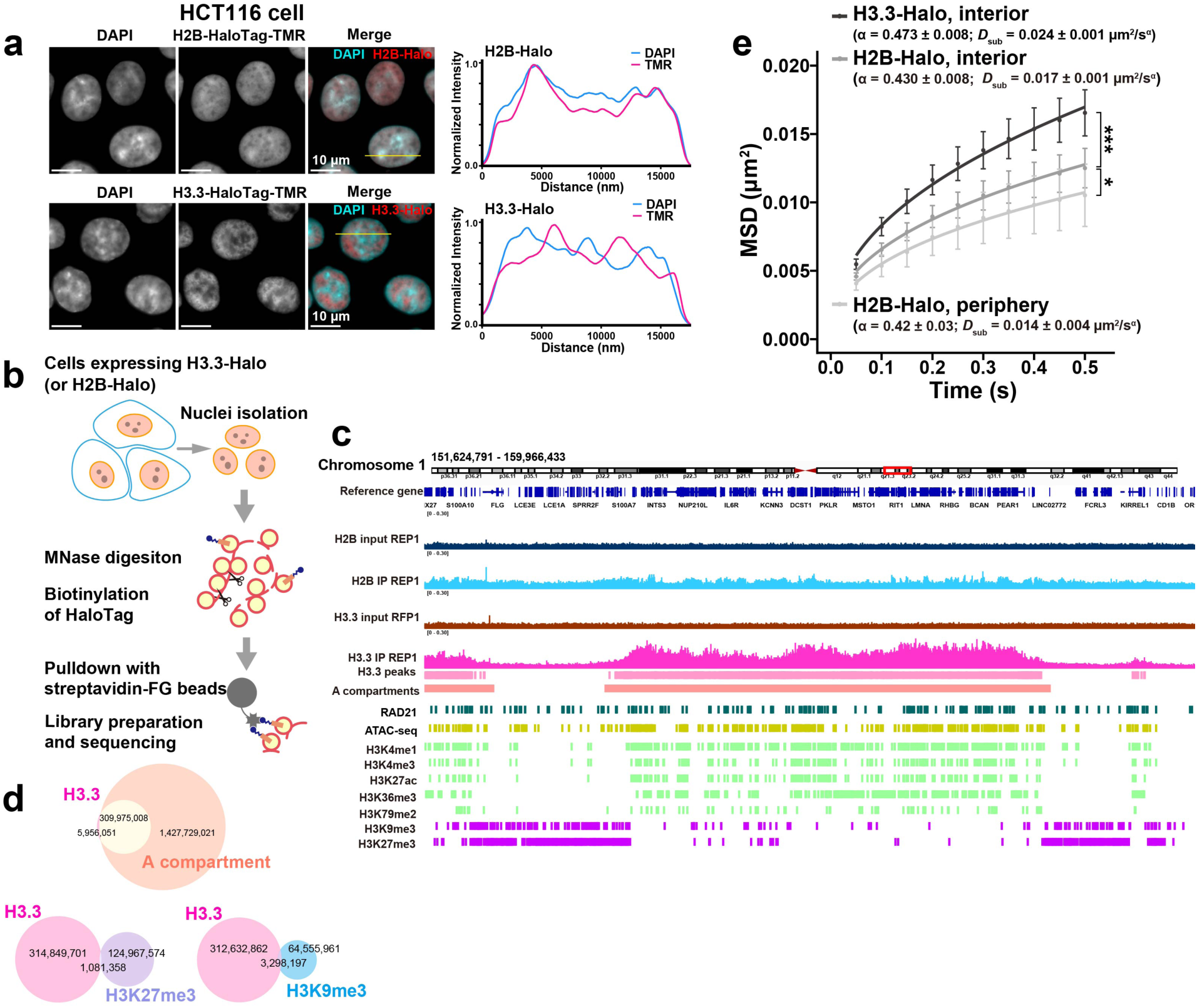
H3.3-Halo is enriched in euchromatin (Hi-C A compartment). **a** (upper), HCT116 cells expressing H2B-Halo labeled with DAPI and TMR. Right, intensity line profiles of DAPI and H2B-Halo at the yellow line of the merged image. Note that the localization of H2B-Halo is similar to the DAPI-stained DNA. (lower) HCT116 cells expressing H3.3-Halo labeled with DAPI and TMR. Right, intensity line profiles of DAPI and H3.3-Halo at the yellow line of the merged image. Note that H3.3-Halo is enriched at DAPI-less (euchromatin) regions. **b**, A purification scheme for nucleosomes with H3.3-Halo or H2B-Halo and their associated genomic DNA. The enriched DNA fractions were then indexed, amplified, and sequenced. **c**, H3.3-Halo–incorporated genomic regions were enriched with active histone marks and excluded from an inactive mark: view from chr1, 151,624,791–159,966,433 (also see Fig. S5d). **d**, Overlap of the detected between H3.3-Halo peaks and the A compartment ^131^, between H3.3-Halo peaks and H3K27me3 regions, and between H3.3-Halo peaks and H3K9me3 regions. **e**, MSD plots (± SD among cells) of H2B-Halo (nuclear interior; *N* = 30 cells), H2B-Halo (nuclear periphery (basal nuclear surface); *N* = 8 cells), or H3.3-Halo (nuclear interior; *N* = 30 cells) in HCT116 cells. ***, *P* = 8.3 × 10⁻¹² for H2B-Halo, interior vs H3.3-Halo, interior by the two-sided Kolmogorov–Smirnov test, *, *P* = 4.2 × 10⁻^3^ for H2B-Halo, interior vs H2B-Halo, periphery by the two-sided Kolmogorov–Smirnov test.

Expressed H3.3-Halo was properly incorporated into the nucleosome (Fig. S5a) and localized to DAPI-poor regions in nuclei (Fig. 4a). Pull-down of H3.3-Halo-incorporated nucleosomes and genomic sequencing (Figs. 4b, S5b,c) confirmed enrichment in Hi-C A compartments (98.1%) with active histone marks, and depletion from repressive marks (Figs. 4c,d, S5d–r). We observed a similar trend in HeLa cells stably expressing H3.3-Halo as well (Fig. S6).

Single-nucleosome imaging in HCT116 cells showed that H3.3-Halo nucleosomes were more mobile than H2B-Halo ones (Figs. 4e, S5s) and had a stronger response to RAD21 or WAPL depletion (Fig. S7a,c)^53,65^, indicating a greater cohesin impact in euchromatin. In contrast, H2B-labeled nucleosomes around the nuclear periphery (basal nuclear surface; for details, see Methods), which is enriched with constitutive heterochromatin, or lamin-associated domains (LAD) ^74^, remained largely unaffected (Fig. S7b,d). A similar observation was found in depletion of RAD21 and WAPL via siRNA in HeLa cells expressing H3.3-Halo (Fig. S7e) and H2B-Halo around the nuclear periphery (Fig. S7f). These results suggest that cohesin primarily constrains euchromatic regions while our results do not exclude the possibility that cohesin may also influence facultative heterochromatin regions under certain conditions.

### Super-resolution imaging uncovers condensed euchromatic domains in live human cells

The finding that cohesin constrains euchromatic nucleosome motion motivated us to explore how cohesin affects the structure and physical properties of euchromatic domains. For this purpose, we used 3D-structured illumination microscopy (3D-SIM) ^7,44–46^ on HCT116 cells expressing H3.3-Halo and visualized euchromatic domain structures in live or formaldehyde (FA)-fixed cells at super-resolution. 3D-SIM enables fast multicolor 3D acquisition of whole cells with higher resolution (xy ∼130 nm; z ∼250 nm) and strongly enhanced contrast compared to conventional fluorescence microscopy (panels 1–2 and 4–5 in Fig. 5a; Movie S4) ^7,45^. Using quality-controlled 3D-SIM (Figs. S8a-e and Methods) ^75^, we found that euchromatin in live HCT116 cells appeared as connected, condensed chromatin domains with irregular shapes and diameters in the range of a few hundred nanometers (Fig. 5b; Movie S5). A similar architecture was observed in FA-fixed cells labeled with H3.3-Halo (panel 5 in Fig. 5a).

**Fig. 5:**
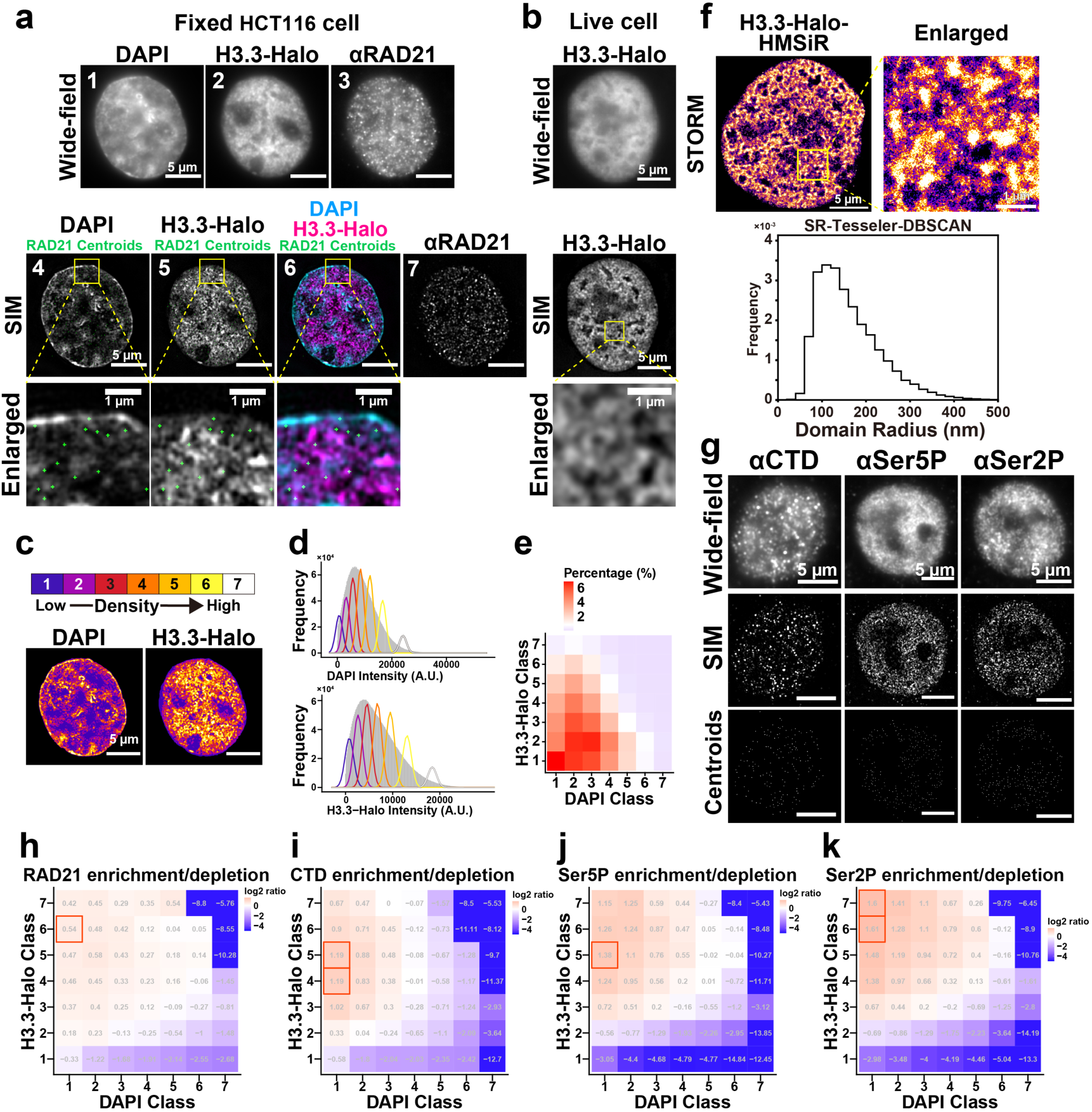
3D-SIM imaging revealed condensed euchromatic domains in human cells and localization of cohesin and RNA polymerase II. **a**, Representative wide-field view (panels 1–3) and 3D-SIM reconstructed view (panels 4–6) of FA-fixed HCT116 cells. Chromatin was labeled with DAPI (panels 1, 4) and H3.3-Halo-TMR (panels 2, 5). Immunostaining wide-field image of RAD21 is also shown (panel 3). Panel 6 is a merged image of panels 4 and 5. Bottom images are enlarged views of the boxed regions in panels 4–6. Panel 7 is a SIM image of RAD21 immunostaining. The wide-field image (panels 1– 3) represents the total intensity projection of 15 raw SIM frames. Green dots in the enlarged images indicate the centroids of RAD21 immunostaining signals from panel 7. **b**, Representative wide-field view (top row) and 3D-SIM reconstructed view (bottom row) of live HCT116 cells. **c**, Heatmap of DAPI or H3.3-Halo compaction proxies, based on intensity classification into seven classes ^144^. The color code assigns DAPI or H3.3-Halo signals into seven classes with equal intensity variance. This approach enables threshold-independent signal intensity classification based on the intensity of individual voxels. **d**, Histogram of the DAPI (upper) or H3.3-Halo (lower) intensity from a representative nucleus image (gray), overlaid with probability density functions of the seven Gaussians generated by the classification analysis. A.U., arbitrary units. **e**, Proportion of nuclear volume across 7 × 7 categories based on DAPI and H3.3-Halo intensity classes in a representative nucleus. **f**, Representative STORM reconstructed view of H3.3-Halo**-** HMSiR. Histogram of estimated radius of euchromatic domains (bottom) by SR-Tesseler-DBSCAN pipeline (see Methods). **g**, Representative images of immunostaining signals of various RNA polymerase II (RNAP II) antibodies. Wide-field view (top), 3D-SIM reconstructed view (middle), and detected centroids of immunostaining signals (bottom). The wide-field image represents the total intensity projection of 15 raw SIM frames. Note that 3D-SIM reconstruction efficiently suppresses out-of-focus blur and therefor provides highly confined optical sectioning with a narrow depth of field. **h–k**, Each voxel within a single nucleus was classified into 7 × 7 categories based on DAPI and H3.3-Halo intensity classes. Target signal enrichment/depletion is shown as the log₂ ratio of the signal in each class to the relative volume of that class. The displayed values represent the median across multiple cells. For categories with no detected signal, a small constant was added to avoid undefined log values; these categories, with resulting values below –5, are shown in blue. (h) RAD21, (i) RNAP II CTD, (j) RNAP II Ser5P, (k) RNAP II Ser2P enrichment/depletion. The same data are used as in Fig. S8i-l.

Furthermore, we used STORM imaging of H3.3-Halo-HMSiR in HCT116 cells, another super-resolution method with ∼5 nm resolution (Fig. S8f) ^5,76^ ^77^, to investigate euchromatic domain organization. Reconstructed STORM images revealed condensed euchromatin domains (Fig. 5f) similar to those observed by 3D-SIM. Clustering analysis of nucleosomes using SR-Tesseler-DBSCAN pipeline ^78^ yielded a peak domain radius of ∼100–120 nm (diameter of ∼200–240 nm, Fig. 5f). These findings of condensed euchromatic domains in living human cells challenge the established view of euchromatin as largely open (Fig. S8g-h).

### Preferential localization of cohesin to euchromatin domains

We investigated the spatial distributions of cohesin and active transcription sites within the nucleus by combining 3D-SIM with immunostaining. RAD21 immunostaining revealed that cohesin appeared as punctate foci in SIM images (panel 7 in Fig. 5a), likely representing chromatin-bound cohesin. The centroids of these foci were frequently located near the edges of H3.3-Halo domains, suggesting that cohesin localizes to the surfaces of euchromatin domains (enlarged images in Fig. 5a). Classification analysis of signal density in 3D-SIM ^7,79^ confirmed that cohesin is enriched in high-density H3.3 regions (classes 3–7; Figs. 5c-e, S8i), indicating a preferential localization of cohesin to euchromatin domains. Consistently, cohesin enrichment was observed in low-density DAPI regions (classes 1–2) and depleted in denser regions (Fig. S8j), in good agreement with previous observations ^7^. This spatial distribution of cohesin also supports our finding that euchromatic nucleosome motion is more sensitive to cohesin depletion (Fig. S7a,b).

### Active RNAP II localizes around euchromatic domain surfaces

We next examined the localization of active transcription sites. We performed immunostaining using antibodies against the C-terminal domain (CTD) of RNA polymerase II (RNAP II) and its phosphorylated forms at serine 5 (Ser5P) and serine 2 (Ser2P) to visualize RNAP II signals composed predominantly of inactive, active paused, and elongating RNAP II, respectively (Fig. 5g). All signals appeared as punctate foci (Fig. 5g). These signals were enriched in DAPI-poor regions (classes 1–2; Fig. S8k) and in H3.3-Halo-rich regions (classes 4–7; Fig. S8l), indicating a euchromatin-biased localization.

We observed distinct differences in nuclear localization among cohesin and RNAP II forms. Cohesin showed a relatively uniform distribution with a preference for euchromatin (Fig. 5h). In contrast, RNAP II showed increased localization to H3.3-denser areas upon transcriptional activation (Fig. 5i–k). Within the H3.3-Halo classification, CTD signals peaked at classes 4–5 (Fig. 5i), Ser5P at class 5 (Fig. 5j), and Ser2P at classes 6–7 (Fig. 5k). These patterns suggest that inactive RNAP II remains at the periphery (or outside) of euchromatic domains, paused active RNAP II localizes near the domain surface, and elongating RNAP II may penetrate the domain.

Furthermore, we investigated the relative genomic locations of RNAP II, cohesin, and other histone marks to H3.3-Halo domains (Figs. 6, S8m-s). Consistent with the 3D-SIM imaging data, RNAP II, cohesin, and active enhancers/promoters are preferentially located on the edge of the H3.3 domain, which corresponds to the surface of condensed euchromatic domains in 3D (Fig. 6a).

**Fig. 6:**
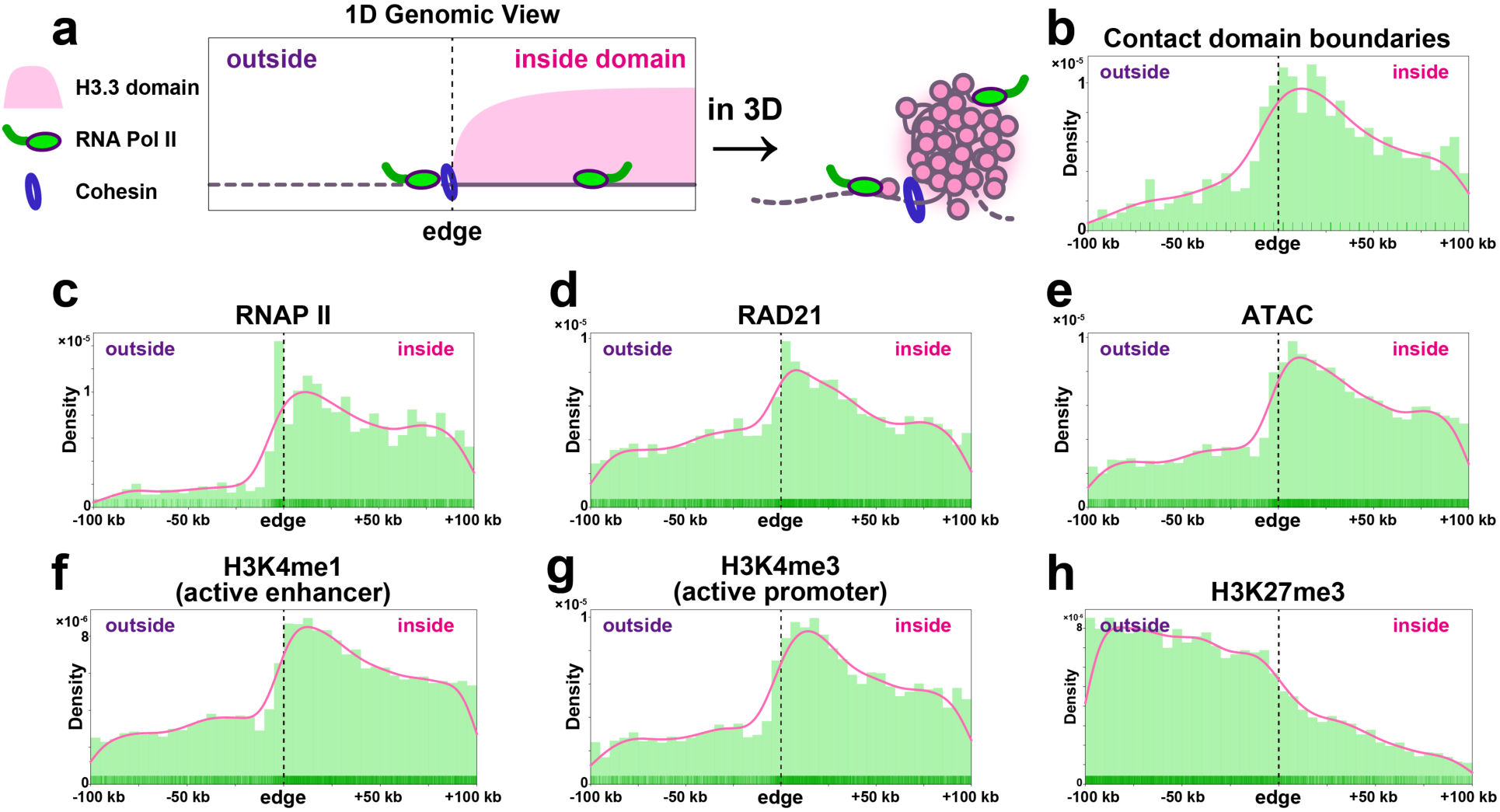
RNA polymerase II and cohesin localize on the edge, which is the surface of condensed euchromatic domains. **a**, Relative position of RNAP II, cohesin, or other markers to the edge of H3.3 domains (median: 210 kb; see also Fig. S8m) is examined. The H3.3 domain is detected from the pull-down experiment (Fig. 4). Molecules on the edge of an H3.3 domain in 1D will be localized on the surface of a condensed euchromatic chromatin domain in 3D. **b-h**, Distribution of (b) contact domain (or TAD) boundaries ^87^, (c) RNAP II ^128^, (d) RAD21 ^128^, (e) ATAC-seq peaks, (f) H3K4me1 ^87^, (g) H3K4me3 ^87^, and (h) H3K27me3 ^87^ relative to the closest edge of a H3.3 domain. Positive and negative values indicate inside and outside of H3.3 domains, respectively. The pink line is a kernel density estimation on the distribution histogram. See also Fig. S8n-s for other markers.

### Cohesin depletion does not alter overall compaction of euchromatic domains

To assess the effect of cohesin depletion on chromatin organization, especially euchromatic domains, we acutely depleted RAD21 in HCT116 cells expressing H3.3-Halo and visualized their chromatin organization using 3D-SIM. Interestingly, no obvious qualitative changes were observed in the overall chromatin organization (DAPI; Fig. 7a; Movies S6–S7) or in the euchromatic domains (H3.3-Halo; Fig. 7b) upon cohesin depletion. For quantitative analysis, we classified DAPI and H3.3-Halo signals. Cohesin depletion did not change the relative volumes of each class, i.e., the compaction levels of DAPI-stained chromatin (Fig. S9a) and H3.3-Halo– labeled euchromatin domains (Fig. S9b), in agreement with previous findings ^7^.

**Fig. 7:**
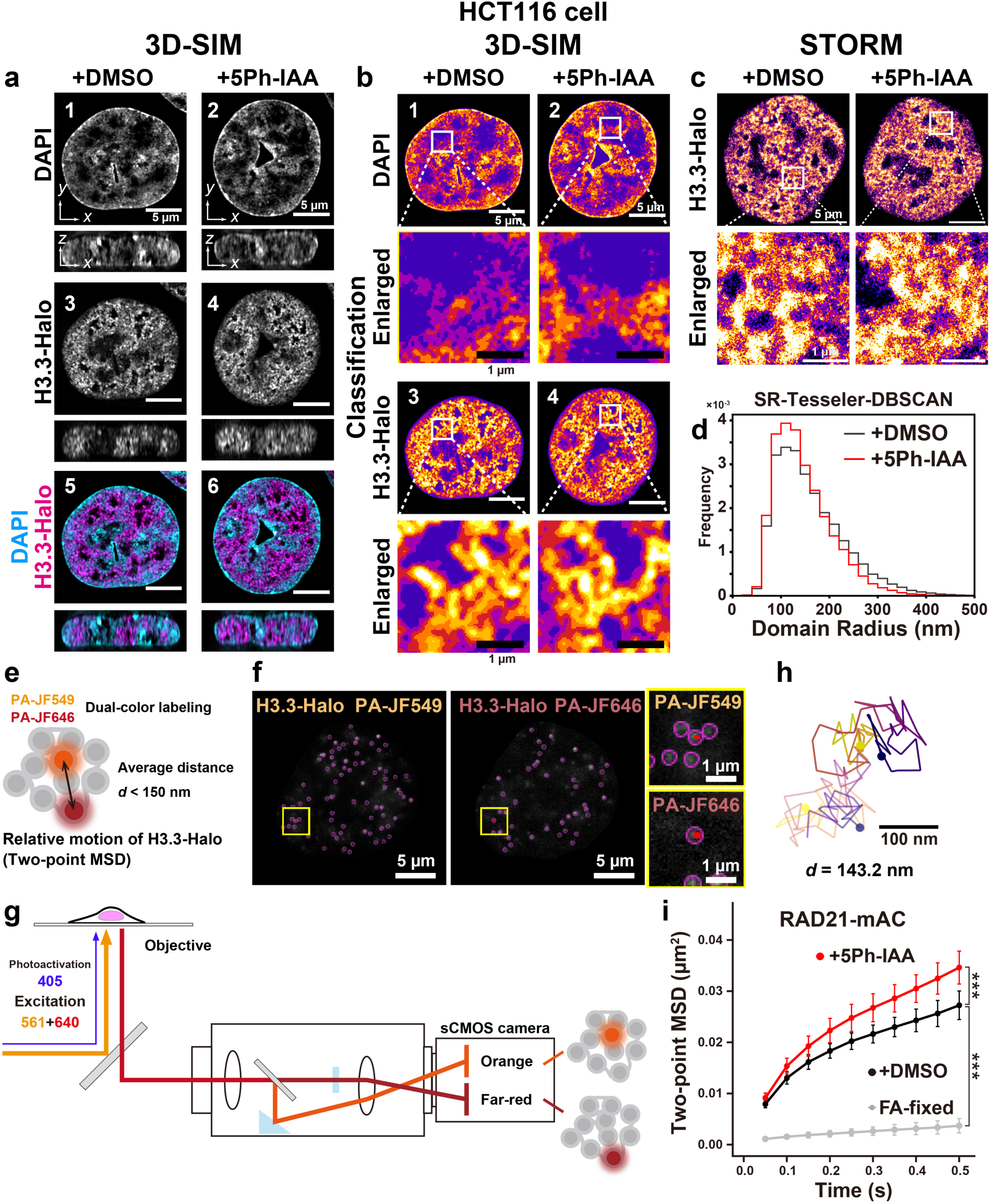
3D-SIM and STORM imaging and two-point MSD analysis of condensed euchromatic domains. **a**, Representative single z-slices of 3D-SIM images on chromatin labeled with DAPI (panels 1 and 2) or H3.3-Halo-TMR (panels 3 and 4) with DMSO treatment (panels 1 and 3) or cohesin depletion (panels 2 and 4). Panels 5 and 6 are merged images. Their Z-X views are also shown below each image. **b**, Heatmaps of DAPI or H3.3-Halo compaction proxies, based on intensity classification into seven classes. Panels 1-4 correspond to the same panel number in (a). Enlarged images from the corresponding squared regions in panels 1-4 are also shown below each panel. **c**, Representative STORM reconstructed view of FA-fixed HCT116 RAD21-mAC H3.3-Halo cells. Chromatin was labeled with H3.3-Halo-HMSiR with DMSO treatment (+DMSO) or cohesin depletion (+5Ph-IAA). Bottom images are enlarged views of the boxed regions in the upper panels. Euchromatic chromatin is condensed. **d**, Histogram of estimated radius of euchromatic domains by SR-Tesseler-DBSCAN pipeline (see Methods.) **e**, Scheme for dual-color imaging of H3.3-Halo and two-point MSD. **f**, A single-nucleosome (H3.3-Halo-PAJF dyes) image of a living HCT116 nucleus after background subtraction. Magenta circles highlight the tracked dots. A pair of dots with an average inter-point distance *d* < 150 nm (square) is shown in the zoomed-in panel with red trajectories. **g**, Schematic for dual-color imaging with a beam splitter system (W-VIEW GEMINI, Hamamatsu Photonics). The images of two single-nucleosomes with different colors were acquired with a single sCMOS camera (left half, orange; right half, far-red.). **h**, Representative pair trajectories with average distance *d* < 150 nm. Darker color: PA-JF549 labeled H3.3-Halo; lighter color: PA-JF647 labeled H3.3-Halo. Trajectories are shown with a yellow-to-navy gradation from starting to ending points. **i**, Two-point MSD plots (*d* < 150 nm; ± SD among cells) of H3.3-Halo in HCT116 cells with indicated conditions: DMSO (black, *N* = 20 cells), 5Ph-IAA (ΔRAD21, red, *N* = 25 cells), FA-fixed (gray, *N* = 25 cells). ***, *P* < 0.001 by the two-sided Kolmogorov-Smirnov test for DMSO vs 5Ph-IAA (*P* = 1.2 × 10^-4^), for DMSO vs FA-fixed (*P* < 2.2 × 10^-16^).

Reconstructed STORM images after cohesin depletion showed a similar condensed domain organization (Fig. 7c). SR-Tesseler-DBSCAN analysis ^78^ of the reconstructed images further reveals that cohesin depletion slightly increased the fraction of domains with smaller radii (∼100–120 nm) and decreased the large-radius tail (∼200–400 nm) (Fig. 7d), consistent with domain splitting upon cohesin loss ^8,20^. The modal radius was largely maintained. Together, these super-resolution results indicate that cohesin depletion retains overall domain compaction while increasing nucleosome mobility.

### Nucleosome–nucleosome interactions and cohesin are involved in condensed domain formation

To investigate which factors condense euchromatic domains, we treated the cells with the HDAC inhibitor trichostatin A (TSA) ^80^, which induces hyperacetylation of histone tails (Fig. S9c,d) ^81^. Tail hyperacetylation weakens local nucleosome–nucleosome interactions ^82^ and decondenses chromatin ^5,83,84^. Indeed, mitotic chromosomes were highly decondensed and expanded in volume in condensin-depleted and TSA-treated cells ^85,86^. Following treatment with TSA for 7 h and acute depletion of RAD21 (Fig. S9c,d), apparent decondensation of H3.3-Halo domains was observed by 3D-SIM (Fig. S9e). Power-spectrum analysis showed that the amplitude at spatial scales of ∼300–600 nm (domain scale) decreased substantially (Fig. S9f,g). Together, these results indicate that electrostatic interactions between positively charged histone tails and DNA/histones, together with cohesin, contribute to condensed domain formation. Moreover, these results show that our 3D-SIM was able to detect chromatin decondensation and confirm that the condensed euchromatic domains were not an imaging artifact. Our observations provide compelling evidence that challenges the established view of euchromatin as largely open (Fig. S8g,h).

### Cohesin regulates the fluidity of nucleosomes inside condensed euchromatic domains

Although euchromatic nucleosome movement increases upon cohesin depletion, euchromatic domain compaction remains largely unchanged. This raised the question of whether nucleosomes might gain mobility within the domains. To address this, we introduced dual-color labeling (Fig. 7e, S9h,i) and tracking of two spatially neighboring single H3.3-Halo-nucleosomes (Fig. 7f-h; Movie S8). Since simple single-nucleosome imaging cannot distinguish whether individual nucleosomes fluctuate within the domain or whether the domain itself moves, we measured the distances between two neighboring H3.3-Halo-nucleosomes within euchromatic domains or possibly between adjacent domains (average distance *d* <150 nm; Fig. 7e) ^8^ and obtained the MSD of their relative positions (two-point MSD) ^8,28,40^. As a control, we analyzed two-point MSD for unrelated (long distance) H3.3–Halo pairs (300 nm < *d* < 1000 nm) (Fig. S9j).

Two-point MSD analysis showed much greater motion between neighboring H3.3-Halo-nucleosomes in living cells than in FA-fixed cells (Fig. 7i), suggesting that nucleosomes fluctuate within the domain. Notably, two-point MSD increased upon acute cohesin depletion (1 h 5Ph-IAA) (Figs. 7i, S9k-m). Longer depletions (3 h or 6 h 5Ph-IAA) did not further change the two-point MSD (Fig. S9k). These results indicate that cohesin depletion increases nucleosome mobility within condensed euchromatic domains and induces a transition to a more liquid-like state of chromatin, or to an unrestricted state of the chromatin polymer.

Two-point MSD for unrelated (long distance) control pairs was approximately twice the one-point MSD at matched lags (Fig. S9j), as theory predicts for uncorrelated motion. By contrast, neighboring pairs showed lower two-point MSD, consistent with positively correlated motion within domains. In both classes, RAD21 depletion increased two-point MSD. Taken together, neighboring-pair two-point MSD reveals the extent of local physical constraints within euchromatic domains.

### Local domain mixing upon cohesin depletion

As described above, cohesin loss increased nucleosome fluidity within condensed euchromatic domains. In this state, domains may become less constrained and more flexible, permitting inter-domain mixing. Consistently, Hi-C contact probability plot in HCT116 cells ^87^ shows that cohesin loss decreases short-range contacts (∼0.1–1.5 Mb, TAD scale) and increases long-range contacts (Mb scale) (Fig. 8a). This effect is slightly enhanced within A compartment regions (Fig. S10a). These findings provide genomic evidence of “domain mixing” upon cohesin depletion and align with previous studies ^19,88^.

**Fig. 8:**
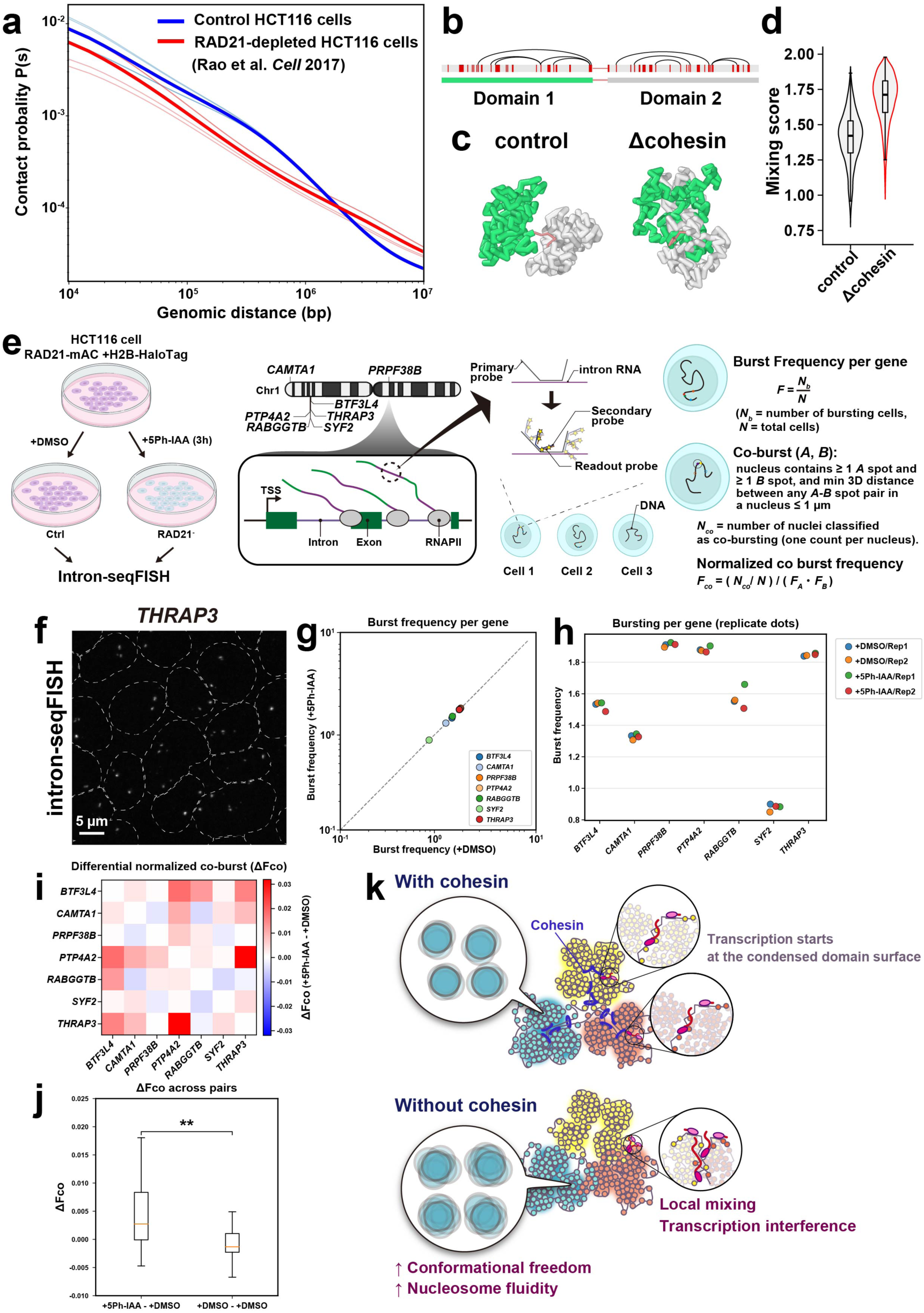
Cohesin prevents local mixing of condensed euchromatic domains. **a**, Hi-C contact probability *P*(*s*) for HCT116 cells with cohesin (control, blue) and after RAD21 depletion (red) were plotted from the Hi-C data of Rao et al. ^87^. Upon RAD21 depletion, short-range (up to ∼1.5 Mb) contacts decrease, whereas long-range (several-Mb scale) contacts increase, consistent with local mixing of chromatin domains at multi-megabase scales after cohesin depletion. **b**, An example chain composed of two 200-kb domains, Domain 1 (green) and Domain 2 (gray), connected by a short linker. The regions containing acetylated nucleosomes are colored red. In the control case, five cohesin molecules are loaded to form five loops within each domain, represented by arcs. **c**, Snapshots of the configurations of the two consecutive domains (green and gray). Cases with cohesin (control) and without cohesin (Δcohesin). **d**, Distributions of the mixing score (see Methods). **e,** Schematic of the intron-seqFISH workflow and the definitions of burst and co-burst metrics used in this study. HCT116 RAD21-mAC+H2B-Halo cells were treated with DMSO or 1 µM 5Ph-IAA for 3 h and then subjected to intron-seqFISH. Intronic nascent transcripts from seven genes located on chromosome 1 (*CAMTA1, PTP4A2, RABGGTB, SYF2, THRAP3, BTF3L4,* and *PRPF38B*) were detected. See also Fig. S10g, h. **f**, Representative image obtained by intron-seqFISH in fixed HCT116 RAD21-mAC H2B-Halo cells. In each hybridization round, nascent intronic RNA from a single target gene was visualized (*THRAP3* for this image). Dashed lines indicate nuclear boundaries determined from DAPI images. **g**, Burst frequency of individual genes before and after RAD21 depletion. **h**, Burst frequency per gene shown for each biological replicate. **i**, Heatmap of differential normalized co-bursting (Δ*F_co_*) for all gene pairs. **j**, Distribution of Δ*F_co_* across all gene pairs. Δ*F_co_* values comparing RAD21⁻ and control conditions (left) are shown alongside control replicate differences (Ctrl/Rep2 − Ctrl/Rep1; right) as a baseline. **, P < 0.01 (P = 0.008) by Wilcoxon rank sum test. **k**, Cohesin holds chromatin fibers together and constrains chromatin motion within condensed euchromatic domains. While cohesin depletion does not alter the compaction level of these domains, it increases nucleosome fluidity (conformational freedom). This enhanced fluidity upon cohesin loss may promote mixing between chromatin domains. Thus, cohesin stiffens condensed euchromatic domains, preventing domain intermingling and ensuring effective insulation of transcriptional programs.

Furthermore, we used a polymer model of chromatin chains to analyze the effects of cohesin depletion on chromatin domain conformation. The model has 1-kb resolution and consists of two domains, each 200 kb long, connected by a short linker (Fig. 8b,c (control in c)). In the simulations, domains appeared more mixed when cohesin was depleted, as quantified by the mixing score (see Methods; Fig. 8c,d; Movie S9). The polymer simulations also showed increased bead MSD with cohesin depletion, consistent with the experimental data (Fig. S10b–e). These results suggest that local domain mixing upon cohesin depletion could weaken insulation between neighboring transcriptional environments.

Because acute cohesin loss often produces only modest changes in bulk gene expression ^87^ and in ATAC-seq signals in HCT116 cells (Fig. S10f), we reasoned that transcriptional bursting— particularly its cis-coordination between loci—might provide a more sensitive functional readout. We therefore quantified this *cis*-coordination of transcriptional bursting in HCT116 using intron-seqFISH (intron sequential fluorescence *in situ* hybridization) (Figs. 8e,f; S10g,h) ^89,90^. We selected seven genes on Chromosome 1 whose per-gene burst frequency changed minimally upon RAD21 depletion (Fig. 8g,h; see Methods for details), enabling us to assess changes in co-bursting independently of large shifts in single-gene activity. Notably, the co-burst frequency of gene pairs significantly increased upon cohesin depletion (Fig. 8i,j), suggesting that cohesin helps prevent gene co-bursting along the chromosome, consistent with a previous study in mouse ES cells ^19^. Thus, cohesin depletion increases burst coupling between genes even when per-gene bursting remains largely unchanged, plausibly via local domain mixing (Fig. 8k).

## Discussion

In this study, we employed a combined approach of single-nucleosome imaging/tracking and super-resolution imaging to explore how cohesin regulates euchromatic chromatin organization in living human cells. Our approach provides a more comprehensive view of both global and local changes in chromatin dynamics and organization than previous studies that primarily focused on specific chromatin loci ^40,41^. Our results show that cohesin constrains local chromatin motion mainly through loop formation, particularly in euchromatin, rather than through sister chromatid cohesion (Fig. 2). Only minimal constraint effects arise from cohesion sites (Fig. 2), presumably because only a limited number of sister regions are aligned in human interphase cells ^91,92^. These findings align with previous studies reporting increased chromatin motion upon cohesin depletion ^5,40,41^. Furthermore, our results using head-tethered cohesin suggest that cohesin constrains euchromatin loop domains without requiring active loop extrusion (Fig. 3g–i).

To label euchromatin specifically, we developed euchromatin-specific labeling using histone H3.3-Halo (Fig. 4). The labeled genomic regions correspond to the Hi-C A compartment ^15^ with active histone marks in our cells (Fig. 4). Combining this euchromatin labeling with 3D-SIM ^7,45,46^ and STORM ^76,77^ enabled super-resolution imaging of euchromatic domains in live and fixed HCT116 cells, revealing their physically condensed nature. Nucleosome–nucleosome interactions and cohesin seem to drive the formation of condensed euchromatic domains.

Notably, a similar principle applies to condensed mitotic chromosomes, which are organized by condensins and nucleosome–nucleosome interactions^85,86^. Our findings provide critical evidence that challenges the established notion of euchromatin as largely open.

The physically condensed euchromatic domains can provide higher-order regulation of various DNA transactions, including transcription, whereas extended fiber loops cannot (Fig. S8g,h). If loop domains were extended or open, unnecessary interactions between domains and misregulation of transcription could occur (Fig. S8g). Condensed domains, per se, help avoid transcriptional interference and ensure proper gene regulation (Fig. S8h). In addition, condensed euchromatic domains likely hinder the access of transcriptional protein complexes to their target sequences, such as promoters or enhancers located in the inner core of chromatin domains ^7^.

These findings support the notion of higher-order gene regulation within a domain, such as a “buoy” model ^22,93^. Partial decondensation of such condensed domains through histone modifications or nucleosome eviction may also serve as a regulatory mechanism to activate gene transcription ^94^. Furthermore, condensed domains are more resistant to radiation than decondensed forms, presumably because condensed chromatin generates lower levels of reactive radicals ^95,96^. This feature is especially important for maintaining the integrity of actively transcribed euchromatic regions, which are directly connected to essential genome functions.

Together, these three features highlight the biological relevance of condensed euchromatic chromatin. While previous studies suggested that cohesin is involved in long-distance promoter–enhancer interactions, particularly those spanning domain boundaries ^97,98^, our findings highlight a previously underappreciated role of cohesin in constraining the conformational freedom of condensed euchromatic domains (Fig. 8k). We revealed how these physical properties of euchromatin domains are regulated by cohesin, and how they are coupled to genome organization and function. Super-resolution imaging of euchromatic domains and two-point MSD measurements between neighboring euchromatic nucleosomes show that, while cohesin depletion does not alter the overall compaction level of euchromatin domains, it increases nucleosome fluidity within these domains (Fig. 7i), leading to mixing of chromatin domains (Fig. 8b-d,k). Indeed, Hi-C contact probability plot in HCT116 cells ^87^ shows that cohesin loss decreases short-range contacts (∼0.1–1.5 Mb, TAD scale) and increases long-range contacts (Mb scale) (Fig. 8a). Co-activation of gene transcription was also observed upon cohesin loss (Fig. 8i,j) ^19^. Consistently, it was reported that cohesin loss increases “entropy” in genome folding and variation in gene expression ^88^. Together, domain mixing upon cohesin loss is essentially another readout of weakened insulation between domains. Condensed euchromatic domains constrained by cohesin ensure effective insulation ^50,51^ of transcriptional programs between domains.

### Limitations of the study

Our experiments were performed in somatic human cells (HCT116 and HeLa), where H3.3 incorporation predominantly marks euchromatin. H3.3 has also been reported in the telomeric region ^48^, or in facultative heterochromatin in specific contexts, such as embryonic stem cells and during cell-fate transitions ^99,100^. The degree of condensation may also differ across cell types. Exploring these contexts will be an interesting next step.

All SIM and STORM analyses were performed on asynchronous populations of HCT116 cells. While cohesin localization patterns and H3.3 distribution did not show noticeable variation across interphase cells, this remains an interesting question for future investigation, as genomic DNA content doubles and nuclear volume increases during cell-cycle progression ^43^.

## Methods

### Establishment of stable cell lines

HeLa S3 cells ^101^ and HT1080 cells with *lacO*/ EGFP-LacI and *tetO*/TetR-4×mCherry (a clone of TT75, TT165, kindly provided by Dr. T. Tanaka, University of Dundee, UK) ^60^ were cultured at 37 °C in 5% CO₂ in DMEM (D5796-500ML; Sigma-Aldrich) supplemented with 10% FBS (FB-1061/500; Biosera). All HCT116 cells (CCL-247; ATCC) with AID2 for rapid depletion ^53^ were cultured at 37 °C in 5% CO₂ in McCoy’s 5A medium (SH30200.01; HyClone) supplemented with 10% FBS.

For the establishment of HeLa cells stably expressing H2B-HaloTag or H3.3-HaloTag, the Flp-In system (V602020; Invitrogen) was used. pFRT-bla was first transfected into HeLa cells to integrate into the genome using the Effectene transfection reagent kit (301425; QIAGEN). Cells containing the FRT site were selected with 5 µg/mL blasticidin S (029-18701; Wako) and used for isolation of stable transformants. The isolation procedure using the Flp-In system was as previously described ^102^. pEF1-H2B-Halo-FRT or pEF1-H3.3-Halo-FRT was then transfected into HeLa cells harboring an FRT site, and transformants were selected with 200 µg/mL hygromycin B (10687-010; Invitrogen).

To stably express H2B-Halo or H3.3-Halo in the HCT116 cell line, the PiggyBac transposon system was used. The constructed plasmid pPB-CAG-IB-H2B-HaloTag or pPB-EF1α-H3.3-HaloTag was co-transfected with pCMV-hyPBase (provided by the Sanger Institute under a materials transfer agreement) into HCT116 cells using the Effectene transfection reagent kit. Transfected cells were selected with 10 µg/mL blasticidin S. HCT116 cells expressing OsTIR1(F74G) and endogenously tagged RAD21-mAID-mClover, CTCF-mAID-mClover, or WAPL-mAID-mClover ^53,103^ were used as parental cells. To stably express H2B-Halo in the HCT116 cell line with the head-tethering cohesin system ^69^, pPB-CAG-IB-H2B-HaloTag was co-transfected with pCMV-hyPBase into HCT116 cells using the Effectene transfection reagent kit. In this system, endogenous SMC1 is tagged with FK506-binding protein (FKBP) and endogenous SMC3 is tagged with a mutated FKBP–rapamycin-binding domain (FRB*).

Addition of the non-toxic rapamycin analog C16-(S)-7-methylindolerapamycin (AP21967/C16-AiRap), which selectively binds FRB* ^70,71^, induces forced head domain closure of cohesin (Fig. 3i). Cells expressing H2B-Halo were sorted based on fluorescence, as described in ^28^. Two weeks after transfection, cells expressing H2B-Halo were labeled with 50 nM HaloTag TMR ligand overnight at 37 °C in 5% CO₂. HaloTag TMR-positive cells were collected by flow cytometry (SH800S, Sony Biotechnology). The collected cells were subcloned, and clones stably expressing H2B-HaloTag were selected.

### Expression and localization of H2B-HaloTag or H3.3-HaloTag

To examine the expression level of H2B-HaloTag or H3.3-HaloTag, transfected cells were lysed in Laemmli sample buffer ^104^ supplemented with 10% 2-mercaptoethanol (133-1457; Wako) and incubated at 95 °C for 5 min to denature proteins. Cell lysates, equivalent to 1 × 10⁵ cells per well, were subjected to 12.5% SDS–PAGE and subsequent western blotting. For western blotting, the fractionated proteins were transferred to a PVDF membrane (IPVH00010; Millipore) using a semi-dry blotter (BE-320; BIO CRAFT). After blocking with 5% skim milk (190-12865; Fujifilm Wako), the membrane was probed with rabbit anti-H2B (1:10000; ab1790; Abcam), rabbit anti-H3 (1:100000; ab1791; Abcam), or mouse anti-HaloTag (1:1000; G9211; Promega) antibodies, followed by horseradish peroxidase–conjugated secondary antibodies: anti-rabbit (1:5000; 170-6515; Bio-Rad) or anti-mouse (1:5000; 170-6516; Bio-Rad). Chemiluminescence was detected using WBKLS0100 (Millipore) and EZ-Capture MG (AE-9300H-CSP; ATTO).

To examine the cellular localization of the H2B-HaloTag or H3.3-HaloTag, cells grown on poly-L-lysine–coated coverslips (P1524-500MG; Sigma-Aldrich; C018001; Matsunami) were treated with 5 nM HaloTag TMR ligand (8251; Promega) overnight at 37 °C in 5% CO₂. The cells were fixed with 1.85% formaldehyde (064-00406; Wako) at room temperature for 15 min, permeabilized with 0.5% Triton X-100 (T-9284; Sigma-Aldrich) for 15 min, and stained with 0.5 μg/mL DAPI (10236276001; Roche) for 5 min. Samples were mounted with PPDI (20 mM HEPES [pH 7.4], 1 mM MgCl₂, 100 mM KCl, 78% glycerol, and 1 mg/mL para-phenylenediamine [695106-1G; Sigma-Aldrich]) ^101^. Z-stack images (0.2 µm steps, 30 sections) were acquired using a DeltaVision Ultra microscope (Applied Precision) equipped with an Olympus PlanApoN 60× objective (NA 1.42), an sCMOS camera, an InsightSSI light source (∼50 mW), and a four-color standard filter set. Generally, H2B-Halo shows genome-wide localization, due to the frequent replacement of histone H2B within a few hours ^105^, while H3.3-Halo shows euchromatic localization ^47–49^.

### Biochemical fractionation of nuclei from cells expressing H2B-HaloTag or H3.3-HaloTag

Nuclei were isolated from HeLa S3 or HCT116 cells expressing H2B-HaloTag or H3.3-HaloTag as described previously ^106–108^. Briefly, collected cells were suspended in nuclei isolation buffer (3.75 mM Tris-HCl [pH 7.5], 20 mM KCl, 0.5 mM EDTA, 0.05 mM spermine, 0.125 mM spermidine, 1 μg/mL Aprotinin [T010A; TaKaRa], and 0.1 mM PMSF [P7626-1G; Sigma-Aldrich]) and centrifuged at 1,936 × *g* for 7 min at room temperature. The cell pellets were resuspended in nuclei isolation buffer and again centrifuged at 1,936 × *g* for 7 min at room temperature. Subsequent steps were performed at 4 ℃, unless otherwise noted. Cell pellets were resuspended in nuclei isolation buffer containing 0.025% Empigen (45165-50ML; Sigma-Aldrich; nuclei isolation buffer+) and homogenized immediately with 10 downward strokes of a tight Dounce-pestle (357546, Wheaton). The cell lysates were centrifuged at 4,336 × *g* for 5 min. Nuclear pellets were washed in nuclei isolation buffer+. The nuclei were incubated on ice for 15 min in the following buffers containing various concentrations of salt: HE (10 mM HEPES-NaOH [pH 7.5], 1 mM EDTA, and 0.1 mM PMSF), HE + 100 mM NaCl, HE + 500 mM NaCl,

HE + 1 M NaCl, and HE + 2 M NaCl. After each buffer incubation with increasing concentrations of salt, centrifugation was performed to separate the nuclear suspensions into supernatant and pellet fractions. The proteins in the supernatant fractions were precipitated by using 17% trichloroacetic acid (208-08081; Wako) and cold acetone. Both pellets were suspended in a Laemmli sample buffer and subjected to 12.5% SDS-PAGE, followed by Coomassie brilliant blue (031-17922; Wako) staining and western blotting using rabbit anti–H2B (1:10000; ab1790; Abcam), rabbit anti-H3 (1:100,000; ab1791; Abcam) or mouse anti–HaloTag (1:1000; G9211; Promega) antibodies, followed by the appropriate secondary antibody: anti-rabbit (1:5000 dilution; 170-6515; Bio-Rad) or anti-mouse (1:5000 dilution; 170-6516; Bio-Rad) horseradish peroxidase-conjugated goat antibody.

### Target protein depletion by AID2

To rapidly degrade RAD21-mAID-mClover, CTCF-mAID-mClover, and WAPL-mAID-mClover, HCT116 cells expressing OsTIR1(F74G) were treated with 1 µM 5Ph-IAA for 1 h (RAD21), 2 h (CTCF), or 4 h (WAPL), respectively, except where noted otherwise ^53,103^. Control cells were treated with 0.1% dimethyl sulfoxide (DMSO; D2650-5X5ML; Sigma-Aldrich).

Degradation of the target proteins was confirmed by the loss of mClover fluorescence. After treatment, cells were fixed, permeabilized, and stained with DAPI as described in the “Expression and localization of H2B-HaloTag or H3.3-HaloTag in HeLa cells” section. Z-stack images (30 sections at 0.2 µm intervals along the z-axis) were acquired using a DeltaVision Ultra microscope (Applied Precision) equipped with an Olympus PlanApoN 60× objective lens (NA 1.42). Nuclear mClover fluorescence intensity was quantified using Fiji, with background signal subtracted.

RAD21-mAC, WAPL-mAC, or CTCF-mAC degradation was also confirmed by western blotting. The procedure was the same as described in the “Expression and localization of H2B-HaloTag or H3.3-HaloTag in HeLa cells” section. Membranes were probed with mouse anti-RAD21 (1:1000; 05-908, Upstate), rabbit anti-WAPL (1:1000; A301-779A-T), rabbit anti-CTCF (1:1000; 10915-1-AP; PGI Proteintech), and mouse anti-mAID (1:1000; M214-3; MBL) antibodies, followed by horseradish peroxidase–conjugated goat anti-mouse secondary antibody (1:5000; 170-6516; Bio-Rad), or anti-rabbit IgG DyLight 800 (1:5000; SA5-35571; Invitrogen). As a loading control, a goat anti-GAPDH antibody (1:2500; Bio-rad; 12004158; StarBright Blue700 conjugated) was used.

### Single-nucleosome imaging

Established cell lines were cultured on poly-L-lysine–coated glass-based dishes (3970-035; Iwaki). H2B-Halo or H3.3-Halo incorporated into nucleosomes was fluorescently labeled with 80 pM HaloTag TMR ligand for 20 min at 37 °C in 5% CO₂, washed three times with 1× HBSS (H1387; Sigma-Aldrich), and then incubated in the following media overnight before single-nucleosome imaging. HeLa S3 cells were observed in DMEM (21063-029; Thermo Fisher Scientific), and HCT116 cells in McCoy’s 5A (1-18F23-1; BioConcept). These media were phenol red–free and supplemented with 10% FBS. To increase the number of tracked nucleosomes when applying the RL algorithm for motion classification, H2B-Halo was labeled with 50 nM PA-JF646 (provided by the Lavis Lab, Janelia Research Campus, VA, USA) ^73^ overnight using the same labeling procedure. A live-cell chamber (INU-TIZ-F1; Tokai Hit) and digital gas mixer (GM-8000; Tokai Hit) were used to maintain cell culture conditions (37 °C, 5% CO₂, and humidity) during microscopy.

Single nucleosomes were observed using an inverted Nikon Eclipse Ti microscope equipped with a 100-mW Sapphire 561-nm laser (Coherent) and an sCMOS ORCA-Flash 4.0 or ORCA-Fusion BT camera (Hamamatsu Photonics). Live cells labeled with TMR were excited with the 561-nm laser through an objective lens (100× PlanApo TIRF, NA 1.49; Nikon) and detected at 575–710 nm. An oblique illumination system with a TIRF unit (Nikon) was used to excite fluorescent nucleosome molecules within a thin area of the cell nucleus and reduce background noise (Fig. 1a). Sequential image frames were acquired using NIS-Elements (Nikon) at a frame rate of 50 ms under continuous illumination. Note that freely diffusing, non-nucleosomal histones cannot be tracked at this frame rate. For PA-JF646–labeled nucleosome tracking, the cells were continuously photoactivated with weak 405-nm illumination (1.0 mW, 25% AOTF attenuation) together with 640-nm laser excitation.

To visualize the basal nuclear surface (nuclear periphery), we adjusted the angle of laser illumination to efficiently capture a nuclear membrane marker, NUP107-Venus^109^. Almost uniform distributions of NUP107-Venus were observed in the nuclear periphery condition, while the nuclear interior condition showed a rim of NUP107-Venus ^110^. As reported previously ^110,111^, nucleosome dynamics on the nuclear surface were lower than those of the nuclear interior, which contained more euchromatin regions. Position determination accuracy is 7.3 nm (Fig. S7g).

### Single-nucleosome tracking analysis

To study nucleosome motion within chromatin domains accurately, we mainly focused on the 0–0.5 s time window, which corresponds to the spatial range of typical chromatin domain sizes (up to ∼300 nm). At longer time scales, other factors, such as higher-order structures (e.g., compartments, territories) and nuclear movements, become more influential (for details, see ^43,112^). To obtain the *Rc* value (radius of constraint) ^55^, we also analyzed longer-time data, up to ∼3 s, in some cases.

Image processing, single-nucleosome tracking, and single-nucleosome movement analysis were performed as previously described ^5,39,43,110^. Briefly, sequential images were converted to a 16-bit grayscale, and background noise was subtracted using the rolling-ball background subtraction (radius, 50 pixels) in ImageJ. Nuclear regions in the images were manually extracted. The fluorescent dots were fitted with a 2D Gaussian function ^76,113^ and tracked using u-track software^114^, or their centers were determined by LoG detector in Fiji plugin Trackmate ^115^.

To assess positional accuracy, we calculated the standard deviation of the two-dimensional movement of immobilized nucleosomes per 50 ms in FA-fixed HeLa S3 cells (*N* = 10 nucleosomes). We found that 12.5 nm (the mean of SDx and SDy) was the localization accuracy (also see Fig. S1d). Single-step photobleaching profile confirmed that the individual dots represent single nucleosomes (Fig. 1c).

For single-nucleosome imaging/tracking, we calculated displacement and mean squared displacement (MSD) of nucleosomes, because MSD captures the relevant constrained nucleosome dynamics, for the following reasons: the vast majority of the labeled histones are incorporated into nucleosomes constrained along a very long polymer rather than freely diffusing, and state mixing is minimal. In this respect, single-nucleosome imaging is very different from single-molecule tracking (SMT) of transcription factors (TFs). MSD in SMT of TFs can be error-prone because multiple diffusion states (free 3D diffusion, 1D sliding, specific binding) interconvert, and displacement-based analyses are often preferable (e.g., ^116^).

For single-nucleosome movement analysis, the displacement and MSD of the fluorescent dots were calculated based on their trajectory using a Python script. In our tracking, trajectories of single nucleosome dots on the XY plane (*x*_*i*_)^*N*^_i=0_ were acquired, where *x*_*i*_ indicates XY-coordinates at the time point *i*. Then, we calculated the MSD for the lag time Δ by:

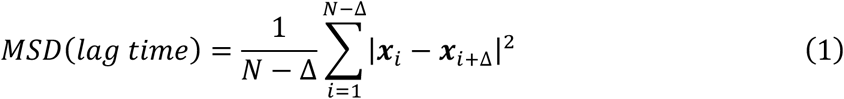

The originally calculated MSD was in 2D. To obtain the 3D value, the 2D value was multiplied by 1.5 (4 to 6 Dt). The calculated MSDs were fitted to a subdiffusive model *MSD*(*t*) = 6*D* · *t*^α^, where α is an anomalous diffusion exponent (0 < α < 1). Statistical analyses of the obtained single-nucleosome MSD captured through each histone protein were performed using Python.

The anomalous exponents (α) and the coefficient of subdiffusion *D*_sub_ were obtained from the slope and the intercept of the log–log MSD plots by linear regression within the 0–0.5 s time window, respectively. Graphs and statistical analyses of single-nucleosome MSD under different conditions were performed using R. For Spot-On analysis, single nucleosome trajectories data taken with 10 ms/frame were analyzed. The 3-state model is used for model fitting.

### Angle-distribution analysis

For the tracked consecutive points {(*x*_0_, *y*_0_), (*x*_1_, *y*_1_), ⋯, (*x*_*n*_, *y*_*n*_), ⋯ } of a single nucleosome on the XY plane, we converted the data into a set of displacement vectors, Δ*r*_*n*_ = (*x*_*n*+1_ − *x*_*n*_, *y*_*n*+1_ − *y*_*n*_)^*t*^. Then, we calculated the angle between two vectors Δ*r*_*n*_ and Δ*r*_*n*+1_. This procedure was carried out for all points in each trajectory. Finally, we plotted the normalized polar histogram by our Python program. The angle distribution was normalized by 2π, and the values correspond to the probability density. The colors indicate the angles. Asymmetric coefficient (AC) was calculated as indicated in Fig. S1h.

### Verification of DNA damage

To verify the level of DNA damage caused by rapid RAD21 depletion, the γH2AX immunostaining signal was quantified. HCT116 RAD21-mAID-mClover cells were treated with 1 µM 5Ph-IAA or 0.1% DMSO. As a positive control, cells were treated with 10 µg/mL bleomycin (4234400D4032; Nippon Kayaku) for 1 h. Immunostaining was performed as follows, with all steps carried out at room temperature. Cells grown on coverslips were fixed and permeabilized as described in the section “Expression and localization of H2B-HaloTag or H3.3-HaloTag.” After washing twice with HMK (20 mM HEPES [pH 7.5], 1 mM MgCl₂, and 100 mM KCl) for 5 min, the cells were incubated with 10% normal goat serum (NGS; 143-06561; Wako) in HMK for 30 min. The cells were then incubated for 1 h with rabbit anti-γH2AX (phospho S139) (1:1000, ab2893; Abcam) primary antibody diluted in 1% NGS in HMK. After four washes with HMK, the cells were incubated for 1 h with goat anti–rabbit IgG Alexa Fluor 594 (1:1000, A11037; Invitrogen) secondary antibody diluted in 1% NGS in HMK, followed by four washes with HMK.

DNA staining and mounting were performed as described in the section “Expression and localization of H2B-HaloTag or H3.3-HaloTag.” Optical section images were acquired at a 0.2 µm step size using a DeltaVision microscope (Applied Precision) as described in the same section. The number of γH2AX foci per nucleus was automatically quantified using Fiji. A fixed threshold value, determined by the Otsu method, was applied to segment the foci, which were then counted within each nucleus.

### Chemical treatment in single-nucleosome imaging

For chemical fixation, cells grown on poly-L-lysine coated glass-based dishes were incubated in 2% formaldehyde (diluted pure 16% formaldehyde) in 1 × PBS at room temperature for 15 min and washed with 1 × PBS. For head-tethering of the cohesin experiment, cells were treated with 1.5 μM AP21967 (A/C Heterodimerizer; 635057; Clontech) for 2 h. For RAD21 depletion in the head-tetherable cohesin cell line, cells were treated with 1 µg/mL doxycycline (631311; Clontech) overnight, followed by 500 µM indole-3-acetic acid (IAA; a natural auxin, 19119-61; Nacalai) for 1 h. Subsequently, 1.5 µM AP21967 was added for 2 h in the continued presence of both doxycycline and IAA (total IAA exposure: 3 h). Control cells were treated with 0.1% MQ (solvent used to dissolve doxycycline), 0.3% EtOH (for AP21967), and 0.1% DMSO (for IAA). RAD21 depletion was confirmed by the loss of mClover fluorescence during single-nucleosome imaging, and only cells with confirmed depletion were used for analysis.

### RNA interference

siRNA reverse transfections into HeLa, HCT116, and HT1080 cells grown on poly-L-lysine–coated glass-based dishes were performed using Lipofectamine RNAiMAX (13778-075; Invitrogen) according to the manufacturer’s instructions. The medium was replaced with fresh medium 16 h after transfection. The transfected cells were used for subsequent experiments 48 h (RAD21, CTCF) or 72 h (WAPL, Sororin) after transfection. As a control, an oligonucleotide (4390843; Ambion; sequence undisclosed) was used. The siRNA oligo sequences and concentrations used were as follows: RAD21 (60 nM, 5’-CAGCUUGAAUCAGAGUAGAGUGGAAdTdT-3’), CTCF (60 nM, 5’-GCGCUCUAAGAAAGAAGAUUCCUCUdTdT-3’), WAPL (80 nM, 5’-GGUUAAGUGUUCCUCUUAUdTdT-3’), Sororin (60 nM, 5’-GCCUAGGUGUCCUUGAGCUdTdT-3’).

Knockdown of target proteins was confirmed by immunostaining, performed as described in the section “Verification of DNA damage.” Cells were incubated for 1 h with primary antibodies diluted in 1% NGS in HMK: mouse anti-RAD21 (1:1000; 05-908; Upstate), mouse anti-CTCF (1:3000; 07-729; Millipore), mouse anti-WAPL (1:20000; MABE1106; Millipore), or rabbit anti-Sororin (CDCA5) (1:2500; ab192237; Abcam). After four washes with HMK, cells were incubated for 1 h with secondary antibodies diluted in 1% NGS in HMK: goat anti–mouse IgG Alexa Fluor 488 (1:1000, A11029; Invitrogen), goat anti–mouse IgG Alexa Fluor 594 (1:1000, A11032; Invitrogen), goat anti–rabbit IgG Alexa Fluor 594 (1:1000, A11037; Invitrogen), or goat anti–rabbit IgG Alexa Fluor 647 (1:1000, A21245; Invitrogen).

Optical section images were acquired at a 0.2 µm step size using a DeltaVision microscope (Applied Precision) as described in the section “Expression and localization of H2B-HaloTag or H3.3-HaloTag in HeLa cells.” To quantify the knockdown effect, the mean nuclear signal intensities were calculated after background subtraction (signals outside nuclei) and plotted.

### Cell cycle synchronization

For synchronization of HCT116 cells treated with siSororin RNA in the late S–G2 phase, cells (1.0 × 10^5^ cells/dish) were plated in medium containing siSororin RNA. Cells were then treated with 9 µM RO3306 (CS-3790; Funakoshi) for 16 h before observation. For synchronization of HCT116 or HT1080 cells in the late S–G2 phase, cells were initially plated at a concentration of 1.0 × 10^5^ cells/mL, incubated for one day, and then treated with 9 µM RO3306 for 16 h.

### Chromosome spreads

To assess the formation of sister chromatid cohesion, mitotic chromosome spreads were prepared. Cells (treated with siControl or siSororin RNA) were incubated with 0.2 µg/mL colcemid (045-16963; Wako) for 1 h at 37 °C. The following steps were performed at room temperature, except for the final fixation step. After washing with PBS, cells were trypsinized and resuspended in hypotonic buffer (75 mM KCl) for 10 min. Cells were then gently fixed by repeating the following treatment three times: incubation in fixation buffer (methanol:acetic acid, 3:1) for 5 min, centrifugation at 850 × *g* for 4 min, and resuspension in fresh fixation buffer. Finally, the cells were completely fixed by incubation in 200 µL of fixation buffer at –30 °C for at least 30 min.

For imaging, 5 µL of the fixed cell suspension was dropped onto coverslips, mixed, and completely dried at 60 °C for 30 min. Dried cells were stained with 0.5 µg/mL DAPI for 5 min, followed by PPDI mounting. Images of mitotic chromosome spreads were obtained using a DeltaVision Ultra microscope with an Olympus PlanApoN 60× objective (NA 1.42) and an sCMOS camera. The number of chromosomes per cell was counted manually. We confirmed the loss of sister chromatid cohesion in Sororin-depleted cells: the percentage of separated or X-shaped sister chromatids increased (Fig. 2b), consistent with a previous report ^58^.

### *lacO*/EGFP-LacI and *tetO*/TetR-4×mCherry foci imaging and tracking

*lacO*/EGFP-LacI and *tetO*/TetR-4×mCherry foci in live HT1080 cells were imaged under the same conditions as described in “Single-nucleosome imaging,” except that a lower laser power was used due to the higher fluorescence intensity of the foci compared with TMR-labeled single nucleosomes. EGFP-LacI was excited with a 488 nm laser, and TetR-mCherry with a 561 nm laser. Imaging was performed sequentially, first acquiring the 561 nm channel, followed by the 488 nm channel. Movies were recorded at a frame rate of 50 ms per frame, with a total of 1200 frames (1 min). Cells were treated with siRNA (see “RNA interference”) and synchronized to the late S–G2 phase (see “Cell cycle synchronization”). The precise positions of the labeled genomic loci were identified and tracked using the TrackMate plugin ^115,117^. Mean squared displacement (MSD) of the foci was calculated from the trajectories using the same procedure described in “Single-nucleosome tracking analysis.” The *tetO* array, visualized with TetR-4×mCherry, more frequently appears as two separate dots compared to the *lacO* array, visualized with EGFP-LacI. This suggests that the *lacO* array is located near a cohesin site, which likely prevents the sister loci from separating ^60^. Consistently, ENCODE data ^61^ indicated that the *lacO* site lies within 5 kb of a cohesin-enriched region (e.g., SMC3 and RAD21; Fig. 2e), as shown in ^60^.

### Bayesian-based Richardson-Lucy algorithm (RL algorithm)

The RL algorithm is the iterative algorithm of Richardson ^118^ and Lucy ^119^ (RL) to derive smooth distributions from the noisy data used in image processing ^120,121^. With sufficient observation, the algorithm converts raw, noisy trajectories into a smooth distribution of MSD, providing a denoised, population-level view of heterogeneous nucleosome mobility.

For the convenience of the reader, we briefly explain the mathematical principles of the RL algorithm. A more detailed description has been reported previously ^72^.

Here, the average MSD of nucleosomes, *M*^-^ = 〈*M*_*i*_〉, where 〈⋯ 〉 is the average taken over *i* and along the observed trajectories.

The distribution of MSD is captured by

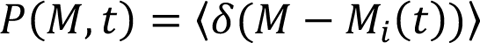

Due to the short lifetime of observed fluorescence of single nucleosomes, the individual nucleosome MSD data are insufficient to provide for a clear *P*(*M, t*). However, this problem is overcome by using the RL algorithm. From the observed data, we first calculate the self-part of the van Hove correlation function (vHC),

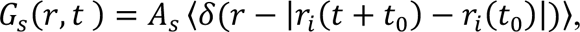

where *r*_*i*_ is the projected coordinate of the *i*th nucleosome on the 2-dimensional imaging plane and *A*_*s*_ is a constant to normalize *G*_*s*_ as ∫ *d*^2^ *rG*_*s*_(*r, t*) = 1. The calculated vHC is shown at *t* =0.5s. *G*_s_ is expanded in Gaussian bases, 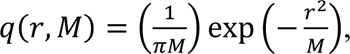, as

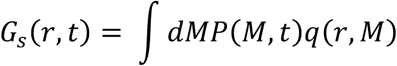

Given a noisy estimate of *G*_*s*_(*r, t*), *P*(*M, t*) is extracted as coefficients of expansion using the RL iterative scheme. For this, *P*(*M, t*) was calculated in an iterative way with the RL algorithm: starting from the initial distribution, 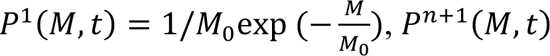 at the (n+1)-th iteration was obtained by 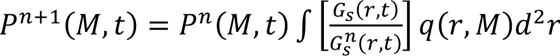, with *G*^*n*^(*r, t*) = ∫ *P*^*n*^ (*M, t*)*q*(*r, M*)*dM*. This equation was iterated under the constraints *P*^*n*+1^(*M, t*) > 0 and ∫ *P*^*n*+1^ (*M, t*)*dM* = 1.

The obtained distribution curves were fitted with multiple Gaussian functions, and the peak position and area of each Gaussian component were calculated. Subpopulation boundaries were estimated from the RL-curve: using the local minima for the “Super slow”–“Slow” and “Slow”– “Fast” transitions, and the midpoint between peaks for the “Fast”–“Super fast” transition. For MSD analysis, trajectories within each cell were categorized based on their MSD at Δt = 0.5 s, and mean squared displacement (MSD) was calculated separately for each subpopulation.

Centroids of the categorized trajectories were mapped to visualize their localization. It should be noted that reliable fitting of α values for the “Super slow” and “Slow” categories is difficult due to their limited mobility.

### H3.3-Halo labeled nucleosome pulldown, purification of nucleosomal DNA, and library preparation

Cell nuclei were isolated as previously described ^106–108^, with minor modifications. Briefly, HCT116 or HeLa cells were maintained at 37 °C and 5% CO₂ in McCoy’s 5A or DMEM supplemented with 10% FBS, respectively. Collected cells were washed with nuclei isolation buffer (3.75 mM Tris–HCl pH 7.5, 20 mM KCl, 0.5 mM EDTA, 0.05 mM spermine, 0.125 mM spermidine, 0.1% Trasylol (14716; Cayman Chemical), 0.1 mM PMSF (P7626-1G; Sigma-Aldrich)) with two cycles of centrifugation at 193 × *g* for 7 min at 23 °C. Pellets were resuspended in nuclei isolation buffer containing 0.05% Empigen BB detergent (45165-50ML; Sigma-Aldrich) (nuclei isolation buffer+) and homogenized with ten downward strokes using a tight Dounce pestle. After centrifugation at 440 × *g* for 5 min, the pellets were washed with nuclei isolation buffer+. This step was repeated four times before nucleosome purification.

Chromatin purification was performed as described by ^43,122^, with modifications. Nuclei (equivalent to ∼2 mg of DNA) in buffer (10 mM Tris–HCl pH 7.5, 1.5 mM MgCl₂, 1.0 mM CaCl₂, 0.25 M sucrose, 0.1 mM PMSF) were digested with 50 U micrococcal nuclease (MNase, LS004797; Worthington) at 30 °C for 1 h. During digestion, 1 µM HaloTag PEG-Biotin Ligand (G859A; Promega) or an equivalent volume of DMSO was added. After centrifugation at 440 × *g* for 5 min at 4 °C, nuclei were lysed in lysis buffer (10 mM Tris–HCl pH 8.0, 5 mM EDTA, 0.1 mM PMSF) on ice for 5 min. Lysates were dialyzed overnight at 4 °C against dialysis buffer (10 mM HEPES–NaOH pH 7.5, 0.1 mM EDTA, 0.1 mM PMSF) using Slide-A-Lyzer (66380; Thermo Fisher Scientific). After centrifugation at 20,400 × *g* for 10 min at 4 °C, the supernatant was recovered as the purified nucleosome fraction (mainly mononucleosomes).

To verify MNase digestion, DNA was purified, run on a 1.5% agarose gel, and stained with ethidium bromide. The biotinylated or untreated nucleosome fraction (28 µg) was diluted with an equal volume of 2× binding buffer (10 mM HEPES–NaOH pH 7.5, 200 mM NaCl, 2 mM DTT, and cOmplete EDTA-free protease inhibitor cocktail (11873580001; Roche)). As described ^123^, chromatin solution was added to streptavidin-FG beads (TAS8848 N1170; Tamagawa-Seiki) pre-equilibrated with coating buffer (PBS, 2.5% BSA, 0.05% Tween 20) for 1 h. Mixtures were incubated at 5 °C for 2 h with shaking. Beads were collected with a magnetic rack and washed as described ^124^. In brief cells were washed once with cold 1× binding buffer (10 mM HEPES– NaOH pH 7.5, 100 mM NaCl, 1 mM DTT), twice with cold wash buffer 1 (10 mM HEPES– NaOH pH 7.5, 140 mM NaCl, 1% Triton X-100, 0.5% NP-40 (11332473001; Roche), 0.1% SDS), twice with cold wash buffer 2 (10 mM HEPES–NaOH pH 7.5, 360 mM NaCl, 1% Triton X-100, 0.5% NP-40, 0.1% SDS), twice with cold wash buffer 3 (10 mM HEPES–NaOH pH 7.5, 250 mM LiCl, 0.5% Triton X-100, 0.5% NP-40), and once with TE buffer (10 mM Tris–HCl pH 8.0, 1 mM EDTA).

Beads were resuspended in 50 µL TE buffer. For DNA extraction, 40 µL of beads were treated with RNase A (1 mg/mL, 10109169001; Roche, 4 µL) for 30 min at 37 °C, followed by proteinase K digestion (20 mg/mL, 169-21041; FUJIFILM Wako, 4 µL) with 10 µL of 10% SDS at 50 °C for 1 h with shaking (1,200 rpm). After magnetic separation, DNA was extracted using AMPure XP beads (A63880; Beckman Coulter) and eluted in Milli-Q water. DNA concentration was quantified with the Qubit system (Q32851; Thermo Fisher Scientific), and quality was assessed by Agilent 2100 Bioanalyzer using a High Sensitivity DNA kit (5067-4626; Agilent). Libraries were prepared using the ThruPLEX DNA-seq Kit (R400675; Takara Bio) and checked with an Agilent 2100 Bioanalyzer. Pooled libraries were sequenced (paired-end, 2 × 100 bp) on the Illumina NovaSeq 6000 platform.

For protein analysis, the remaining 10 µL of beads were resuspended in an equal volume of 2× Laemmli sample buffer ^104^ containing 10% 2-mercaptoethanol (133-1457; FUJIFILM Wako) and incubated at 99 °C for 10 min. After centrifugation at 10000 × *g* for 3 min, input and pulldown samples (biotinylated and negative control) were separated by SDS–PAGE and analyzed by western blotting with anti-HaloTag antibody (G9211; Promega).

For data analysis of purified H3.3-Halo nucleosomal DNA, the nf-core ChIP-seq pipeline (nfcore/chipseq v2.1.0) ^125,126^ was used with Docker configuration and default parameters. The human genome hg19 (Illumina iGenomes) was used as a reference. Peaks were called using MACS2 (broad mode) ^127^. Overlap between H3.3-Halo regions and contact domains (Hi-C A compartments, histone modifications) was assessed against published datasets ^8,87,128^. ChIP-seq peaks for histone modifications in HCT116 cells were downloaded from 4D Nucleome data portal^129,130^: Boundaries (4DNFIBKY9EG9), Compartment (4DNFIZHT1Y8P), RAD21 (4DNFIM7KV9EA), CTCF (4DNFIT3E2YZZ), RNAP II (4DNFIFNJW3NE), H3K4me1 (4DNFI27LAZBR), H3K4me3 (4DNFIRPBX1BM), H3K27ac(4DNFIGINV1VI), H3K36me3(4DNFIPYVJMFK), H3K79me2 (4DNFI7O6ZSBK), H4K16Ac (4DNFI6DW6GVA), H3K27me3 (4DNFI2EWSBH4), H3K9me3 (4DNFIV229BKJ), H4K20me3 (4DNFIMYKFY9Y). ATAC-seq data was generated in this study. Hi-C domains from HeLa ^15^ were obtained from GEO (GSE63525). Compartment annotations were from SNIPER predictions ^131^. ChIP-seq peaks for histone modifications in HeLa cells were downloaded from ENCODE ^61,132^: H3K4me2 (ENCFF108DAJ), H3K4me3 (ENCFF447CLK), H3K9ac (ENCFF723WDR), H3K27ac (ENCFF144EOJ), H3K79me2 (ENCFF916VLX), H3K36me3 (ENCFF001SVY), H3K4me1 (ENCFF162RSB), and H3K27me3 (ENCFF252BLX).

### ATAC-seq experiments and data processing

ATAC-seq was performed as previously described with minor modification ^133^. Briefly, HCT116 cells treated with DMSO or 5Ph-IAA (1 µM, 3 h) were harvested by trypsin dissociation and 50,000 viable cells per sample were collected, washed once with ice-cold phosphate-buffered saline (PBS), and lysed in 50 μL ATAC-Resuspension Buffer (RSB) containing 0.1% NP40, 0.1% Tween-20, and 0.01% Digitonin. Lysed nuclei were washed with 1 mL of cold ATAC-RSB containing 0.1% Tween-20. Isolated nuclei were immediately subjected to transposition using Tn5 transposase described previously ^134^. Transposition reactions were carried out in a total volume of 50 μL containing 25 μL 2x Tagmentation buffer, 0.5 μL transposase (80nM final), 16.5 μL PBS, 0.5 μL 1% digitonin, 0.5 μL 10% Tween-20, 5 μL H_2_O. The tagmentation reaction was incubated at 37°C for 30 min with gentle mixing. Following transposition, DNA was purified using Zymo DNA Clean and Concentrator-5 Kit (cat# D4014) according to the manufacturer’s instructions.

Library amplification was performed using PCR with indexed primers described in ^133^ and NEBNext® High-Fidelity 2X PCR Master Mix (M0541L). The optimal number of amplification cycles was determined by quantitative PCR to minimize over-amplification. PCR was performed involving an initial 5-min extension at 72°C to fill gaps, followed by 8-14 cycles (98°C for 10s, 63°C for 30s, 72°C for 30s). Amplified libraries were purified using AMPure XP beads and assessed for quality and fragment size distribution using an TapeStation. Libraries exhibiting the characteristic nucleosomal laddering pattern were sequenced on an Illumina platform (NovaSeq X Plus) using paired-end 150bp reads.

Paired-end 150-bp reads were collected for each sample. Nextera Transposase adapter sequences were removed using Cutadapt (v4.2) ^135^. Trimmed reads were aligned to the human genome assembly hg38 using Bowtie2 (v2.5.2) ^136^ with the ‘--local’ and ‘--no-discordant’ options.

Alignments with mapping quality scores < 7 were removed using SAMtools (v1.18) ^137^, and only properly paired reads were retained. Open chromatin regions were identified using MACS2 (v2.2.7.1) ^138^ with options: -f BAMPE -g hs -q 0.0001. Peaks across different conditions were merged to generate consensus peaks and HTSeq-count (v2.0.2) ^139^ was used to calculate fragment counts for the consensus peaks. The fragment count matrix was further normalized using DESeq2 (v1.40.1) ^140^ size factors, and the correlation analyses were performed using normalized count matrix in R environment (v4.3.2). ATACseqQC package (v1.34.0) ^141^ was used to calculate transcription start site (TSS) enrichment scores. Fragment size distributions and duplicates levels for each sample were calculated from BAM files using GenomicAlignments (v1.36.0) ^142^ and GenomicRanges (v1.52.0) ^142^ packages within the R/Bioconductor environment.

### Sample preparation for SIM imaging

Two days before labeling, cells were grown on poly-L-lysine-coated, super-resolution–grade coverslips (No. 1S; 0.17 mm thickness; CS01803; Matsunami). H3.3-Halo in HCT116 cells was fluorescently labeled by incubating the cells for more than 1 h with 50 nM TMR. Cell fixation and immunostaining were performed as described in the section “Verification of DNA damage.” Fixed cells were observed at room temperature.

To experimentally “melt” the euchromatin domains, the cells were treated with trichostatin A (TSA; 203-17561, Wako) for 7 h before fixation. For the double treatment with TSA and acute cohesin depletion, cells were also treated with 1 µM 5Ph-IAA for the final 1 h.

For live imaging, cells were grown poly-L-lysine–coated glass-based dishes. H3.3-Halo incorporated into nucleosomes was fluorescently labeled with 50 nM HaloTag TMR ligand for 1 h at 37 °C in 5% CO₂, washed three times with 1× HBSS, and then observed in phenol red–free McCoy’s 5A at 37 °C in 5% CO₂.

For immunostaining, cells were incubated with diluted primary antibodies in 1% NGS in HMK for 1 h: mouse anti-RAD21 (1:1000; 05-908, Upstate), mouse anti-RNAP II CTD (1:1000; ab817, Abcam), mouse anti-RNAP II Ser5P (1:1000; provided by Kimura Lab, Science Tokyo, Japan), or mouse anti-RNAP II Ser2P (1:1000; provided by Kimura Lab, Science Tokyo, Japan) antibody. After four washes with HMK, cells were incubated with diluted secondary antibodies in 1% NGS in HMK for 1 h: goat anti–mouse IgG Alexa Fluor 488 (1:500; A11029, Invitrogen) or goat anti–mouse IgG Alexa Fluor 647 (1:500; A21236, Invitrogen), followed by four additional washes with HMK. DNA staining and mounting were performed as described in the section “Expression and localization of H2B-HaloTag or H3.3-HaloTag.”

### 3D-SIM microscopy

Structured illumination microscopy (SIM) was performed on a Nikon N-SIM S system (Ti-2 stand; Hamamatsu ORCA Fusion BT camera; Nikon Perfect Focus; Chroma ET525/50m, ET595/50m, and ET700/75m emission filters). A Nikon 100× PlanApo TIRF oil-immersion objective (NA 1.49) was used. NIS-Elements (Nikon) software was used to control the system and acquire 3D-SIM images. Illumination was provided by ZIVA light engines (Lumencor) containing 405 nm, 476 nm, 545 nm, and 637 nm lasers. Light was directed to the sample via a specific filter for each channel.

3D image stacks were acquired over the entire cell volume in z, with 15 raw images per plane (five phases and three angles). Optical sectioning images were acquired at 0.12 µm intervals, with a total of 51 sections covering the entire nucleus (6 µm in thickness). Images acquired on the Nikon N-SIM-S system were reconstructed and corrected for chromatic aberrations using stack reconstruction in the NIS-Elements software.

System performance and both raw and reconstructed data quality were carefully assessed and optimized using the SIMcheck ImageJ plugin (Fig. S8) ^75^. To exclude potential false-positive nuclear marker signals, we applied the MCNR map function of SIMcheck, which generates a metric of local stripe contrast across different regions of the raw data and directly correlates this with the level of high-frequency information content in the reconstructed data ^75^. Only immunofluorescent spot signals with MCNR values above a stringent quality threshold were considered, whereas localizations with low MCNR values were discarded as unreliable SIM signals.

### Classification analysis for 3D-SIM images

3D assessment of DAPI intensity classes or H3.3-Halo intensity classes as a proxy for chromatin compaction was performed as previously described ^143,144^. First, 1,000 was subtracted from all voxel intensity values to remove background noise, and nuclei were cropped. Nuclear voxels were automatically identified from the DAPI channel using Gaussian filtering and thresholding via the *dapimask* function in the *nucim* R package.

For chromatin quantification, a 3D nuclear mask was generated to define the region for segmenting DAPI or H3.3-Halo signals into seven intensity classes with equal variance, following the method of Schmid et al. ^144^ and Miron et al. ^7^. In brief, voxel-wise classification was performed using a hidden Markov random field model that combines a finite Gaussian mixture model with a spatial Potts model, implemented in R. This approach enables threshold-independent classification of chromatin compaction states at the voxel level based on DAPI or H3.3-Halo intensity. Heat maps of the seven intensity classes in individual nuclei were generated using Fiji and displayed in either color or grayscale. The relative volume of each class was calculated as the volume of that class divided by the total nuclear volume detected by DAPI staining.

### Power spectrum analysis for 3D-SIM images

Single mid-sections from individual nuclei were cropped from the original reconstructed 3D-SIM images. The cropped images were then converted to 2D Fourier spectra using Python ^145^.

The resulting power spectra were radially averaged to analyze the amplitude at specific spatial periodicities.

### STORM imaging

HCT116 RAD21-mAC H3.3-Halo cells were cultured on poly-L-lysine–coated glass-based dishes (3970-035; Iwaki). H3.3-Halo incorporated into nucleosomes was labeled with 100 nM HMSiR-Halo (A201-01, Goryo Chemical) in medium for 3 h at 37 °C in 5% CO₂. Cells were washed three times with phenol red–free McCoy’s 5A medium (PR-free McCoy’s 5A) with 10% FBS and then incubated with PR-free McCoy’s 5A with 10% FBS and 0.01% DMSO or 10% FBS and 1 µM 5Ph-IAA for 1 h. Cells were fixed with 3.7% formaldehyde (FA) (064-00406, Wako) in Opti-MEM (31985062, Gibco) at 37 °C in 5% CO₂ for 15 min, washed three times with PR-free McCoy’s 5A with 10% FBS, and observed in PR-free McCoy’s 5A with 10% FBS. Single molecules were observed using an inverted Nikon Eclipse Ti-2 microscope with an ILE laser unit (ANDOR) and the sCMOS ORCA-Fusion BT camera. Fluorescent molecules in living cells were excited by the 640-nm laser through an objective lens (100× Apo TIRF, NA 1.49; Nikon) and detected. An oblique illumination system with a TIRF unit (TI2-LA-TIRF-E, Nikon) was used to excite labeled molecules within a limited thin area in the cell nucleus and reduce the background noise. Sequential 5,000 images were acquired using NIS-Elements (Nikon solutions) at a frame rate of 10 ms under continuous illumination. To maintain cell culture conditions (37°C, 5% CO_2_, and humidity) under the microscope, a live cell chamber with a digital gas mixer and a warming box (Tokai Hit) was used.

STORM images were constructed using the Fiji package ThunderSTORM ^146^ based on sequential images of HMSiR. For clustering analysis, TrackMate was also used for localization. The estimated localization accuracy is 4.546 nm for x and 4.672 nm for y (Fig. S7g).

To estimate the individual cluster size, we used the SR-Tesseler-DBSCAN pipeline ^78^. First, SR-Tesseler is used with a low density factor (0.7) to extract chromatin regions, then DBSCAN (with 90 nm distance and 100 minimum samples) detects each cluster.

### Identification of broad H3.3-enriched regions

Broad H3.3-enriched regions were called using epic2 ^147^. Replicate BAM files (Rep1 and Rep2) were processed independently with a 5-kb bin size, allowing up to two consecutive gaps, and using an FDR cutoff of 0.05. To derive a reproducible set of broad H3.3 domains, the two replicate callsets were first combined and overlapping intervals were merged to generate a union set. Subsequently, intervals that reciprocally overlapped by at least 30% of their length between Rep1 and Rep2 were retained, and the supported intervals from both replicates were merged to obtain the final intersection set. The resulting reproducible H3.3 domain set comprised 1,553 intervals with a median length of ∼210 kb.

### Peak sets used for distance-to-boundary analyses

Marker peak sets included ChIP-seq peaks for H3K27Ac, H3K4me1, H3K4me3, H3K36me3, H3K27me3, H3K9me3, H4K16Ac, H4K20me3, and H3K79me2, as well as ChIP-seq peaks for RNAP II, RAD21, and CTCF, and ATAC-seq peaks. Hi-C boundary annotations were obtained from in situ Hi-C ^87^. RNAP II/RAD21/CTCF ChIP-seq datasets were taken from ^128^, while ATAC-seq was generated in this study.

### Signed distance of peaks to H3.3 domain boundaries

H3.3 domains were represented as intervals (*chr, s*_*i*_, *e* _*i*_) and each marker peak (*chr, a* _j_, *b* _j_) was represented by its midpoint *m* _j_ = (*a* _j_ + *b* _j_)/2. For each midpoint, the closest H3.3 domain on the same chromosome was defined as the one minimizing the distance to either boundary (start or end). A signed distance *d*(bp) was computed as the minimum distance to the nearest boundary, assigned positive if the midpoint lies inside the domain (*s*_*i*_ ≤ *m* _j_ < *e* _*i*_) and negative if it lies outside (*m* _j_ < *s*_*i*_ or *m* _j_ ≥ *e* _*i*_). Thus, *d* = 0 corresponds to the H3.3 boundary, negative values indicate outside positions, and positive values indicate inside positions, enabling direct comparison of distances on the same bp scale.

### Dual-color labeling and imaging

HCT116 RAD21-mAID-mClover H3.3-Halo cells were cultured on poly-L-lysine–coated glass-based dishes (3970-035; Iwaki). H3.3-Halo incorporated into nucleosomes was sequentially labeled, first with 200 nM PA-JF549 ^73^ for 20 min and then with 50 nM PA-JF646 ^73^ for 20 min at 37 °C in 5% CO₂. Cells were washed three times with 1× HBSS (H1387; Sigma-Aldrich) and incubated in phenol red–free McCoy’s 5A medium for more than 1 h before single-nucleosome imaging (Fig. S9).

Simultaneous dual-color imaging in live HCT116 cells was performed under the same conditions as described in “Single-nucleosome imaging,” except that a beam splitter (W-VIEW GEMINI; Hamamatsu Photonics) was used, equipped with an image-splitting dichroic beamsplitter (FF640-FDi01-25×36; Hamamatsu Photonics) and bandpass filters (FF01-593/40-25 and FF01-676/29-25; Hamamatsu Photonics). The cells were continuously photoactivated with weak 405 nm illumination (1.0 mW, 25% AOTF attenuation).

### Two-point MSD analysis

In our single-nucleosome tracking, trajectories of PA-JF549 and PA-JF646 labeled nucleosomes on the XY-plane, 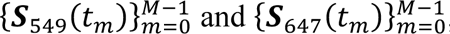, were simultaneously acquired, with a time interval of Δ*t* = 0.05 s and *t*_*m*_ = *m* Δ*t* (*m* = 0, 1, 2, ⋯, *M* − 1). To evaluate dynamic fluctuations between two points, we considered the relative vector between two trajectories, *Q* (*t*_*m*_) = *S* _549_(*t*_*m*_) − *S* _647_(*t*_*m*_). Then, we calculated the two-point MSD for the lag time

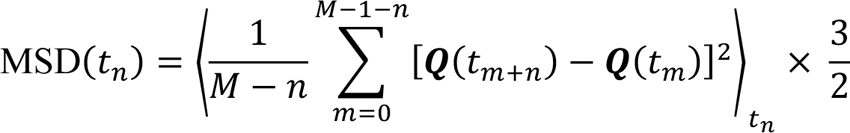

where ⟨⋅⟩_*tn*_ represents the ensemble average for trajectories at the lag time *t*_*n*_ and the coefficient 3/2 is a correction factor for conversion from 2-dimensional to 3-dimensional values.

### Coarse-grained polymer model

We simulate chromatin movement by using a bead-and-spring model of polymer chains. This coarse-grained polymer model was used to analyze how cohesin-mediated constraints influence chromatin movement during short microscopic observation times. In this model, the position of each bead is represented by *r*_*i*_ for *i* = 1, ⋯, 2*n* + 5, where *n* = 200. Each bead corresponds to a coarse-grained 1-kb chromatin region, and the chain consists of two 200-kb domains (*i* = 1, ⋯, *n* and *i* = *n* + 6, ⋯, 2*n* + 5), connected by a linker (*i* = *n* + 1, ⋯, *n* + 5). The temporal change of these positions is governed by the Brownian dynamics,

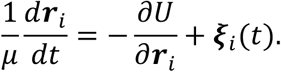

Here, ξ_*i*_ is a Gaussian random force, satisfying 〈ξ_*i*_(*t*)〉 = 0 and 〈ξ_*i*α_(*t*)ξ_jβ_(*t*^′^)〉 = (2*k*_B_*T*/μ)δ(*t* − *t*^′^)δ_*i*j_δ_αβ_, where *k*_B_ is the Boltzmann constant, *T* is temperature, and α, β = *x, y*, or *z*. The potential energy *U* consists of the following components,

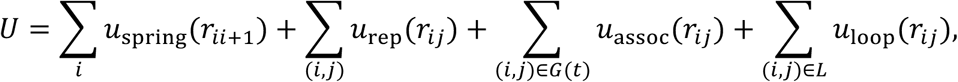

with *r*_*i*j_ = {*r*_*i*_ − *r*_j_}. Here, 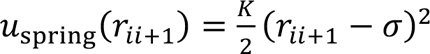 represents the spring that connects the *i*th and *i* + 1th beads. 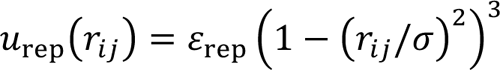 accounts for the soft, repulsive interaction between a pair of beads (*i, j*), with the sum taken over all pairs within the distance σ. 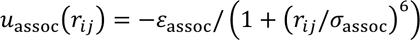 is the attractive interaction between beads (*i, j*) listed in a set 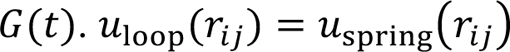 represents the constraint imposed by cohesin, which connects beads (*i, j*) in a set *L*.

In the present simulation, attractive interactions between beads drive the condensation of chromatin, leading to the formation of cluster-like domains of beads, as observed in microscopy. These attractive interactions should depend on the structural characteristics of the nucleosome cluster represented by each bead, which vary stochastically. To account for this variability, we update the set *G*(*t*) stochastically at each simulation step of the Brownian dynamics. To prevent instability caused by excessively dense attractive interactions, we introduce a valency parameter *z*, which represents the maximum number of attractive partner beads for each bead ^148^. At each simulation step in the Brownian dynamics, we randomly select a pair (*i, j*) from those not already included in *G*(*t*) and whose mutual distance *r*_*i*j_ is less than σ_assoc_. We then add the selected pair to the set *G*(*t*) at a rate *k*_assoc_ if the number of partners for either bead *i* or *j* does not exceed *z*. Conversely, a pair that is already included in *G*(*t*) is removed from *G*(*t*) at a rate *k*_dissoc_. We consider nucleosomes in the linker chain to be either acetylated or depleted and refer to such an area as the acetylated region. We assume that a fraction Φ of the chain within each domain also consists of acetylated regions. To determine the distribution pattern of the acetylated regions with the fraction Φ and the average region size *w*, we generate a Markov chain of transitions between acetylated and deacetylated states along the chromatin chain. The distribution pattern is rejected and re-generated until the fraction of acetylated regions coincide with φ. We assign the value *z* = 0 to the linker region, *z* = 1 to the acetylated regions within each domain, and *z* = 3 to the deacetylated regions. Thus, acetylated regions were modeled as having reduced effective attractive capacity compared to deacetylated regions, consistent with weaker compaction propensity of acetylated chromatin.

The temporal variation of *L* depends on how cohesin interacts with the chromatin chain. In this study, we do not make specific assumptions about these interactions; instead, we focus on the constraints that cohesin imposes during the observed short period (∼0.5 s) of nucleosome movement. Therefore, we assume that *L* remains fixed for each chain we simulate. Cohesin is expected to bind more frequently in regions where transcription factors or CTCF molecules are enriched. We anticipate that nucleosomes in these regions are either acetylated or depleted.

Because regions characterized by nucleosome depletion and acetylation are correlated and likely to overlap at the current 1-kb resolution, we randomly select 2*m* sites within the modeled acetylated regions of each domain as cohesin-bound sites and connect them with *m* loops under the condition that the loop length is larger than *l*_min_. We generated 10 chains with varying acetylation patterns, defining a set *L* for each chain. Simulations were performed under two cohesin conditions: a loop-constrained condition (control; *L* defined as above, with five loops per 200-kb domain) and a cohesin-depleted condition (Δcohesin; *L* = ∅, i.e., no loop constraints).

We consider two scenarios for generating *L*: one in which loops connect the randomly chosen 2*m* acetylated sites, allowing for mutual crossings of loops, and another in which loops are replaced until all loops are free of crossing. While crossings of loops may not occur under the loop-extrusion mechanism ^149,150^, they can happen through the loop-capture mechanism ^151^ ^152^. In Fig. 8 in the main text, loop crossing is prohibited. In Fig. S10d-e, we compare two scenarios: one allowing loop crossing and the other prohibiting it. This comparison demonstrates that allowing loop crossings slightly enhances the looping effects on chromatin movement.

We define the parameter σ = 30‒40 nm as a unit of spatial length, τ = 1 s as a unit of time, and *k*_B_*T* as a unit of energy. The parameters were set as 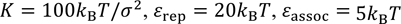, and σ_assoc_ = 1.5σ. The mobility parameter μ was chosen for the simulated MSD of the beads to align with the order of magnitude of the observed MSD, with μ = 200σ^2^/(τ*k*_B_*T*). We should note that *k*_B_*T* is absorbed in the other parameters with this setting, allowing to put *k*_B_*T* = 1 without loss of generality. The rate of association of beads to form attractive pairs was set to approximate the frequency of diffusive movement of nucleosomes as *k*_assoc_ = 200 τ^-1^. We used *k*_dissoc_ = 50 τ^-1^, considering that the dissociation rate is lower in the crowded cellular environment than the value for the dissociation of attractive nucleosome pair observed in vitro^153^. We used the acetylation fraction Φ = 20% and the average region size *w* = 2 kb. It was suggested that the distance between two cohesin molecules bound on chromatin is 30∼70 kb ^154^, and we assumed that the number of cohesin molecules bound on each 200-kb domain is *m* = 5 and set *l*_min_ = 30 kb.

We conducted a simulation for each chain in an open boundary space. The initial conformation of the chain at *t* = 0 was generated as a random walk consisting of 40 consecutive steps. This structure was fine-grained by interpolating 40 positions with a third-order spline, resulting in 405 beads placed equidistantly along the curve. To monitor MSD, we imposed a constraint that the center of mass of the chain does not move, mimicking a scenario where the chain is gently confined within a finite space. The simulation time step was set to δ*t* = 10^-5^ τ, and both MSD and the mixing score were monitored over 5 × 10^6^steps, following a relaxation run of 10^7^steps. One trajectory was obtained with each of 10 chains having different acetylation and loop patterns, and the total 10 trajectories were used to analyze MSD and the mixing score in each scenario of simulations.

Mixing score: to quantify the spatial mixing between the two domains, we computed a mixing score *M* = (*R*_11_ + *R*_22_)/*R*_12_, where *R*_*a b*_ is the average distance between beads in domain *a* and those in domain 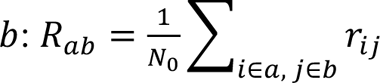. Here, *r*_*i*j_ is the distance between beads *i* and *j, N*_0_ = *n*(*n* − 1) for *a* = *b*, *N*_0_ = *n*^2^ for *a* ≠ *b*, and *n* = 200 is the number of beads in each domain. With this definition, *M* ≈ 2 indicates strong mixing (domains largely merged), whereas *M* ≈ 0.9 indicates strong separation.

### Intron-seqFISH probe selection and design

Candidate intron-seqFISH target genes were selected based on genome-wide transcriptional bursting parameters quantified in HCT116 cells, with an emphasis on genes showing no significant change in expression after acute RAD21 degradation ^155^. To avoid regimes where co-burst differences become difficult to resolve, targets were further restricted to an intermediate bursting/expression range (mean expression 1–3 and burst frequency 0.0033–0.01). Because many genes meeting these criteria were located on chromosome 1, we prioritized chromosome 1 and computed pairwise genomic distances among candidates to maximize the number of analyzable gene pairs within tens of megabases (≤ 40 Mb).

To design intron-targeting primary oligonucleotides, we evaluated probe designability in PaintSHOP and, in practice, extracted intronic genomic intervals for each shortlisted gene and retrieved pre-designed oligonucleotides from a PaintSHOP “full probe file” (DNA-FISH oligo resource) that overlapped these intronic regions ^156^. Genes with fewer than 15 available intronic probes were excluded to ensure robust detection sensitivity, yielding seven target genes (*BTF3L4, CAMTA1, PRPF38B, PTP4A2, RABGGTB, SYF2, and THRAP3*). For genes with many candidate probes, a maximum of 30 probes per gene were selected by prioritizing probes proximal to the transcription start site (TSS) and with favorable off-target/repeat metrics.

Probe selection was implemented as follows. Gene genomic coordinates and strand information were obtained via Ensembl BioMart export, and the TSS was defined as the gene start coordinate for + strand genes and the gene end coordinate for − strand genes. Candidate probes were ranked by the absolute distance between the probe midpoint and the TSS. Probes were first selected within ± 5 kb of the TSS; if fewer than 30 probes were obtained, the window was progressively expanded up to ± 50 kb, and if fewer than 30 probes were available even after expansion, all available probes were used. Candidate probes were filtered to prioritize probes with repeat_seq = 0 (relaxed to repeat_seq ≤ 1 if needed) and off_target = 0 (relaxed to off_target ≤ 1 if needed), with additional ranking based on sequence complexity (e.g., max_kmer) and melting temperature metrics as needed. To reduce local redundancy, a greedy thinning step enforcing a minimum probe–probe spacing of 150 bp was applied, after which 30 probes were retained when possible. To generate RNA-complementary probe sequences, probe sequences were oriented according to gene strand: because PaintSHOP outputs sequences corresponding to the genomic + strand, probes targeting + strand genes were reverse-complemented, whereas probes for − strand genes were used as provided (already matching the reverse-complement orientation relative to the transcribed sequence). Each final primary probe was constructed with a central gene-specific target-complementary sequence flanked by two copies of the secondary-probe binding sequence on both the 5′ and 3′ ends. The complete list of primary probe oligonucleotides, as well as corresponding secondary and readout probes, is provided in Supplementary Table 1. All probe pools were synthesized as oPools (Integrated DNA Technologies, IDT) and used for intron-seqFISH experiments.

### Intron-seqFISH

Cell culture inserts (ibidi, 80209) were fitted on 24 mm × 50 mm No. 1S glass coverslips (thickness 0.16–0.19 mm; Matsunami) that were coated with poly-D-lysine (0.1 mg/mL; Thermo Fisher Scientific, A3890401). HCT116 CMV-OsTIR1(F74G) RAD21-mAID-mClover H2B-Halo cells were seeded at 1 × 10⁴ cells per insert well in McCoy’s 5A medium supplemented with 10% fetal bovine serum (FBS) and cultured overnight. Cells were then labeled with 100 nM JFX549-HaloTag ligand for 1 h at 37 °C, washed 3 times with fresh McCoy’s 5A + 10% FBS, and allowed to recover for 1 h at 37 °C. For cohesin depletion, cells were incubated with 1 µM 5Ph-IAA or vehicle control for 3 h; two biological replicates were prepared for each condition.

Cells were washed with D-PBS, fixed with 4% paraformaldehyde (PFA) for 10 min at room temperature, and washed in D-PBS (3 × 5 min). Samples were then exchanged into 70% ethanol and stored at −20°C prior to intron-seqFISH.

To enable repeated buffer exchange during sequential hybridization–imaging cycles, a thin flow chamber was assembled on the sample coverslip. Before flow-chamber assembly, ethanol was completely removed by aspiration and the samples were allowed to air-dry briefly at room temperature. A Fluorescent Friendly™ double-sided adhesive film (FL-S-2L; Grace Bio-Labs) was cut into a rectangular frame (24 mm × 50 mm) with a central elliptical aperture (major axis ≈ 22 mm, minor axis ≈ 19 mm; projected area ≈ 328 mm²; film thickness 0.28 mm). One adhesive face was laminated onto the sample coverslip such that the specimen region was centered in the aperture. The opposite adhesive face was sealed with a custom cover glass (26 mm × 76 mm × 2 mm; Matsunami) bearing two circular access ports (Ø 1.3 mm, center-to-center spacing 18 mm) serving as inlet and outlet. This formed a leak-tight chamber (∼0.28 mm internal height; ∼92 µL volume) for sequential hybridization, washing, and imaging.

After rehydration in 2× SSC, samples were permeabilized with 0.5% Triton X-100 in PBS for 15 min at room temperature. Blocking was performed by incubating samples in PBS containing 10 mg/mL UltraPure bovine serum albumin (Invitrogen, AM2616), 0.3% Triton X-100, 0.1% dextran sulfate, and 0.5 mg/mL sheared salmon sperm DNA (Invitrogen, AM9680) for 15 min at room temperature. Samples were rinsed with 2× SSC three times.

Primary probe mixtures were prepared at 1 nM per probe in hybridization buffer consisting of 50% formamide, 10% dextran sulfate, and 2× SSC. Hybridization was performed at 37 °C for 72h in a humidified chamber. Following hybridization, samples were washed in 55% formamide wash buffer (55% formamide, 2× SSC, 0.1% Triton X-100) for 30 min at room temperature, followed by three washes in 4× SSC.

Sequential imaging was performed as ten rounds of hybridization–washing–imaging. In each round, the sample was incubated with a target-specific secondary probe together with a universal Alexa Fluor 647-labeled readout oligonucleotide. Secondary and readout probes were diluted together in EC buffer (10% ethylene carbonate, 2× SSC) to final concentrations of 0.1 µM (secondary probe) and 0.4 µM (readout oligo) and hybridized for 20 min at room temperature.

After each 20 min readout hybridization, excess probe was removed by three rinses in 4× SSC, followed by a brief wash in 4× SSC + 0.1% Triton X-100. Samples were then stringently washed in 12.5% wash buffer (12.5% formamide, 2× SSC, 0.1% Triton X-100) and rinsed three additional times in 4× SSC. The buffer was replaced with a DAPI staining solution (4× SSC, 0.1% Triton X-100, 5 µg/mL DAPI) and incubated for 1 min at room temperature. Finally, the chamber was filled with an anti-bleaching imaging buffer consisting of 50 mM Tris-HCl (pH 8.0), 4× SSC, 3 mM Trolox (Sigma-Aldrich, 238813), 10% D-(+)-glucose (Nacalai Tesque, 16806-25), catalase (Sigma-Aldrich, C3155; 1:100 dilution), and glucose oxidase (1 mg/mL) (Sigma-Aldrich, G2133).

Imaging was performed on a Nikon Ti-2 inverted microscope equipped with a Yokogawa CSU-W1 spinning-disk confocal scanner, a Plan Apo λD 60× oil-immersion objective (NA 1.42), and an ORCA-Fusion sCMOS camera (Hamamatsu Photonics), controlled by Nikon NIS-Elements. Excitation was provided by 405 nm, 561 nm, and 637 nm lasers (LightHUB+, Omicron). Z-stacks spanning 20 µm were acquired at 500 nm z-steps (41 optical sections), yielding a voxel size of 110 nm × 110 nm × 500 nm.

In the first cycle, imaging was performed for Alexa 647 (readout), HaloTag (JFX549), and DAPI. In subsequent cycles, imaging was performed for Alexa 647 (readout) and DAPI only. After imaging each round, the fluorophore-labeled readout oligo was stripped by incubation in 55% formamide stripping buffer (55% formamide, 2× SSC, 0.1% Triton X-100) for 15 min at room temperature, followed by rinsing in 4× SSC to remove formamide. The next round’s readout oligo was then introduced, and the cycle was repeated.

Round order was: round 1 *BTF3L4*, round 2 no probe, round 3 *BTF3L4*, round 4 *CAMTA1*, round 5 *PRPF38B*, round 6 *PTP4A2*, round 7 *RABGGTB*, round 8 *SYF2*, round 9 *THRAP3*, round 10 no probe. “No-probe” rounds were included as negative controls to monitor residual signal and stripping efficiency. Round 03 repeated *BTF3L4* to validate round-to-round registration and detection reproducibility (see Fig. S10h)

Raw images from all rounds were organized per field of view and channel. Image files for each round and field of view were imported into Fiji (ImageJ). For quality control and downstream segmentation, maximum-intensity projection (MIP) images were generated. After channel splitting, the readout (Alexa Fluor 647) and DAPI channels were re-merged into a two-channel stack, and all rounds were concatenated and converted into a 4D hyperstack (2 channels × Z × rounds).

Inter-round rigid drift was first corrected in Fiji using Correct 3D drift with the DAPI channel as the reference. After drift correction, stacks were cropped to a user-defined region of interest, and the Z-range containing the specimen was determined automatically from intensity profiles to exclude empty/zero-padded regions. The resulting drift-corrected stacks were exported as per-field-of-view TIFF files and used as input for subsequent deformable registration.

Deformable registration was performed in Python using FireANTs ^157^. For deformation estimation, DAPI volumes were winsorized (clipped at the 0.5th and 99.5th percentiles) and rescaled to the [0,1] range. The DAPI volume from round 1 was used as the fixed reference, and each round was registered as a moving image using greedy optimization (cross-correlation similarity; kernel size 5) at a single resolution scale (factor 4; 200 iterations; Adam optimizer with learning rate 0.5) with Gaussian smoothing regularization of the deformation field (smooth_grad_sigma = 1.0; smooth_warp_sigma = 0.5). The resulting deformation field was then applied to both the unnormalized DAPI and readout (Alexa Fluor 647) volumes for each round.

Nuclei were segmented from DAPI maximum-intensity projections using Cellpose with the pretrained “nuclei” model. Resulting masks were inspected and manually corrected where needed before downstream quantification.

Nascent intronic RNA signals were detected from the registered Alexa 647 (FISH) volumes using bigFISH spot detection ^158^. Spot detection was performed in 3D using the acquisition voxel size (110 × 110 × 500 nm) and an empirically selected spot radius in nanometers. Detected spots were optionally refined by subpixel localization. Spots were then assigned to nuclei using the nuclear masks. To reduce false positives and over-counting, intensity filtering was applied using empirically determined intensity thresholds, and for each gene in each cell, up to two brightest spots were retained to represent (at most) two alleles in a near-diploid cell line.

For each gene *A* and experimental group, the burst frequency was computed as:

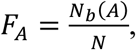

where *N* is the total number of analyzed cells in the group and *N_b_*(*A*) is the total number of detected nascent transcription spots (bursting sites) for gene *A* summed across all cells in that group.

For each gene pair (A, B), co-bursting was defined: a cell was classified as co-bursting if it contained at least one spot for gene A and at least one spot for gene B, and the minimum 3D distance between any A–B spot pair in that cell was ≤ 1 µm. The co-burst frequency was: co-burst frequency = *N_co_*/*N*, where *N_co_* is the number of co-bursting cells.

A normalized co-burst frequency was computed as:

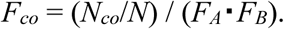

For comparisons between conditions (e.g., RAD21 depleted vs control), Δ*F_co_* was computed for each gene pair as the difference in *F_co_* between groups.

## Acknowledgments

We are grateful to Dr. K. M. Marshall for critical reading of this manuscript and Mr. A. Tsurumune (NIKON) for setting up 3D-SIM. We thank Dr. H. Kimura for providing antibodies, Dr. T. Tanaka for providing the HT1080 cell line, and Dr. S. Ide for technical assistance with pull-down and library preparation. We thank Dr. L. Lavis for providing PA-JF549-HaloTag, PA-JF646-HaloTag, and JFX549-HaloTag ligands. We also thank Dr. B. Zhang and Dr. J.M. Paggi for sharing unpublished results and helpful discussion, Mr. A. Otsuka and Ms. H. Ochi for establishing HeLa and HCT116 cells expressing H3.3-Halo, Dr. S. Shinkai for assistance with angle-distribution and two-point MSD analyses, and Dr. K. Hibino for establishing HCT116 cells expressing H2B-Halo and for helpful discussions. Finally, we thank Dr. T. Cremer, Dr. C. Cremer, Dr. M. Cremer, Dr. Y. Shimamoto, Dr. H. Niki, and all members of the Maeshima laboratory for valuable discussions and continuous support.

## Author contributions

S.I., M.A.S., and K. Maeshima designed the research. S.I., M.A.S, and K. Minami performed the experiments including cell generation, imaging, and analyses. S.I. led the initial development of the study, including experimental design, generation and analysis of the original data, and drafting of the initial manuscript that established the core narrative of the bioRxiv submission (ver. 1). M.A.S. established the 2-point MSD analysis, RL analysis, 3D-SIM, and STORM. S.T. and K.N. performed some biochemical experiments. S.S.A. and M.S. developed the RL analysis. S.T., K. Maeshima, and K. Minami purified H3.3-Halo-labeled nucleosomal DNA and prepared the sequencing library. L.S. assisted with 3D-SIM setup and analysis. A.T. performed sequencing. K.H. and K.K. analyzed the sequencing data. M.T.K. contributed to AID2 cell generation. A.K. and Y.N. contributed to some genomics analysis. T.N. generated the head-tethered cohesin cells. Y.K., Y.O. and H.O. performed co-burst analysis. S.F. and M.S. performed simulation. L. Xiong and L. Xie performed ATAC-seq. M.A.S., S.I., K. Minami and K. Maeshima wrote the manuscript with input from all authors.

## Data and materials availability

The pull-down sequencing data have been deposited in the DDBJ BioProject database under accession numbers PRJDB17378 (HeLa H2B-HaloTag pull-down) ^108^, PRJDB17380 (HeLa H3.3-HaloTag pull-down), and PRJDB39674 (HCT116 H2B-HaloTag and H3.3-HaloTag pull-downs). ATAC-seq data have been deposited in GEO (GSE318385). The scripts for RL-algorithm classification, TrackMate Batch Tracker, SIM image analysis, and two-point MSD calculation are available at https://doi.org/10.5281/zenodo.16959115. The codebase for the polymer model of chromatin domains is available at https://doi.org/10.5281/zenodo.18404051. Other data are available in the main text and the supplementary materials.

## Supplementary Materials

Figs. S1 to S15

Movies S1 to S9

Table S1

