## Supplementary Materials for "Cohesin prevents local mixing of condensed euchromatic domains in living human cells"

**This PDF file includes:**

Figs. S1 to S15
Legends for Movies S1 to S9

**Other Supplementary Materials for this manuscript include the following:**

Movies S1 to S9

**References for Supplementary Materials**

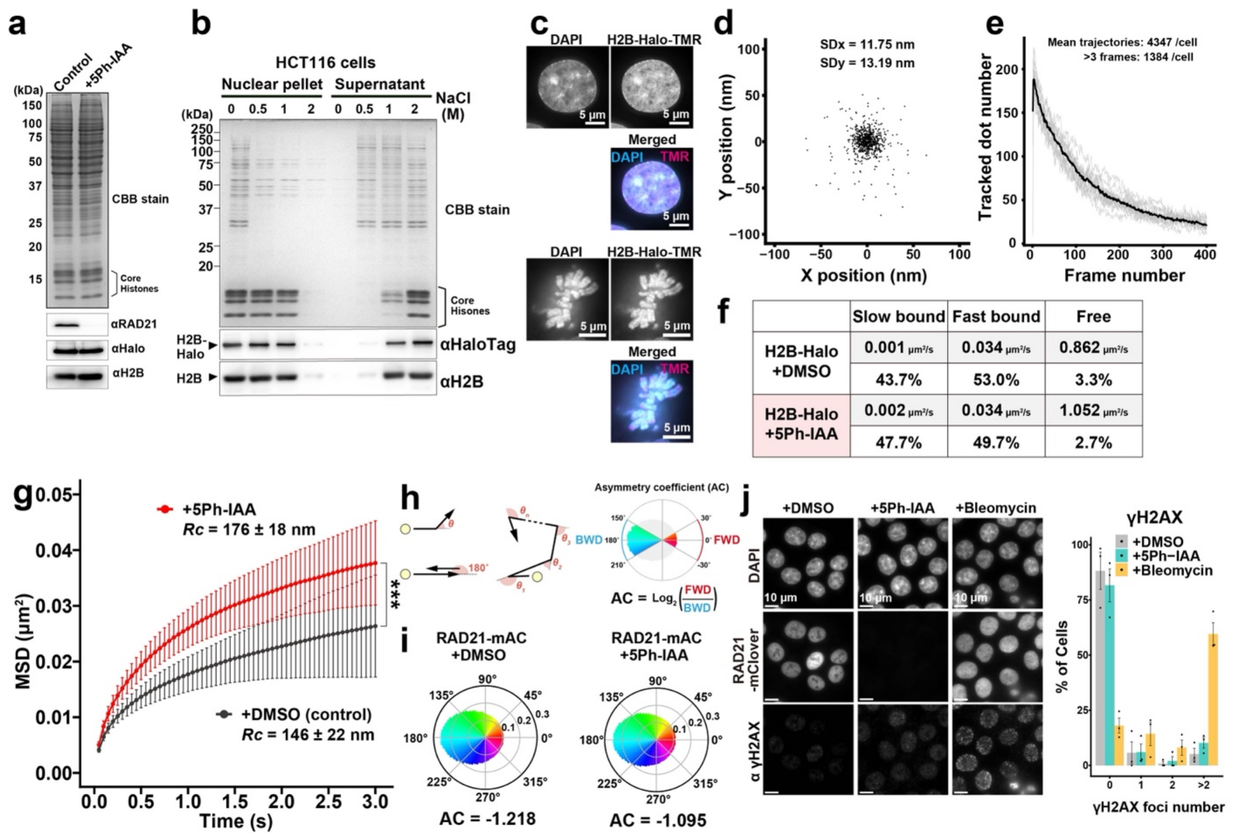


**Fig. S1: Single-nucleosome imaging using HCT116 cells with RAD21-AID2.**
**a**, Immunoblotting of HCT116 cells expressing H2B-HaloTag (H2B-Halo) and RAD21-mAID-mClover (mAC) (left lane). Verification of RAD21 depletion using AID2 by immunoblotting (right lane). **b**, Stepwise salt washing of nuclei expressing H2B-Halo: nucleosomes with H2B-Halo have biochemically similar stability as those with endogenous H2B. The nuclei isolated from HCT116 cells expressing H2B-Halo were washed with increasing concentrations of NaCl. The resultant nuclear pellets (left) and supernatants (right) were analyzed by SDS-PAGE, and subsequently stained with Coomassie brilliant blue (top) or immunoblotted for H2B and HaloTag (bottom). Note that endogenous H2B and H2B-Halo started to dissociate from chromatin with 1 M NaCl and were detected in the supernatant fraction, suggesting that nucleosomes with H2B-Halo have similar stability as those with endogenous H2B. **c**, HCT116 cells expressing H2B-Halo were fluorescently labeled with DAPI and an excess amount of TMR-HaloTag ligand. Merged images are shown with DNA in cyan and H2B-Halo in magenta. Top: interphase cell; bottom: mitotic cell. **d**, Position determination for the accuracy of H2B-Halo-TMR. Distribution of nucleosome displacements from their centroid in the x–y plane in the 50-ms interval. $N = 10$ nucleosomes in an FA-fixed cell. SDx and SDy were 11.8 nm and 13.2 nm, respectively. **e**, Example of the number of detected dots per frame ($N = 15$ cells). The dot count decreases over time due to photobleaching, while new dots occasionally appear as they come into focus. Note that the average number of trajectories does not correspond to individual nucleosomes, as some trajectories may be fragmented. **f**, Diffusion coefficients and percentages of fractions estimated by Spot-On. H2B-Halo in HCT116 RAD21-mAC labeled with TMR is examined by 10

ms/frame imaging. Note that freely diffusing fractions are almost negligible in all cases. **g**, MSD plots (± SD among cells) of H2B-Halo in HCT116 RAD21-mAC cells under the indicated conditions: DMSO (black, $N = 30$ cells), 5Ph-IAA (ΔRAD21, red, $N = 30$ cells). ***, $P = 4.32 \times 10^{-8}$ (DMSO vs 5Ph-IAA) by two-sided Kolmogorov-Smirnov test. Radius of constraints is DMSO: 146 ± 22 nm; 5Ph-IAA: 176 ± 18 nm. **h**, Schematic for angle-distribution analysis and asymmetric coefficient (AC). AC indicates deviation from a homogeneous distribution and is negative for angular distributions where the pulling-back force is dominant. **i**, Angle distributions of DMSO (816,941 angles) and 5Ph-IAA (ΔRAD21; 828,733 angles). **j**, Verification of RAD21 depletion by the mClover signal and of DNA damage by γH2AX immunostaining. Center: RAD21-mClover intensity quantification of fixed cells. Mean values: $2.3 \times 10^3$ for DMSO ($N = 59$ cells), 43 for siCTCF ($N = 51$ cells), and $2.2 \times 10^3$ for Bleomycin ($N = 53$ cells). ***, $P < 0.0001$ by Wilcoxon rank sum test for DMSO vs 5Ph-IAA ($P = 1.7 \times 10^{-19}$) and 5Ph-IAA vs Bleomycin ($P = 1.4 \times 10^{-18}$). N.S., not significant for DMSO vs Bleomycin ($P = 0.11$). Right: fractions of cells with the noted numbers of γH2AX foci. The mean value of three independent experiments is plotted as a bar graph.

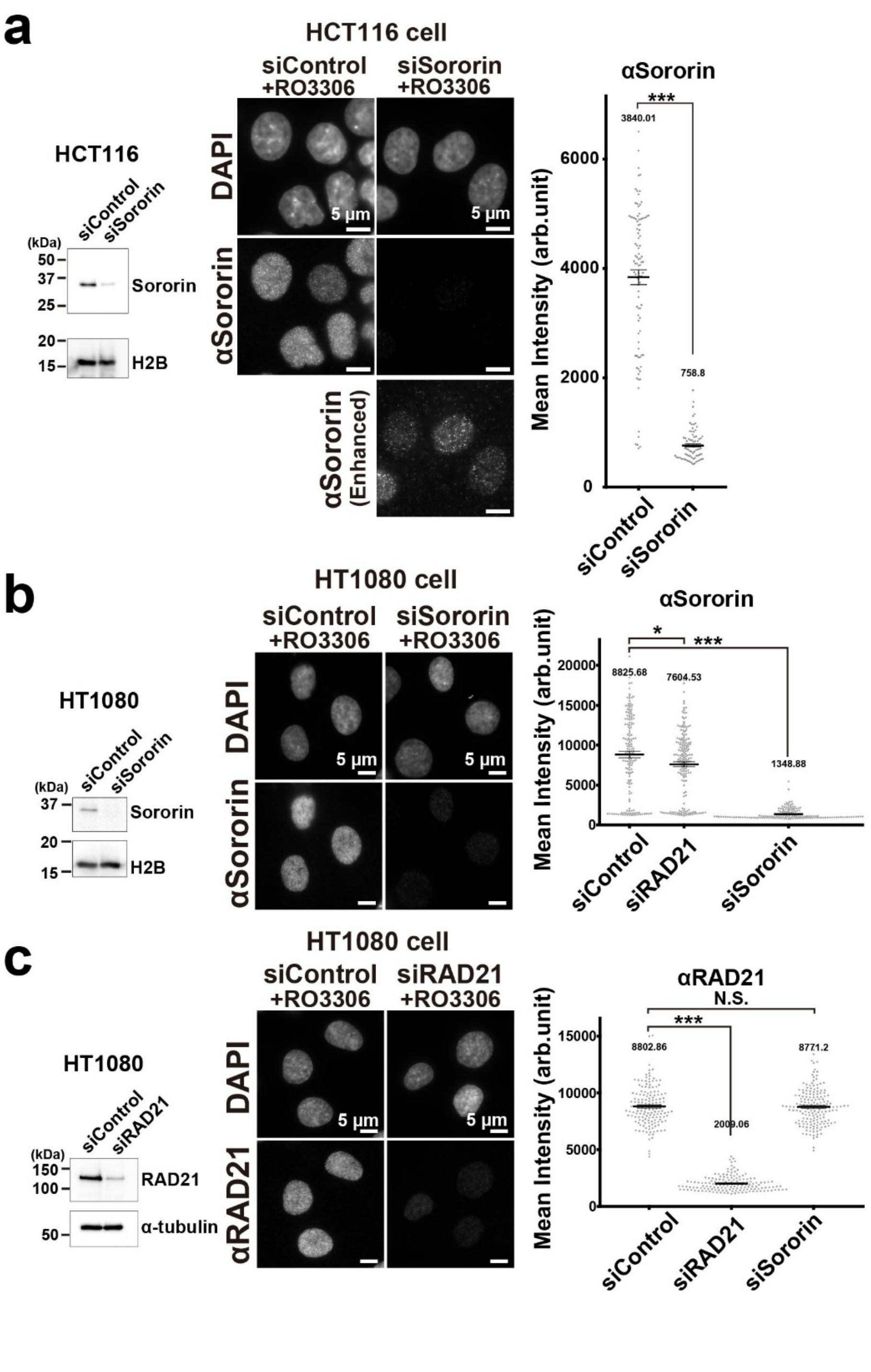


**Fig. S2: Validation of knockdown efficiency in HCT116 and HT1080 cells.**
**a**, Verification of the Sororin knockdown by immunoblotting (left: 77.3% reduction) and immunostaining (mid, right). Mean intensity values in immunostaining: $3.8 \times 10^3$ for siControl ($N = 97$ cells) and $7.6 \times 10^2$ for siSororin ($N = 82$ cells). ***, $P < 0.0001$ ($P = 3.2 \times 10^{-28}$) by Wilcoxon rank sum test. **b**, Verification of the Sororin knockdown by immunoblotting (left: 87.3% reduction) and immunostaining (mid, right). Mean intensity values in immunostaining: $8.8 \times 10^3$ for siControl ($N = 160$ cells), $7.6 \times 10^3$ for siRAD21 ($N = 183$ cells), and $1.3 \times 10^3$ for

siSororin ($N$ = 157 cells). ***, $P < 0.0001$ by Wilcoxon rank sum test for siControl vs siSororin ($P = 5.0 \times 10^{-41}$). *, $P < 0.05$ for siControl vs siRAD21 ($P = 0.030$). **c**, Verification of the RAD21 knockdown by immunoblotting (left: 82.0% reduction) and immunostaining (mid, right). Mean intensity values in immunostaining: $8.8 \times 10^3$ for siControl ($N$ = 154 cells), $2.0 \times 10^3$ for siRAD21 ($N$ = 153 cells), and $8.8 \times 10^3$ for siSororin ($N$ = 171 cells). ***, $P < 0.0001$ by Wilcoxon rank sum test for siControl vs siRAD21 ($P = 7.9 \times 10^{-52}$). N.S., not significant for siControl vs siSororin ($P = 0.90$).

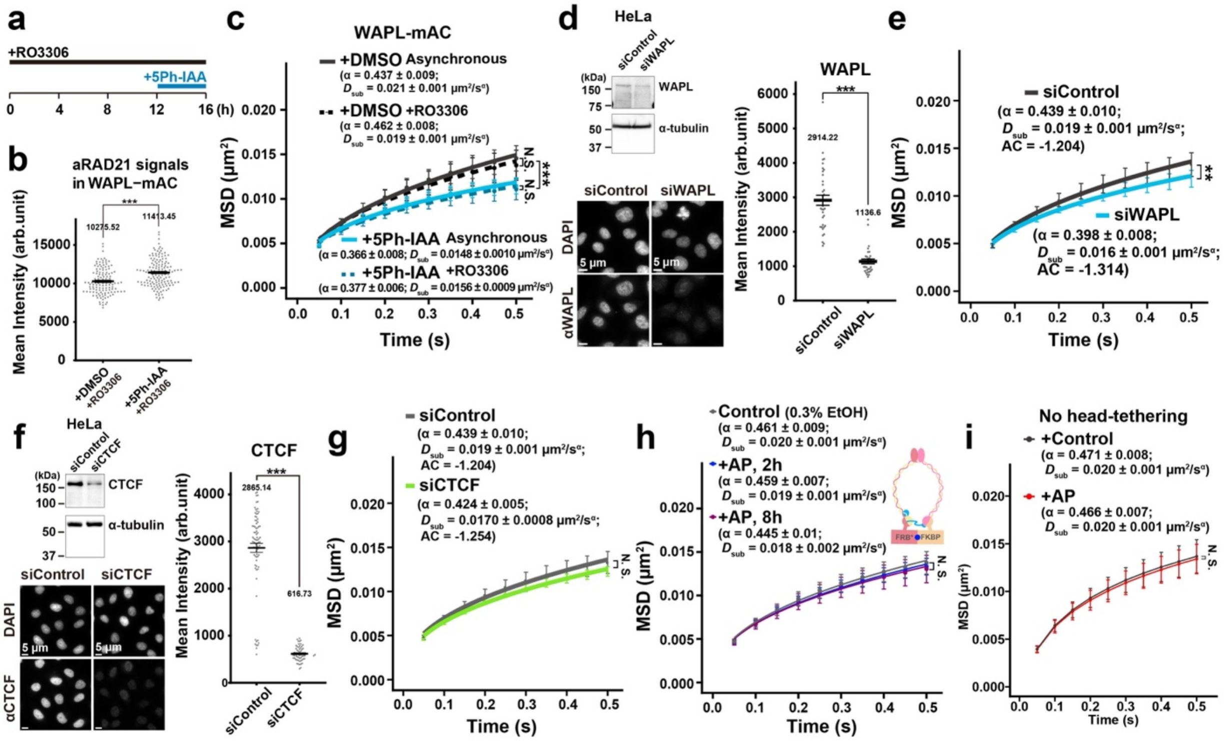


**Fig. S3: WAPL and CTCF depletion using siRNA in HeLa cells.**

**a**, Schematic for the treatment timeline of cells with RO3306 and 5Ph-IAA. **b**, Examination of RAD21 intensity in WAPL-depleted cells by immunostaining. Mean intensity values: $1.0 \times 10^4$ for DMSO ($N$= 125 cells) and $1.1 \times 10^4$ for 5Ph-IAA ($N$ = 146 cells). ***, $P < 0.0001$ ($P = 6.1 \times 10^{-7}$) by Wilcoxon rank sum test. **c**, MSD plots (± SD among cells) of H2B-Halo in HCT116 cells with the indicated conditions: DMSO asynchronous (black solid line, $N$ = 29 cells); DMSO+RO3306 (black dotted line, $N$ = 30 cells); 5Ph-IAA asynchronous (light blue solid line, $N$ = 30 cells); 5Ph-IAA+RO3306 (light blue dotted line, $N$ = 28 cells). ***, $P < 0.001$ for DMSO asynchronous vs 5Ph-IAA asynchronous ($P = 1.1 \times 10^{-6}$) and DMSO+RO3306 vs 5Ph-IAA+RO3306 ($P = 1.1 \times 10^{-11}$). N.S., not significant for DMSO asynchronous vs DMSO+RO3306 ($P = 0.11$) and 5Ph-IAA asynchronous vs 5Ph-IAA+RO3306 ($P = 0.34$) by the two-sided Kolmogorov–Smirnov test. **d**, Verification of WAPL depletion by immunoblotting (left top: 55.8% reduction) and immunostaining. Mean intensity values in immunostaining: $2.9 \times 10^3$ for siControl ($N$ = 37 cells) and $1.1 \times 10^3$ for siWAPL ($N$ = 31 cells). ***, $P < 0.0001$ ($P = 4.1 \times 10^{-16}$) by Wilcoxon rank sum test. **e**, MSD plots (± SD among cells) of H2B-Halo in HeLa cells with the indicated conditions: siControl (black, $N$ = 15 cells) and siWAPL (blue, $N$ = 15 cells). **, $P < 0.01$ by the two-sided Kolmogorov–Smirnov test for siControl vs siWAPL ($P = 7.7 \times 10^{-3}$). **f**, Verification of CTCF depletion by immunoblotting (left top: 80.1% reduction) and immunostaining. Mean intensity values in immunostaining: $2.9 \times 10^3$ for siControl ($N$ = 80 cells) and $6.1 \times 10^2$ for siCTCF ($N$ = 60 cells). ***, $P < 0.0001$ ($P = 1.6 \times 10^{-22}$) by Wilcoxon rank sum test. **g**, MSD plots (± SD among cells) of H2B-Halo in HeLa cells with the indicated conditions: siControl (black, $N$ = 15 cells) and siCTCF (light green, $N$ = 15 cells). N.S., not significant ($P = 0.18$) by the two-sided Kolmogorov–Smirnov test. **h**, MSD plots (± SD among cells) of H2B-Halo in HCT116 cells with the indicated conditions: Control (black, $N$ = 20 cells); AP21967 (AP) 2 h (blue, $N$ = 20 cells); AP 8 h (purple, $N$ = 20 cells). N.S., not significant for

Control vs AP 2 h ($P$ = 0.17) and Control vs AP 8 h ($P$ = 0.081) by the two-sided Kolmogorov–Smirnov test. **i**, MSD plots (± SD among cells) of H2B-Halo in HCT116 cells with no head-tethering components. Control (black, $N$ = 30 cells); AP21967 (AP) 2 h (red, $N$ = 30 cells). N.S., not significant ($P$ = 0.59) by the two-sided Kolmogorov–Smirnov test, indicating that AP21967 does not affect nucleosome motion.

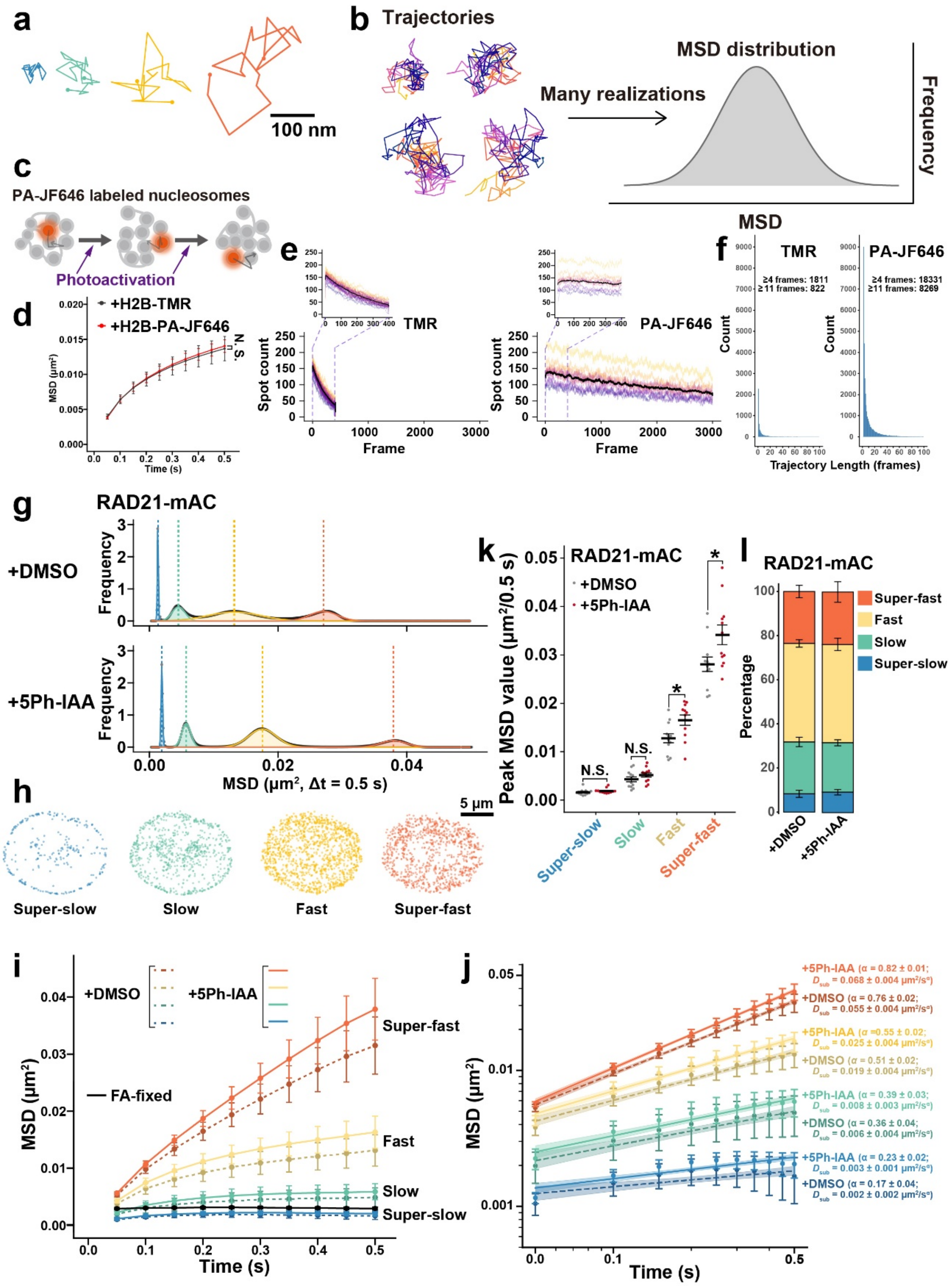


**Fig. S4: Comparison of dot number and trajectory length between TMR and PA-JF646.** **a**, Representative trajectories of four tracked single nucleosomes, each belonging to a different subpopulation. **b**, Schematic of the RL algorithm. **c**, Schematic for PA-JF646 labeling and photoactivation of labeled nucleosomes. **d**, MSD plots (± SD among cells) of H2B-Halo in HCT116 cells, labeled with TMR (black, $N$ = 30 cells); PA-JF646 (red, $N$ = 30 cells). N.S., not significant ($P$ = 0.09) by the two-sided Kolmogorov–Smirnov test, indicating that a similar nucleosome motion was observed with two fluorescent dyes. **e**, Number of detected dots per frame in H2B-Halo–TMR (left, $N$ = 30 cells) and H2B-Halo–PA-JF646 (right, $N$ = 11 cells). Both conditions showed a similar initial dot count. The number of detected dots decreased over time due to photobleaching in both cases; however, in the PA-JF646 condition, newly appearing dots from photoactivation led to a milder decrease. **f**, Distribution of trajectory lengths for TMR-labeled and PA-JF646-labeled H2B-Halo in a representative nucleus. Approximately 10-fold more trajectories were obtained with PA-JF646 labeling. **g**, Distributions of MSD ($\Delta$t = 0.5 s) from H2B-Halo trajectories were denoised using the RL algorithm and fitted with four Gaussian components: "Super-slow" (blue), "Slow" (green), "Fast" (yellow), and "Super-fast" (orange). Black lines indicate the denoised raw data, while colored lines represent the fitted Gaussians. The peak of each Gaussian is marked with a dotted line. Notably, only the "Fast" (yellow) and "Super-fast" (orange) peaks shifted rightward upon cohesin depletion (+5Ph-IAA), indicating further acceleration of these subpopulations. **h**, Nuclear localization of nucleosomes categorized into each subpopulation. Each dot represents the centroid of a trajectory. "Super-slow" and "Slow" trajectories are biased toward the nuclear periphery, presumably corresponding to heterochromatin. **i**, MSD plots (± SD among cells) of subpopulations classified using the RL algorithm based on the 0.5-second MSD distribution. **j**, Log–log plot of MSD corresponding to Fig. S4i. The plots were fitted linearly. Shaded regions represent the 95% confidence interval of the fitting. Note that reliable fitting of $\alpha$ values for the "Super slow" and "Slow" categories is difficult due to their limited mobility. **k**, Peak MSD values of the four Gaussian components. Each dot represents data from a single nucleus. $N$ = 12 cells per condition. *, $P < 0.05$ for "Super-fast" ($P$ = 0.025) and for "Fast" ($P$ = 0.015). N.S., not significant for "Slow" ($P$ = 0.19) and "Super-slow" ($P$ = 0.24) by the two-sided Student's t-test. **l**, Fractions of each subpopulation. Cohesin depletion did not alter the relative proportions of the subpopulations, whereas the fast and super-fast components shifted.

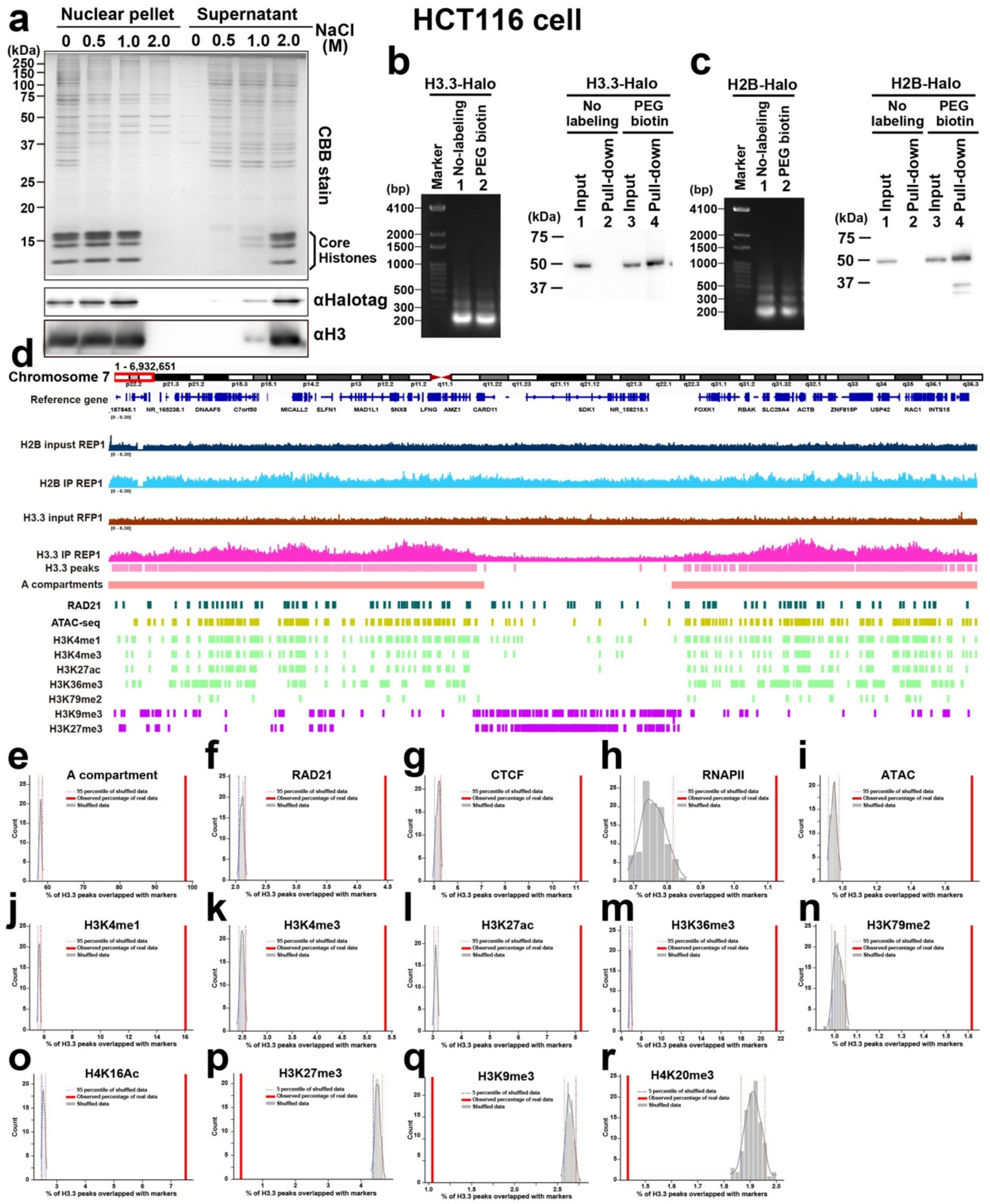


**s**

| | Slow bound | Fast bound | Free |
|---|---|---|---|
| H3.3-Halo +DMSO | 0.004 µm²/s | 0.032 µm²/s | 0.872 µm²/s |
| | 52.0% | 45.9% | 2.1% |
| H3.3-Halo +5Ph-IAA | 0.005 µm²/s | 0.038 µm²/s | 0.909 µm²/s |
| | 38.6% | 59.2% | 2.1% |

**Fig. S5: H3.3-Halo pull-down shows H3.3 is enriched in euchromatin.**
**a**, Stepwise salt washing of nuclei expressing H3.3-Halo. The nuclei isolated from HCT116 cells expressing H3.3-Halo were washed with increasing concentrations of NaCl. The resulting nuclear pellets (left) and supernatants (right) were analyzed by SDS–PAGE, followed by Coomassie brilliant blue staining (top) or immunoblotting for H3 and HaloTag (bottom).

Endogenous H3 and H3.3-Halo started to dissociate from chromatin with 1 M NaCl and were detected in the supernatant fraction, suggesting that nucleosomes containing H3.3-Halo have similar stability to those containing endogenous H3. **b** (left), Agarose gel electrophoresis of nucleosomal DNA purified from HCT116 cells expressing H3.3-Halo. Note that the major fractions are mononucleosomal DNA. (right) Enrichment of the pull-down nucleosomes validated by western blotting with an anti-HaloTag antibody. **c** (left), Agarose gel electrophoresis of nucleosomal DNA purified from HCT116 cells expressing H2B-Halo. Note that the major fractions are mononucleosomal DNA. (right) Enrichment of the pull-down nucleosomes validated by western blotting with an anti-HaloTag antibody. **d,** Another example of H3.3-Halo–nucleosome regions: chr7_1–6,932,651. **e-r**, Distributions of the percentages of (e) A compartment, (f) RAD21, (g) CTCF, (h) RNAP II, (i) ATAC-seq,  active histone marks (j) H3K4me1, (k) H3K4me3, (l) H3K27ac, (m) H3K36me3, (n) H3K79me2, (o) H4K16Ac, and the inactive histone mark (p) H3K27me3, (q) H3K9me3, and (r) H4K20me3 overlapped with 100 randomly shuffled datasets of H3.3-Halo peaks. The red solid line indicates the percentage observed in the real H3.3-Halo dataset, while the dotted red and blue lines indicate the 95th and 5th percentiles of the shuffled distribution, respectively. **s**, Diffusion coefficients and percentages of fractions estimated by Spot-On. H2B-Halo in HCT116 RAD21-mAC labeled with TMR is examined by 10 ms/frame imaging. Note that free diffusing fractions are almost negligible in all cases.

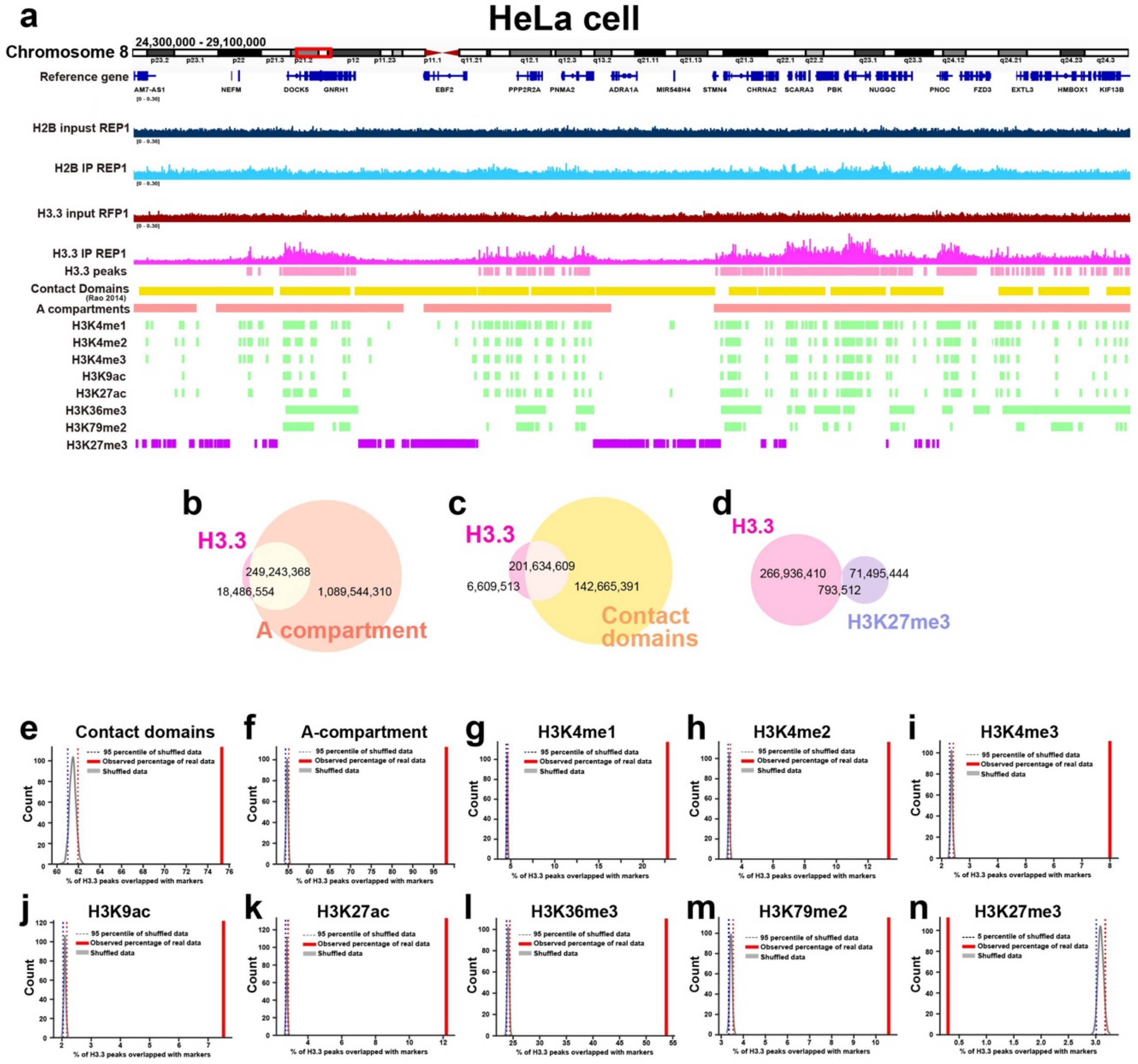


**Fig. S6: H3.3-Halo is also enriched in euchromatin in HeLa cells.**

**a,** Examples of H3.3-Halo–nucleosome regions: chr8_24,300,000–29,100,000. **b-d**, Venn diagrams showing the overlap (in bp) between H3.3-Halo–nucleosome regions and A compartment (b), between H3.3-Halo–nucleosome regions and contact domains (c), and between H3.3-Halo–nucleosome regions and H3K27me3 regions (d). **e-n**, Distributions of the percentages of (**e**) contact domains, (**f**) A compartment, active histone marks (**g**) H3K4me1, (**h**) H3K4me2, (**i**) H3K4me3, (**j**) H3K9ac, (**k**) H3K27ac, (**l**) H3K36me3, (**m**) H3K79me2, and the inactive histone mark (**n**) H3K27me3 overlapped with 1000 randomly shuffled datasets of H3.3-Halo peaks. The red solid line indicates the percentage observed in the real H3.3-Halo dataset, while the dotted red and blue lines indicate the 95th and 5th percentiles of the shuffled distribution, respectively.

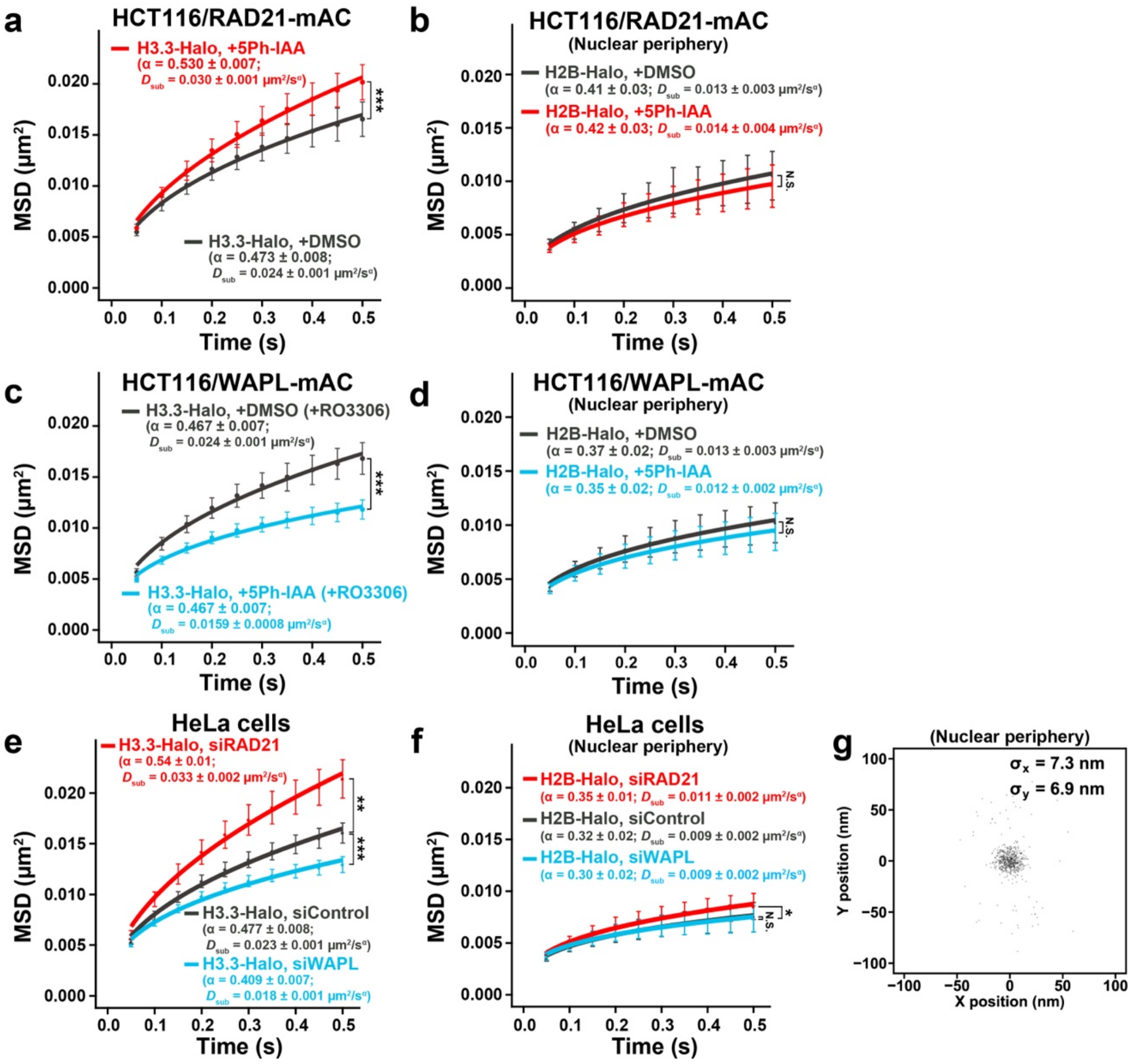


**Fig. S7: Changes in the motion of euchromatic nucleosomes and peripheral nucleosomes upon cohesin depletion.**

**a**, MSD plots (± SD among cells) of H3.3-Halo in HCT116 cells with indicated conditions: DMSO (black, $N = 30$ cells) or 5Ph-IAA (ΔRAD21, red, $N = 30$ cells). ***, $P < 0.001$ by the two-sided Kolmogorov–Smirnov test ($P = 8.2 \times 10^{-12}$). **b**, MSD plots (± SD among cells) of H2B-Halo at the nuclear periphery in HCT116 cells with indicated conditions: DMSO (black, $N = 8$ cells) or 5Ph-IAA (ΔRAD21, red, $N = 9$ cells). N.S., not significant by the two-sided Kolmogorov–Smirnov test ($P = 0.41$). **c**, MSD plots (± SD among cells) of H3.3-Halo in HCT116 cells with indicated conditions: DMSO (black, $N = 30$ cells) or 5Ph-IAA (ΔWAPL, light blue, $N = 30$ cells). ***, $P < 0.001$ by the two-sided Kolmogorov–Smirnov test ($P = 1.7 \times 10^{-17}$). **d**, MSD plots (± SD among cells) of H2B-Halo at the nuclear periphery in HCT116 cells with indicated conditions: DMSO (black, $N = 13$ cells) or 5Ph-IAA (ΔWAPL, light blue, $N = 14$ cells). N.S., not significant by the two-sided Kolmogorov–Smirnov test ($P = 0.26$). **e**, MSD plots

(± SD among cells) of H3.3-Halo in HeLa cells with indicated conditions: siControl (black, $N$ = 14 cells), siRAD21 (red, $N$ = 14 cells), and siWAPL (light blue, $N$ = 15 cells). $P = 1.0 \times 10^{-8}$ by Kruskal–Wallis test, indicating significant overall differences among groups. **, $P < 0.01$ for siControl vs siRAD21 ($P = 1.4 \times 10^{-3}$) and siControl vs siWAPL ($P = 2.5 \times 10^{-3}$). **f**, MSD plots (± SD among cells) of H2B-Halo at the nuclear periphery (basal nuclear surface) in HeLa cells with indicated conditions: siControl (black, $N$ = 15 cells), siRAD21 (red, $N$ = 15 cells), and siWAPL (light blue, $N$ = 15 cells). $P = 4.9 \times 10^{-3}$ by Kruskal–Wallis test, indicating significant overall differences among groups. **, $P < 0.01$ for siControl vs siRAD21 ($P = 7.0 \times 10^{-3}$). N.S., not significant for siControl vs siWAPL ($P = 0.40$). **g**, Position determination for the accuracy of H2B-Halo-TMR on the nuclear periphery. Distribution of nucleosome displacements from their centroid in the x–y plane in the 50-ms interval. $N$ = 11 nucleosomes in an FA-fixed cell. SDx and SDy were 7.3 nm and 6.9 nm, respectively.

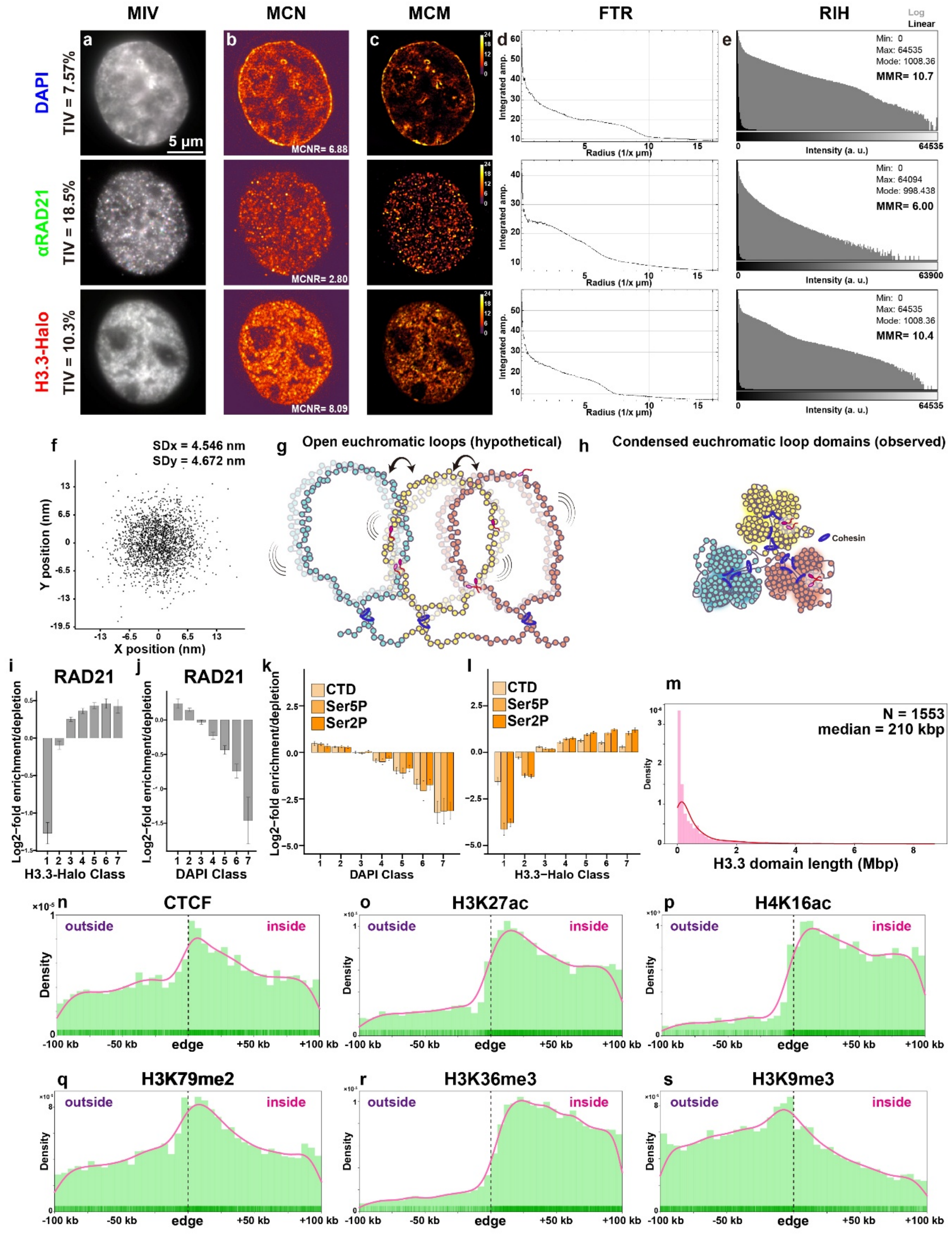


**Fig. S8. Quality control of 3D-SIM and STORM imaging, and localization of histone marks relative to H3.3-Halo domains.**

**a–e**, Quality of 3D-SIM images and reconstructions assessed with SIMcheck [1]. The images also achieved the best N-SIM score (8 on a 1–8 scale). Results for the three channels (DAPI, anti-

RAD21, H3.3-Halo) are shown. **a**, TIV: total intensity variation; MIV: motion & illumination variation. TIV and MIV indicate minimal motion and uniform illumination in the raw images. **b**, MCN: modulation contrast to noise; MCNR: modulation contrast-to-noise ratio. MCN maps the local stripe modulation contrast in the raw data. MCNR represents the average value of features. **c**, MCM: modulation contrast map, indicates the high underlying modulation contrast of the features in the reconstructed image. **d**, FTR: radially averaged Fourier transform. FTR amplitude profile plots indicate an effective lateral resolution of ~100 nm for the blue channel and ~140 nm for the green and red channels. **e**, RIH: intensity histogram of the reconstructions; MMR: max-to-min intensity ratio. Both indicate sufficiently high reconstruction contrast. **f**, Localization precision for STORM (H2B-Halo-HMSiR). Distribution of nucleosome displacements from their centroid in the x–y plane at 10-ms intervals in an FA-fixed cell (*N* = 360 nucleosomes). SDx = 4.546 nm; SDy = 4.672 nm. **g,h**, Hypothetical open euchromatic loops (g) may provide weaker transcriptional insulation between domains. By contrast, the condensed euchromatic loop domains observed here (h) can provide stronger insulation between domains. **i,j**, Bar plots of the $\log_2$-fold enrichment or depletion of RAD21 in each segmented chromatin class relative to class volume for H3.3-Halo-TMR classes (i) and DAPI classes (j). *N* = 37 cells. **k,l**, Bar plots of the $\log_2$-fold enrichment or depletion of RNAP II forms in each segmented chromatin class, relative to class volume. Panels k and l are based on DAPI classes and H3.3-Halo classes, respectively. CTD, *N* = 15 cells; Ser5P, *N* = 19 cells; Ser2P, *N* = 23 cells. **m**, Distribution of detected H3.3 domain lengths. The red line shows a kernel density estimate (KDE) overlaid on the histogram. **n–s**, Distributions of (n) CTCF [2], (o) H3K27me3 [3], (p) H4K16ac [3], (q) H3K79me2 [3], (r) H3K36me3 [3], and (s) H3K9me3 [3] relative to the closest edge of an H3.3 domain. Positive and negative values indicate positions inside and outside H3.3 domains, respectively. The pink line shows a KDE overlaid on the histogram.

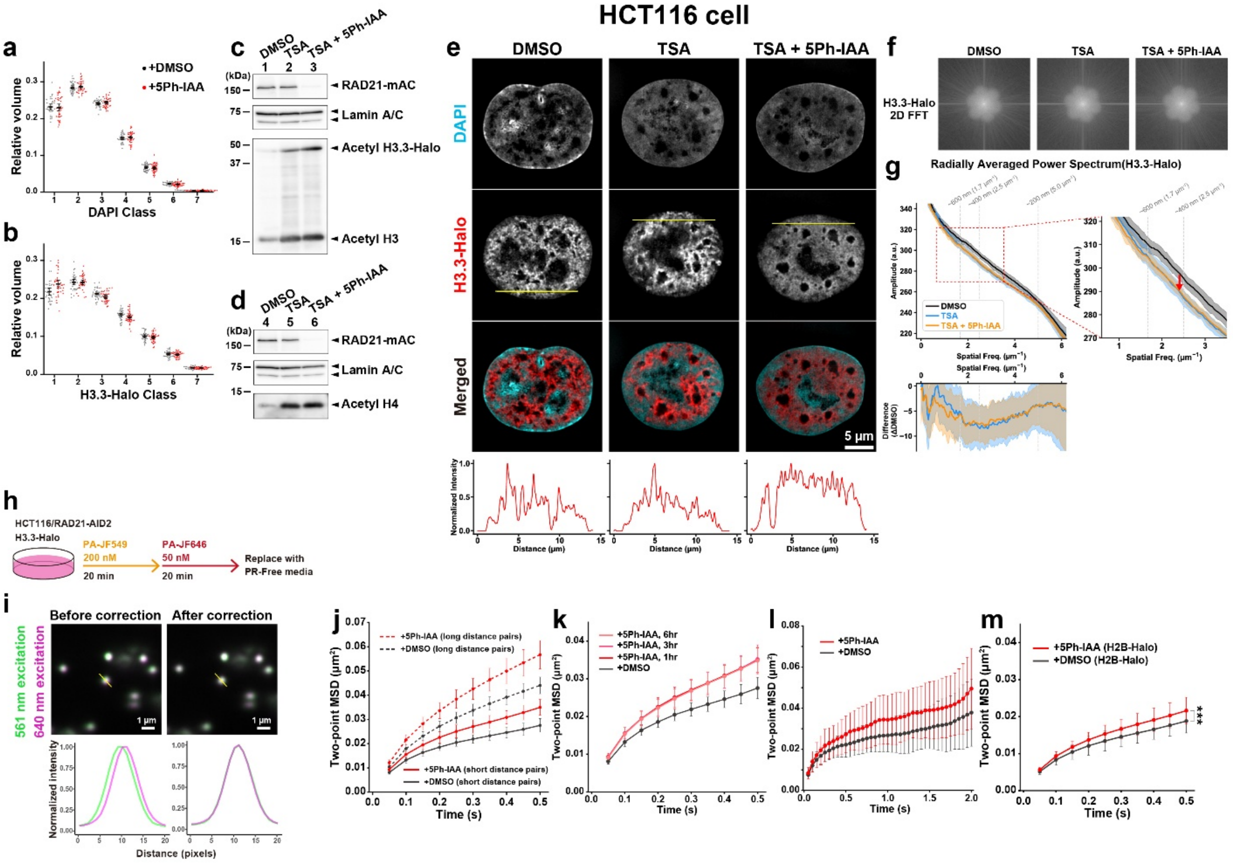


**Fig. S9: "Melting" of chromatin domains and dual-color single molecule imaging.**
**a**,**b**, Proportion of nuclear volume for each class of DAPI (a) and H3.3-Halo (b), showing no significant change after cohesin depletion (mean ± 95% CI). Each dot represents data from a single nucleus. DMSO: $N = 35$ cells. 5Ph-IAA: $N = 39$ cells. **c**,**d**, Immunoblotting of HCT116 cells expressing RAD21-mAID-mClover (mAC) with DMSO treatment (lanes 1,4), TSA (Trichostatin A) treatment (lanes 2,5), or TSA + 5Ph-IAA treatment (cohesin depletion) (lanes 3, 6). Verification of RAD21 depletion using AID2 (top lanes) and histone acetylation with TSA treatment (bottom lanes). **e**, Representative single z-slices of 3D-SIM images on chromatin labeled with DAPI or H3.3-Halo-TMR with DMSO treatment, TSA treatment, or TSA + 5Ph-IAA (cohesin depletion) treatment. Bottom, intensity line profiles of H3.3-Halo at the yellow line of the image. A combination of histone hyperacetylation and cohesin depletion causes "melting" of chromatin domains. **f**, Representative 2D FFT images of the H3.3-Halo-TMR images shown in e. **g**, Radially averaged power spectrum of H3.3-Halo images (shaded regions represent SD among cells, $N = 13$, 6, 10 cells for DMSO, TSA, TSA + 5Ph-IAA treatment, respectively). Decrease in the 300-600 nm range with TSA treatment (histone hyperacetylation) indicates "melting" of chromatin domains. **h**, Schematic of the dual-color labeling strategy. PR, phenol red. **i**, Chromatic aberration was corrected using inter-channel shifts determined from fluorescent beads. Transformation parameters were obtained with the ImageJ plugin *"Descriptor-based registration (2D/3D)"* and applied as an affine transformation to the trajectory data. Representative images before and after correction using this plugin are shown. **j**, Two-point MSD plots (± SD among cells) of H3.3-Halo in HCT116 cells with indicated conditions: DMSO (black, $N = 20$ cells), 5Ph-IAA (ΔRAD21, red, $N = 20$ cells), short distance pairs ($d < 150$ nm)

and long distance ("unrelated") pairs (300 nm < *d* < 1000 nm). Short distances pick up neighboring nucleosomes pairs, while long distances pick up far away (unrelated) nucleosome pairs. **k**, Two-point MSD plots (*d* < 150 nm; ± SD among cells) of H3.3-Halo in HCT116 cells with indicated conditions: DMSO (*N* = 20 cells), 5Ph-IAA 1 h (*N* = 20 cells), 5Ph-IAA 3 h (*N* = 20 cells), 5Ph-IAA 6 h (*N* = 20 cells). Note that the MSD curves for the three 5Ph-IAA time points largely overlap. **l**, Two-point MSD plots (*d* < 150 nm; ± SD among cells) of H3.3-Halo in HCT116 cells with indicated conditions: DMSO (black, *N* = 20 cells), 5Ph-IAA (ΔRAD21, red, *N* = 20 cells). **m**, Two-point MSD plots (*d* < 150 nm; ± SD among cells) of H2B-Halo in HCT116 cells with indicated conditions: DMSO (black, *N* = 20 cells), 5Ph-IAA (ΔRAD21, red, *N* = 20 cells). ***, P < 0.001 by the two-sided Kolmogorov-Smirnov test for DMSO vs 5Ph-IAA (P = $2.70 \times 10^{-4}$).

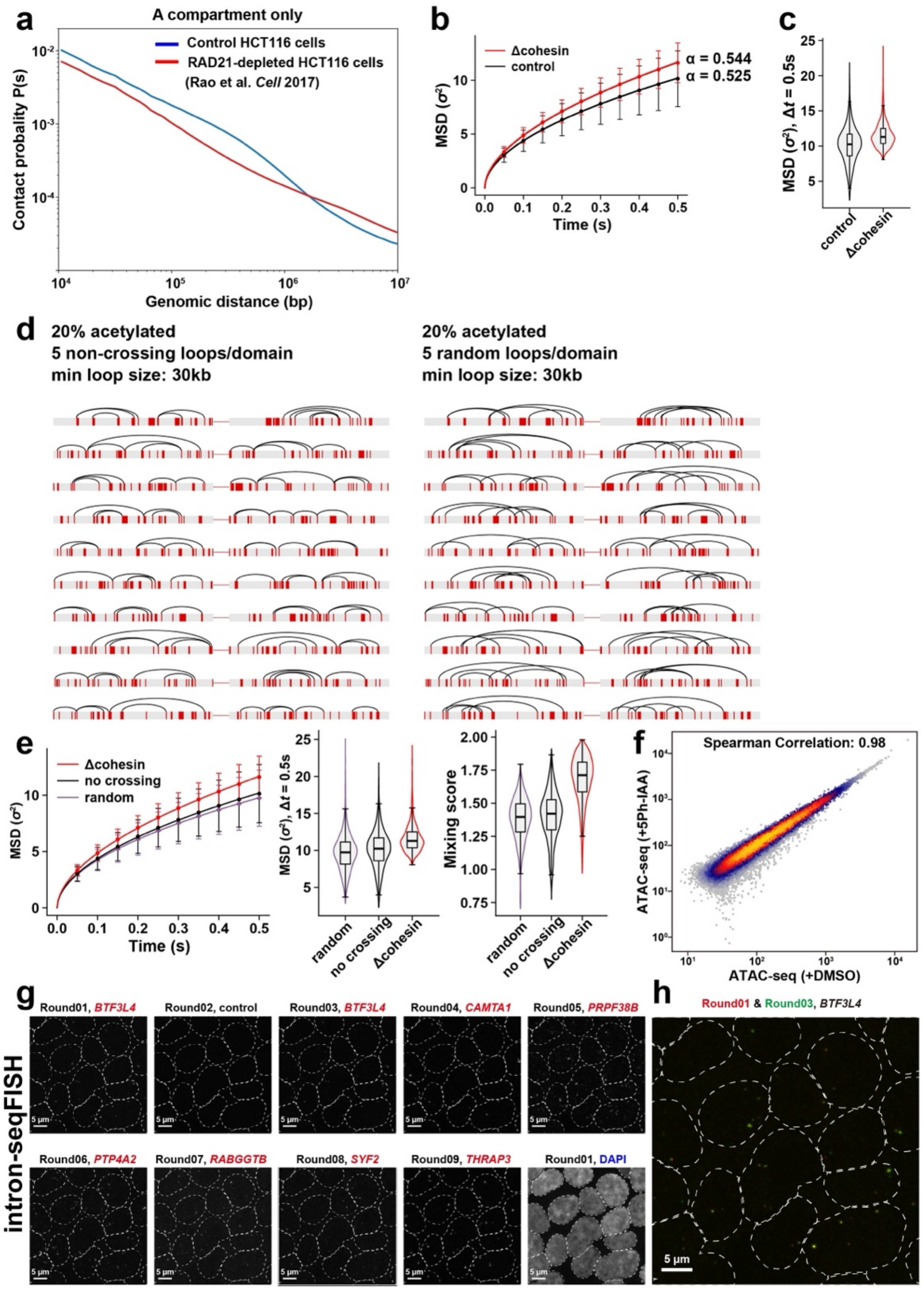


**Fig. S10: Hi-C and polymer simulation suggest local domain mixing by cohesin depletion.**
**a**, Hi-C contact probability $P(s)$ only for the A compartment region in HCT116 cells, with cohesin (control, blue) and after RAD21 depletion (red), were plotted from the Hi-C data of Rao

et al. (Cell, 2017). Upon RAD21 depletion, short-range (up to ~1.5 Mb) contacts decreased, whereas long-range (several-Mb scale) contacts increased, consistent with local mixing of chromatin domains at multi-megabase scales after cohesin depletion. **b**, The temporal variation of MSD of simulated 1-kb beads plotted by showing the average and the standard deviation. The average MSD can be fitted by $14.7\sigma^2\Delta t^{0.525}$ (control) and $17.0\sigma^2\Delta t^{0.544}$ (Δcohesin) with $\sigma$ being the length unit in the simulations. **c**, Distributions of MSD of simulated 1-kb beads at $\Delta t = 0.5$ s. **d,** Chains having different acetylation and loop patterns used in simulations. The acetylated regions (red) of fraction $\varphi = 20\%$ were randomly generated, and five loops connecting 10 sites in the acetylated regions were created under the condition of the minimum loop length of 30 kb. The case that loops do not cross each other (left), and the case that loops are permitted to cross (right). **e,** The simulated chromatin movement. The temporal variation of MSD of 1-kb beads (left), the distributions of MSD of 1-kb beads at $\Delta t = 0.5$ s (middle), and the distributions of the mixing score of two consecutive domains (right). The control model in the main text and the panels **b** and **c** in the present figure corresponds to the no-crossing model in the panel **e**. The average temporal variation of MSD can be fitted by $\sim\Delta t^{\alpha}$ with $\alpha = 0.544$ for the Δcohesin case, while $\alpha = 0.518$ for chains allowing random crossing and $\alpha = 0.525$ for chains without crossing in the control case with cohesin. The average MSD at $\Delta t = 0.5$ s in the Δcohesin case is 1.19 times larger than that in the control case for chains allowing random crossing, and 1.15 times larger for chains without crossing. The average mixing score in the Δcohesin case is 1.22 times larger than that in the control case for chains allowing random crossing, and 1.20 times larger for chains without crossing. **f**, Genome-wide comparison of ATAC-seq libraries prepared from HCT116; RAD21-mAC treated with DMSO or 5Ph-IAA. A Pearson correlation coefficient was calculated by genome-wide correlation analysis. **g**, Representative maximum-intensity-projection images obtained by intron-seqFISH in fixed HCT116; RAD21-mAC cells. In each hybridization round, nascent intronic RNA from a single target gene was visualized. The gene targeted in each round is indicated above. Dashed lines indicate nuclear boundaries determined from DAPI images. **h**, Overlay of signals detecting the same gene in two independent rounds (Round 01 and 03). The strong overlap demonstrates accurate round-to-round alignment and high reproducibility of seqFISH detection.

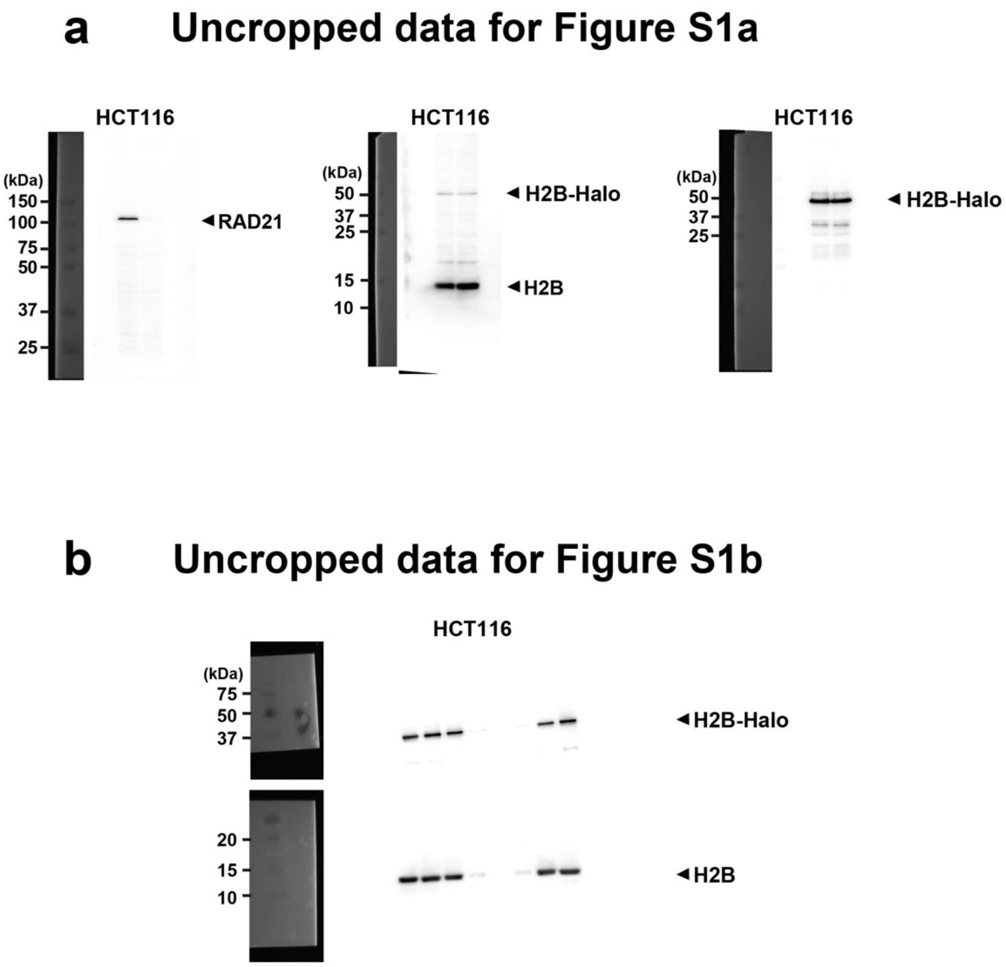


**Fig. S11: Uncropped western blot images.**

**a**, Uncropped western blot images for Fig. S1a. **b**, Uncropped western blot images for Fig. S1b.

### Uncropped data for Figure 3

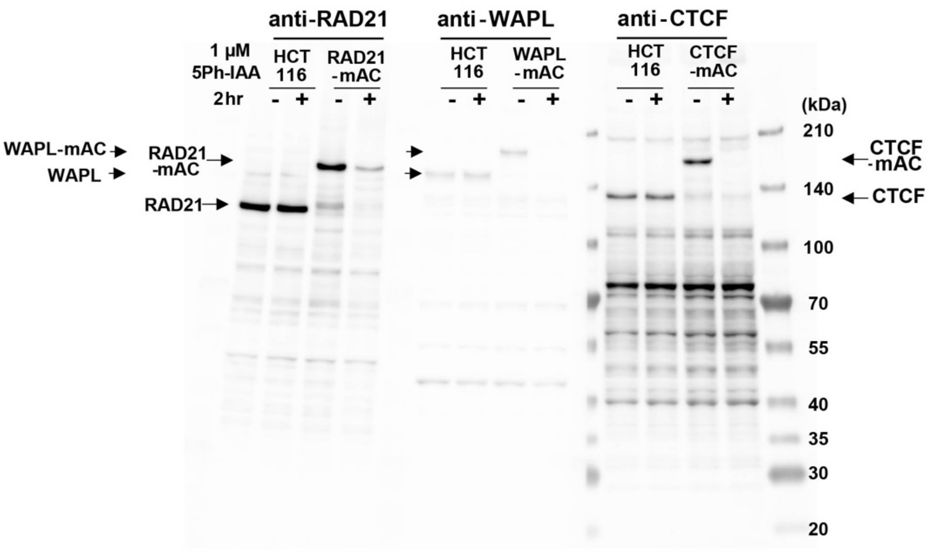


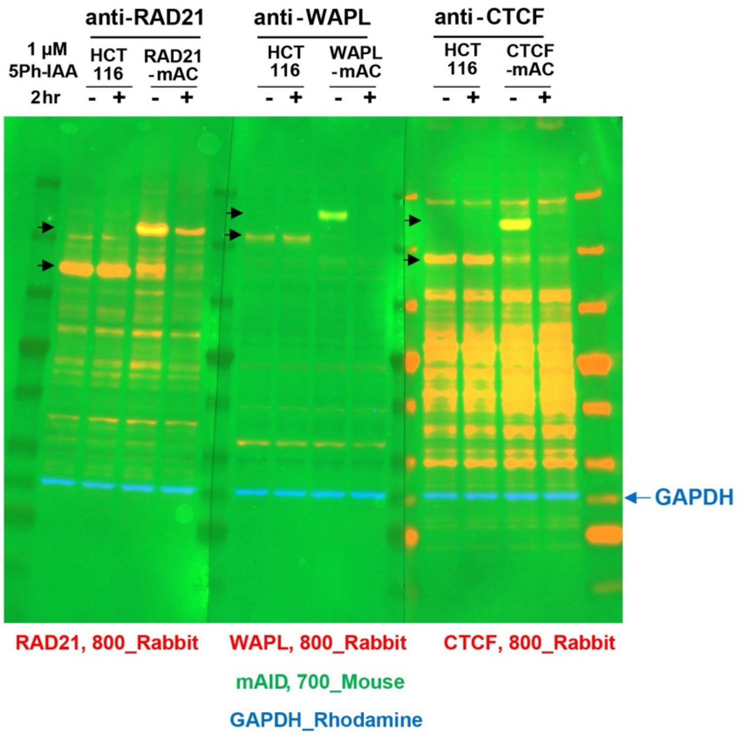


**Fig. S12: Uncropped western blot images.**

**a**, Uncropped western blot images for Fig. 3. **b**, Uncropped fluorescent western blot images for Fig. 3.

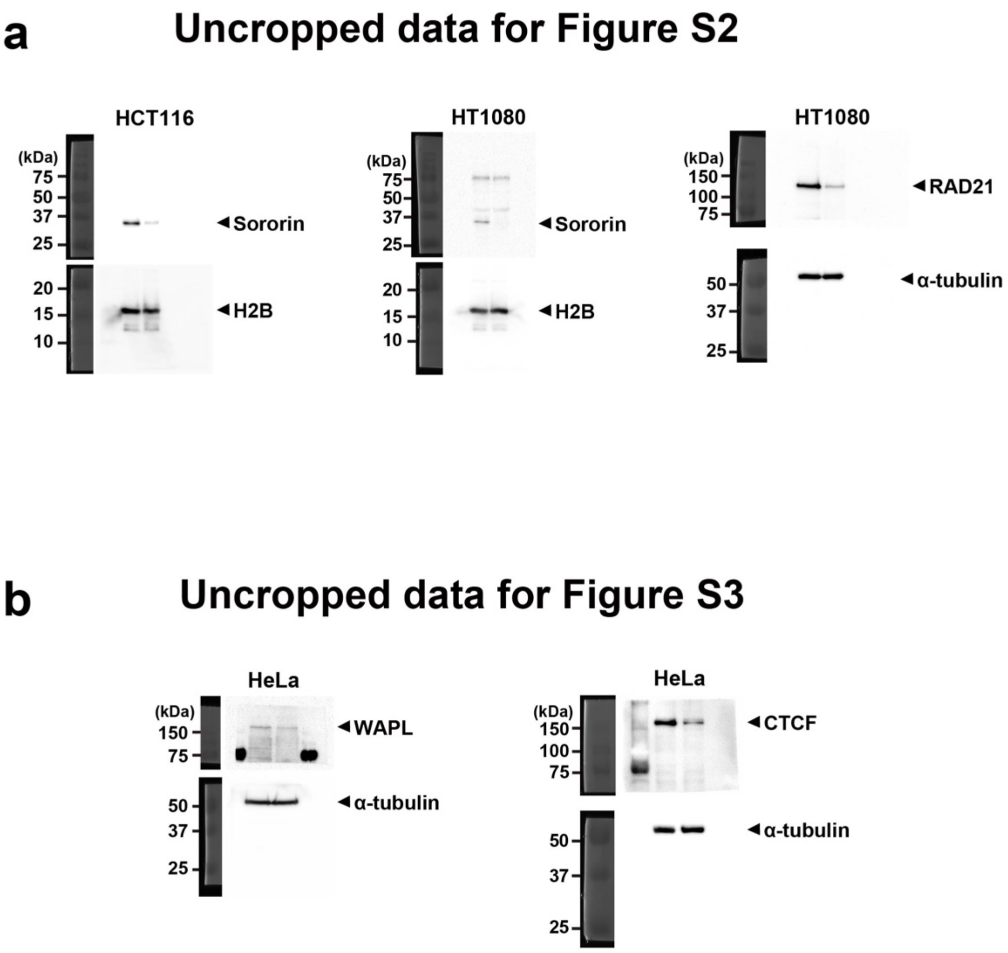


**Fig. S13: Uncropped western blot images.**

**a**, Uncropped western blot images for Fig. S2. **b**, Uncropped western blot images for Fig. S3.

**a** Uncropped data for Figure S5a

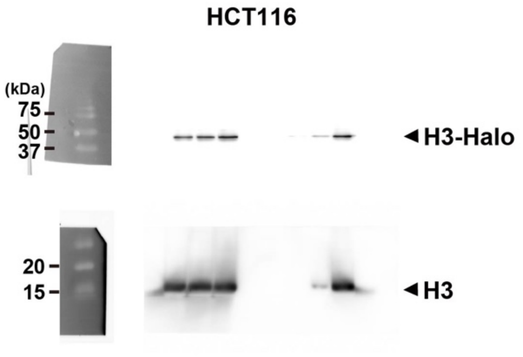


**b** Uncropped data for Figure S5b

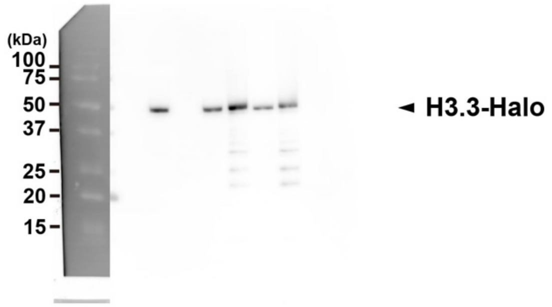


**c** Uncropped data for Figure S5c

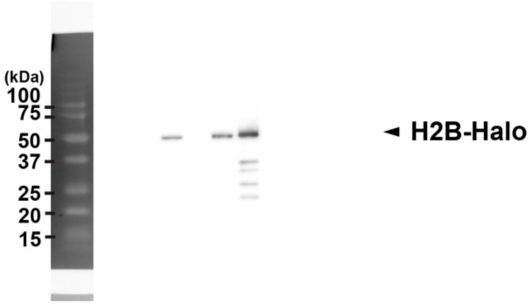


**Fig. S14: Uncropped western blot images.**

**a**, Uncropped western blot images for Fig. S5a. **b**, Uncropped western blot images for Fig. S5b. **c**, Uncropped western blot images for Fig. S5c.

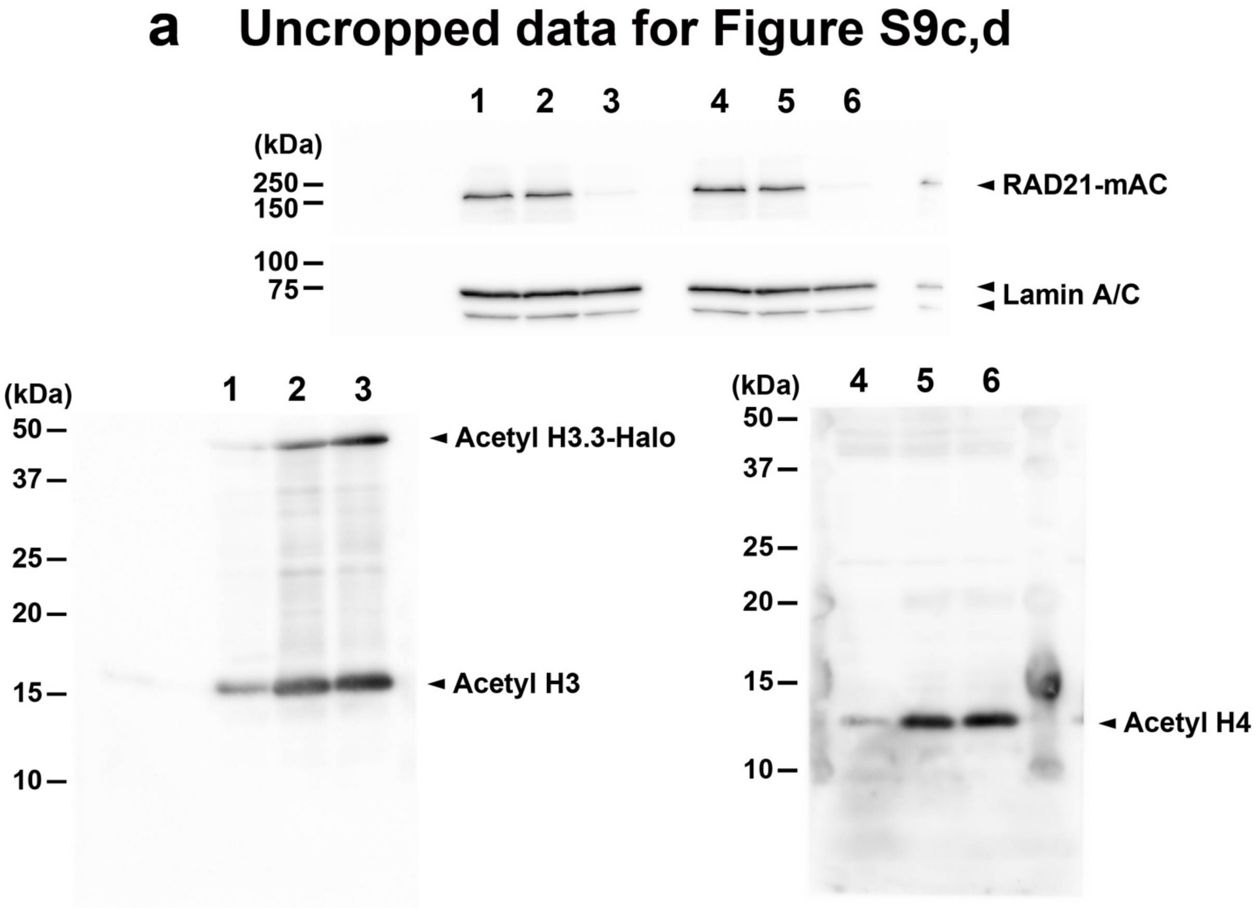


**Fig. S15: Uncropped western blot images.**

**a**, Uncropped western blot images for Fig. S9c,d.

#### Movie legends

**Movie S1.**
Movie data (50 ms/frame) of single H2B-Halo nucleosomes labeled with TMR in a living HCT116 cell recorded with an sCMOS ORCA-Fusion BT camera (Hamamatsu Photonics). Clear, well-separated dots exhibiting single-step photobleaching after background subtraction (see Fig. 1c) indicate that each dot corresponds to a single H2B–Halo–TMR molecule within a nucleosome. Circles in the right movie highlight tracked dots. Dots observed for more than three frames were used for further analysis.

**Movie S2.**
Movie data (50 ms/frame) of single H2B-Halo nucleosomes labeled with TMR in a living HCT116 cell treated with 0.1% DMSO (Control; left) or 1 μM 5Ph-IAA (ΔRAD21; right). Note that nucleosomes move faster when treated with 5Ph-IAA than with DMSO.

**Movie S3.**
Movie data (50 ms/frame) of single H2B-Halo nucleosomes labeled with TMR in a living HCT116 cell. Each moving dot is outlined by a colored circle indicating its RL-classified subpopulation: orange, Superfast; yellow, Fast; green, Slow; blue, Super-slow. Some dots remain unclassified, as the RL algorithm classifies only trajectories tracked for more than 11 consecutive frames (0.5 s). Note that Superfast does not correspond to free histones. The nuclear region is outlined by a dotted yellow line.

**Movie S4.**
3D-structured illumination microscopy (3D-SIM) volume reconstruction of a fixed HCT116 nucleus labeled with DAPI and H3.3–Halo–TMR. Images were acquired with a Nikon N-SIM-S system using a piezo Z-drive at 0.12 μm step size. Images were reconstructed with stack reconstruction in NIS-Elements. The movie was generated with the NIS-Elements Volume Viewer (mode: MAX-IP).

**Movie S5.**
3D-SIM volume reconstruction of a live HCT116 nucleus labeled with H3.3–Halo–TMR.

**Movie S6.**
3D-SIM volume reconstruction of a fixed HCT116 nucleus treated with 0.1% DMSO (Control) and labeled with DAPI and H3.3–Halo–TMR.

**Movie S7.**
3D-SIM volume reconstruction of a fixed HCT116 nucleus labeled with H3.3–Halo–TMR. The cell was treated with 1 μM 5Ph-IAA (ΔRAD21).

**Movie S8.**
Video of two closely positioned single nucleosomes (left, labeled with PA-JF549; right, labeled with PA-JF646). The nucleosomes were recorded simultaneously at 50 ms/frame using W-VIEW GEMINI (Hamamatsu Photonics) with a pixel size of 65 nm. Tracked dots are outlined with magenta circles. An example of two dots that coexisted in the same frame and, by chance, had an

average inter-point distance of <150 nm is highlighted by a square and enlarged in the inset. Trajectories of these two dots are shown in red. The movie was generated using the TrackMate plugin.

**Movie S9.**
Simulated dynamics of an example chromatin chain consisting of two consecutive domains (green and grey) with cohesin (left, control) and without cohesin (right, Δcohesin) (see Fig. 8b-c).
